## Supplemental Data for "Single amino-acid mutation in a *Drosophila melanogaster* ribosomal protein: an insight in uL11 transcriptional activity"

**Supplementary Figure 1: Specificity of the anti-uL11K3me3 antibody**

0.2 (left) and 0.05 µg (right) of each peptide were deposited on a nitrocellulose membrane. Membranes were then incubated with the indicated primary antibodies. Secondary antibodies were as described in Materials and Methods.

**Peptides:** unmethylated uL11, uL11K10me3, uL11K3A, uL11K3me2, and uL11K3me3 peptides were synthesized at the proteomic platform of the Institute of Biology Paris Seine; H3K4me3 and H3K9me3 peptides were from Diagenode, C16000003 and C160000056, respectively.

**Antibodies:** PI: rabbit preimmun serum; $\alpha$-uL11: 1/14000, described in Materials and Methods; $\alpha$-uL11K3me3: 1/10000, described in Materials and Methods; $\alpha$-H3K4me3: 1/1000, Diagenode C15310003; $\alpha$-H3K9me3: 1/1000, Diagenode C15100146. Secondary antibodies: 1/10000.

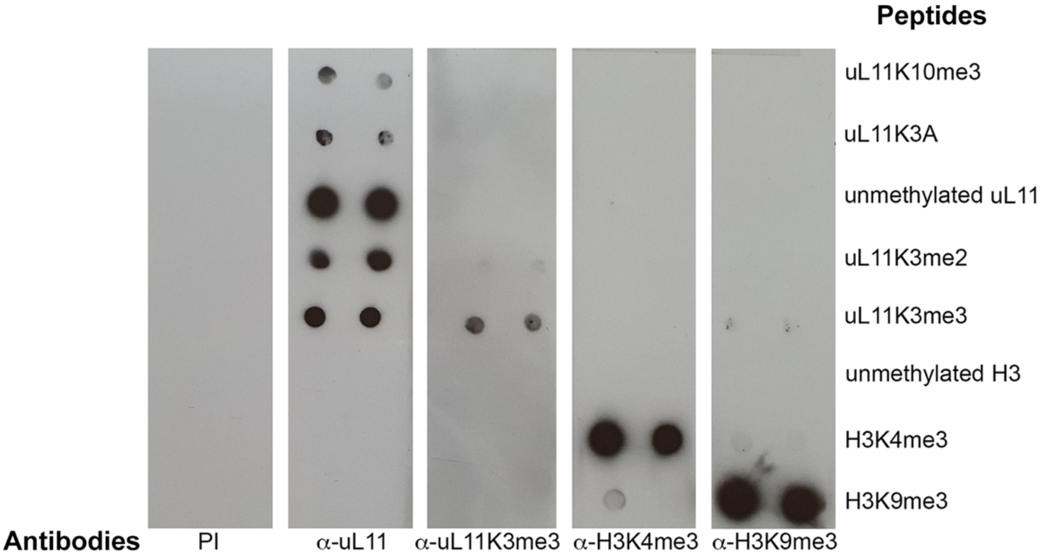

**Supplementary Figure 2: Molecular screening for the *uL11*$^{K3A}$ allele**

**A –** Rationale for discriminative PCR. Purple bases correspond to the target codon. Red bases stand for locked nucleic acids (LNA). The *LNAWT* primer ended with the lysine AAA codon of the wild-type *uL11* gene whereas the *LNAK3A* primer ended with the alanine GCC codon corresponding to the desired mutation.

**B –** qPCRs were performed with the *LNAK3A* primer matching the *uL11*$^{K3A}$ allele. Red curve: plasmid carrying the *uL11*$^{K3A}$ allele as positive control. Black curve: genomic DNA from a wild-type fly. Blue curves: pools of up to 5 different genomic DNAs from candidate G1 flies considered to be positive. Green curves: pools of up to 5 different genomic DNAs from candidate G1 flies considered to be negative.

**C –** The same qPCRs were performed on individual genomic DNAs from the pools that were previously found to be positive for the *uL11*$^{K3A}$ allele. Several individuals wearing the mutation were thus identified (blue curves).

**D –** High Resolution Melting Analysis (HMRA) of *uL11* mutants. Melting profile of the *uL11* amplicons from genomic DNAs of G1 flies. Melting peaks flatter and broader than the reference (black) revealed the presence of two different amplicons, indicating that the tested DNA contained a mutation at the *uL11* locus. Melting curves were normalized according to the method described by (26). RFU: Relative Fluorescence Unit.

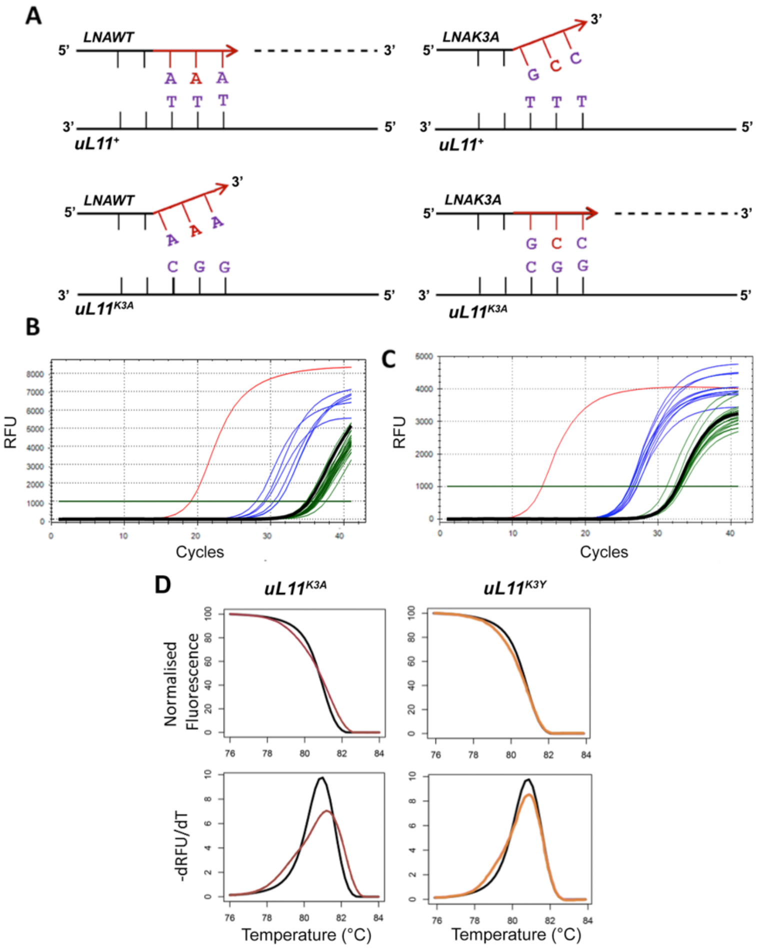

**Supplementary Figure 3: Sequence of the *uL11* alleles of G0 flies**

Founder G0 flies were named after their gender (M, male; F, female) and the order of their emergence. Each allele was recovered in several descendants of the same founders. The *uL11*$^{K3A}$ and *uL11*$^{\Delta K3}$ alleles were found in the progeny of two different founders. Substitution alleles were named to reflect the amino acid change in the uL11 protein, following the amino acid one letter code. The bottom two alleles were named after the reading frameshift they introduce in the *uL11* gene. The wild-type *uL11* sequence is provided as reference. The start and the lysine 3 codons of *uL11* are highlighted in grey.

Mutants F-4 and F+2 introduce a +2 reading frame shift that puts the uL11 CDS in frame with an ATG codon located in the 5'UTR. A protein with a 24 amino acid extension might then be produced.

| Founder | Allele | Sequence |
|---|---|---|
| | wild-type | ACCGCTATGCCTCCCAAATTCGACCCAACGGAA |
| M-12 | K3A-12 | ACCGCTATGCCTCCCGCCTTCGACCCAACGGAA |
| M-43 | K3A-43 | ACCGCTATGCCTCCCGCCTTCGACCCAACGGAA |
| F-6 | K3A-6 | ACCGCTATGCCTCCCGCCTTCGACCCAACGGAA |
| M-12 | ΔK3-12 | ACCGCTATGCCTCCC---TTCGACCCAACGGAA |
| M-43 | ΔK3-43 | ACCGCTATGCCTCCC---TTCGACCCAACGGAA |
| M-31 | K3Y | ACCGCTATGCCTCCCTACTTCGACCCAACGGAA |
| M-5 | ΔK3F4 | ACCGCTATGCCTCCC------GACCCAACGGAA |
| M-5 | P2QK3R | ACCGCTATGCCTCAACGCTTCGACCCAACGGAA |
| M-12 | P2LK3E | ACCGCTATGCCTCTTGAATTCGACCCAACGGAA |
| M-43 | F-4 | ACCGCTATGCCTCC----TTCGACCCAACGGAA |
| F-6 | F+2 | ACCGCTATGCCTCCCTATGCTTCGACCCAACGGAA |

**Supplementary Figure 4: Analysis of bristles and wings in female *uL11* mutants**

**A –** Length of anterior scutellar bristles of wild-type females (blue; n = 54), $uL11^{K3A}/uL11^{+}$ (burgundy, n = 33) and $uL11^{K3Y}/uL11^{K3Y}$ (orange, n = 25).

**B –** Length of posterior scutellar bristles of wild-type males (blue; n = 51), $uL11^{K3A}/uL11^{+}$ (burgundy, n = 36) and ($uL11^{K3Y}/uL11^{K3Y}$ (orange, n = 50).

**C –** Wing size of *uL11* wild-type females (blue; n = 29), $uL11^{K3A}/uL11^{+}$ (burgundy, n = 30) and $uL11^{K3Y}/uL11^{K3Y}$ (orange, n = 25).

t-tests: *** p-value < 0.001; ** p-value < 0.01; * p-value < 0.05; ns: non significant.

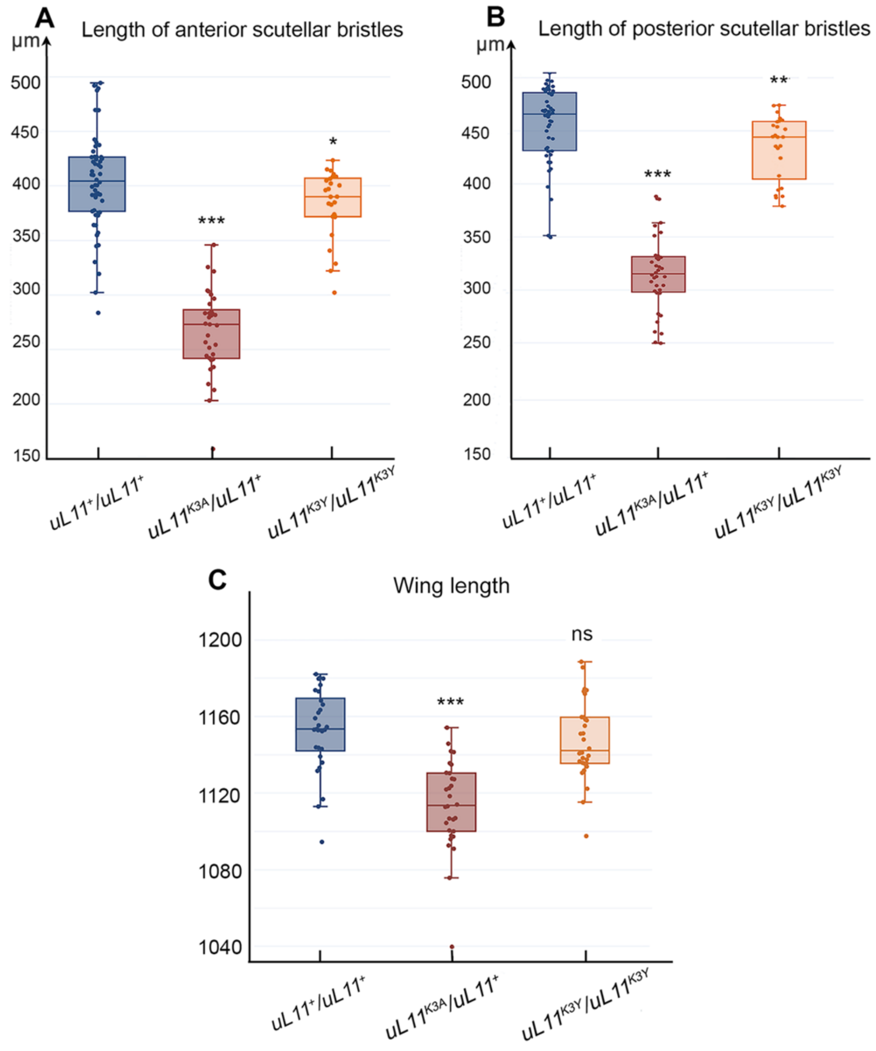

**Supplementary Table 1: Oligonucleotides used in this study.**
In primers *LNA-WT* and *LNA-K3A*, uppercase nucleotides correspond to the LNA bases. In primers *pho-sgRNA_F* and *pho-sgRNA_R*, uppercase nucleotides correspond to the floating sequences used for cloning. The bold guanosine was introduced to increase efficiency of the U6 promoter. In the *ssODN*, the complementary ATG and alanine codon sequences are bold and in uppercases, the PAM sequence corresponding to the single guide RNA is bold.

| | |
|---|---|
| ***High Resolution Melting Analysis*** | |
| *uL11-HRMA_F* | 5'-tgcggtaaagtacatgagctg-3' |
| *uL11-HRMA_R* | 5'-tcgaagctcaactcctcaca-3' |
| ***LNA Primers*** | |
| *LNA-WT* | 5'-gataccgctatgcctcccaAa-3' |
| *LNA-K3A* | 5'-gataccgctatgcctcccgCc-3' |
| *CRISPR1_R* | 5'-gaccgaggggaccgatctt-3' |
| ***uL11 Sequencing*** | |
| *uL11-708_F* | 5'-cgctactgagctttgctacacccc-3' |
| *uL11+805_R* | 5'-caataacatcgtgagggtgct-3' |
| ***Guide RNA*** | |
| *sgRNA* | 5'-tccgttgggtcgaatttggg-3' |
| ***Cloning of sgRNA in pU6-BbsI-chiRNA*** | |
| *pho-sgRNA_F* | 5'-CTTC**G**tccgttgggtcgaatttggg-3' |
| *pho-sgRNA_R* | 5'-AAACcccaaattcgacccaacggac-3' |
| ***Template for fly mutagenesis*** | |
| *ssODN* | 5'-agctcaactcctcacaaaaacactcgcttacttacccaatttaacttccgttgggtcgaa**GGC**ggg**aggCAT**a gcggtatcttggcttgaacagtcgctgtaaggcaaagattacgttagttt-3' |
| ***uL11$^{K3Y}$ directed mutagenesis*** | |
| *K3Y_F* | 5'-caccatgcctccctacttcgacccaacgg-3' |
| *K3Y_R* | 5'-ccgttgggtcgaagtagggaggcatggtg-3' |

**Supplementary Table 2: RNA-seq of wing imaginal discs (GEO accession number GSE181926).**

| Sample name | Genotype | Total reads | Aligned reads | Unmap reads | Reads with multiple alignment | Reads used for analysis |
|---|---|---|---|---|---|---|
| wc_1 | $w^{1118}$ ; $corto^{+}$ | 1.50E+07 | 6.70E+06 | 9.00E+05 | 2.20E+06 | 6.71E+06 |
| wc_2 | $w^{1118}$ ; $corto^{+}$ | 1.50E+07 | 6.40E+06 | 1.20E+06 | 2.20E+06 | 6.44E+06 |
| cortoL1420_1 | $w^{1118}$ ; $corto^{L1}/corto^{420}$ | 2.71E+07 | 1.18E+07 | 1.20E+06 | 1.20E+07 | 1.18E+07 |
| cortoL1420_2 | $w^{1118}$ ; $corto^{L1}/corto^{420}$ | 2.71E+07 | 1.14E+07 | 1.90E+06 | 1.20E+07 | 1.14E+07 |
| w_1 | $w^{1118}$ ; $uL11^{+}$ | 2.58E+07 | 3.88E+07 | 5.39E+05 | 1.71E+07 | 2.08E+07 |
| w_2 | $w^{1118}$ ; $uL11^{+}$ | 3.22E+07 | 4.03E+07 | 3.50E+05 | 1.17E+07 | 2.79E+07 |
| w_3 | $w^{1118}$ ; $uL11^{+}$ | 3.49E+07 | 4.87E+07 | 6.90E+05 | 1.87E+07 | 2.86E+07 |
| K3A_1 | $w^{1118}$ ; $uL11^{K3A}$ | 2.53E+07 | 3.34E+07 | 1.62E+06 | 1.14E+07 | 2.00E+07 |
| K3A_2 | $w^{1118}$ ; $uL11^{K3A}$ | 4.01E+07 | 1.54E+07 | 4.96E+06 | 1.38E+08 | 1.06E+07 |
| K3A_3 | $w^{1118}$ ; $uL11^{K3A}$ | 3.28E+07 | 8.30E+07 | 2.93E+06 | 6.18E+07 | 1.97E+07 |
| K3Y_1 | $w^{1118}$ ; $uL11^{K3Y}$ | 3.07E+07 | 3.87E+07 | 4.65E+05 | 1.14E+07 | 2.64E+07 |
| K3Y_2 | $w^{1118}$ ; $uL11^{K3Y}$ | 3.16E+07 | 4.32E+07 | 1.17E+06 | 1.58E+07 | 2.59E+07 |
| K3Y_3 | $w^{1118}$ ; $uL11^{K3Y}$ | 3.34E+07 | 3.88E+07 | 4.45E+07 | 8.65E+06 | 2.92E+07 |

**Supplementary Table 3: Genes deregulated in at least one of the three genotypes (*corto$^{L1}$/corto$^{420}$*, *uL11$^{K3A}$*, *uL11$^{K3Y}$*).**

Green: Up-regulated genes, $log_2$ fold-change > 0.5, adjusted p-value < 5.E-02

Orange: Down-regulated genes, $log_2$ fold-change < -0.5, adjusted p-value < 5.E-02

Blue: Genes up-regulated in other *RPG* mutants (35, 36)

| Gene ID | Gene symbol | Base Mean | $log_2$FoldChange *corto$^{L1}$/corto$^{420}$* versus *w$^{1118}$* | adj p-value *corto$^{L1}$/corto$^{420}$* versus *w$^{1118}$* | $log_2$FoldChange *uL11$^{K3A}$* versus *w$^{1118}$* | adj p-value *uL11$^{K3A}$* versus *w$^{1118}$* | $log_2$FoldChange *uL11$^{K3Y}$* versus *w$^{1118}$* | adj p-value *uL11$^{K3Y}$* versus *w$^{1118}$* |
|---|---|---|---|---|---|---|---|---|
| FBgn0085813 | 18SrRNA-Psi:CR41602 | 492 | 0.12 | 5.47E-01 | 2.53 | 1.23E-03 | -0.01 | 1.00E+00 |
| FBgn0085753 | 28SrRNA-Psi:CR40596 | 815 | 0.17 | 2.96E-01 | 1.41 | 1.30E-02 | -0.01 | 1.00E+00 |
| FBgn0085771 | 28SrRNA-Psi:CR40741 | 96 | 3.34 | 8.74E-05 | 3.25 | 5.48E-05 | -0.01 | 1.00E+00 |
| FBgn0267508 | 28SrRNA-Psi:CR45848 | 67 | 0.18 | 3.14E-01 | 2.82 | 4.86E-04 | 0.00 | 1.00E+00 |
| FBgn0267511 | 28SrRNA-Psi:CR45851 | 67 | -0.02 | 8.65E-01 | 1.85 | 1.04E-02 | 0.00 | 1.00E+00 |
| FBgn0267513 | 28SrRNA-Psi:CR45853 | 36 | 0.04 | 8.53E-01 | 2.51 | 1.39E-03 | -0.01 | 1.00E+00 |
| FBgn0267515 | 28SrRNA-Psi:CR45855 | 55 | 0.08 | 5.85E-01 | 1.60 | 9.23E-03 | 0.00 | 1.00E+00 |
| FBgn0267520 | 28SrRNA-Psi:CR45860 | 204 | 0.27 | 8.96E-02 | 1.47 | 1.26E-02 | 0.00 | 1.00E+00 |
| FBgn0052016 | 4E-T | 2991 | -0.97 | 6.64E-13 | -0.07 | 3.82E-01 | 0.00 | 1.00E+00 |
| FBgn0023129 | aay | 91 | -0.12 | 6.58E-01 | -1.00 | 6.49E-03 | -0.18 | 1.45E-01 |
| FBgn0039890 | ABCD | 2965 | -0.64 | 4.77E-10 | -0.08 | 2.74E-01 | -0.06 | 2.18E-01 |
| FBgn0004852 | Ac76E | 330 | -3.26 | 6.23E-33 | 0.04 | 6.42E-01 | 0.00 | 1.00E+00 |
| FBgn0027620 | Acf | 2690 | -0.62 | 3.33E-06 | 0.02 | 8.64E-01 | 0.01 | 9.06E-01 |
| FBgn0051865 | Ada1-1 | 19 | 0.05 | 8.42E-01 | -5.48 | 5.96E-05 | 0.07 | 3.78E-01 |
| FBgn0026086 | Adar | 2162 | -0.71 | 2.33E-05 | -0.01 | 9.70E-01 | 0.00 | 1.00E+00 |
| FBgn0026573 | ADD1 | 1787 | -0.61 | 3.07E-10 | -0.02 | 9.06E-01 | 0.01 | 1.00E+00 |
| FBgn0052068 | Adi1 | 187 | 0.12 | 6.12E-01 | 0.41 | 5.04E-03 | 0.79 | 4.87E-07 |
| FBgn0038223 | Afti | 793 | -0.64 | 8.75E-06 | 0.00 | 1.00E+00 | 0.00 | 1.00E+00 |
| FBgn0029813 | AgmNAT | 22 | 0.59 | 2.40E-02 | 0.05 | 6.08E-01 | 0.00 | 1.00E+00 |
| FBgn0250816 | AGO3 | 64 | -0.64 | 1.52E-02 | 2.09 | 1.17E-13 | 0.38 | 5.07E-02 |
| FBgn0024912 | agt | 851 | 0.61 | 3.04E-06 | 1.63 | 1.55E-43 | 0.21 | 4.55E-02 |
| FBgn0014455 | Ahcy | 3702 | 0.65 | 5.05E-11 | 0.02 | 8.74E-01 | 0.00 | 1.00E+00 |
| FBgn0033351 | AIMP1 | 804 | 0.72 | 4.73E-05 | 0.05 | 5.43E-01 | 0.00 | 1.00E+00 |
| FBgn0050185 | AIMP3 | 582 | 0.65 | 2.84E-04 | 0.03 | 7.70E-01 | 0.00 | 1.00E+00 |
| FBgn0027932 | Akap200 | 21721 | -1.15 | 5.83E-49 | 0.04 | 6.85E-01 | 0.00 | 1.00E+00 |
| FBgn0012036 | Aldh | 651 | 0.77 | 6.08E-03 | 0.00 | 9.72E-01 | 0.07 | 4.09E-01 |
| FBgn0011297 | Alg3 | 292 | 0.70 | 8.24E-06 | -0.28 | 7.21E-03 | -0.01 | 9.05E-01 |
| FBgn0283479 | Alp1 | 47 | 0.10 | 5.76E-01 | -0.60 | 5.71E-02 | -0.01 | 1.00E+00 |
| FBgn0033423 | Alp6 | 12 | -0.02 | 8.84E-01 | -1.02 | 3.05E-02 | -0.09 | 3.70E-01 |
| FBgn0034710 | Alp7 | 85 | -0.13 | 2.20E-01 | -8.44 | 5.06E-06 | -0.01 | 1.00E+00 |
| FBgn0034712 | Alp8 | 99 | -0.07 | 4.75E-01 | -0.66 | 5.26E-02 | 0.00 | 1.00E+00 |
| FBgn0086361 | alph | 3610 | -0.65 | 1.44E-17 | 0.04 | 6.45E-01 | 0.00 | 1.00E+00 |
| FBgn0015571 | alpha-Est3 | 211 | 0.63 | 2.46E-03 | 0.42 | 6.49E-03 | 0.00 | 1.00E+00 |
| FBgn0003884 | alphaTub84B | 98958 | 0.58 | 2.30E-12 | -0.03 | 7.61E-01 | -0.03 | 4.82E-01 |
| FBgn0003886 | alphaTub85E | 97 | 3.34 | 1.93E-15 | 0.01 | 8.95E-01 | 0.01 | 9.37E-01 |
| FBgn0031068 | Alr | 351 | 0.61 | 1.35E-03 | 0.39 | 6.66E-03 | 0.02 | 8.04E-01 |
| FBgn0000075 | amd | 647 | 0.26 | 1.88E-01 | 1.20 | 1.13E-18 | 0.00 | 1.00E+00 |
| FBgn0025686 | Amnionless | 12 | -0.02 | 9.37E-01 | -0.61 | 5.12E-02 | -0.01 | 1.00E+00 |
| FBgn0030328 | Amun | 7039 | -0.79 | 7.59E-17 | 0.01 | 9.64E-01 | 0.00 | 1.00E+00 |
| FBgn0012037 | Ance | 10590 | -0.08 | 5.00E-01 | -0.70 | 3.42E-22 | -0.03 | 4.37E-01 |
| FBgn0032535 | Ance-2 | 17 | -0.11 | 5.24E-01 | -0.59 | 4.28E-02 | 0.00 | 1.00E+00 |
| FBgn0032536 | Ance-3 | 9 | 0.02 | 8.74E-01 | 1.73 | 1.06E-02 | 0.01 | 1.00E+00 |
| FBgn0011747 | Ank | 7198 | -0.65 | 1.55E-07 | -0.04 | 6.39E-01 | 0.00 | 1.00E+00 |
| FBgn0052000 | anne | 6510 | -0.58 | 4.18E-05 | 0.01 | 9.54E-01 | 0.01 | 1.00E+00 |
| FBgn0260642 | Antp | 1348 | -0.77 | 7.50E-06 | -0.02 | 8.95E-01 | 0.01 | 1.00E+00 |
| FBgn0267408 | AOX1 | 329 | 0.55 | 2.96E-02 | 3.81 | 3.49E-23 | 0.01 | 1.00E+00 |
| FBgn0036111 | Aps | 7280 | -0.69 | 1.35E-08 | 0.02 | 8.52E-01 | 0.00 | 1.00E+00 |
| FBgn0286516 | aqz | 7288 | -1.14 | 3.19E-11 | -0.03 | 8.04E-01 | 0.00 | 1.00E+00 |
| FBgn0033928 | Arc2 | 184 | 0.03 | 9.37E-01 | 3.19 | 8.75E-82 | 0.22 | 8.41E-02 |
| FBgn0038893 | Archease | 334 | 0.25 | 2.21E-01 | 0.63 | 7.35E-06 | 0.00 | 1.00E+00 |
| FBgn0013749 | Arf102F | 1709 | -0.65 | 3.80E-07 | -0.06 | 5.12E-01 | -0.01 | 1.00E+00 |
| FBgn0017418 | ari-1 | 1097 | -0.60 | 1.00E-06 | 0.04 | 6.78E-01 | 0.00 | 1.00E+00 |
| FBgn0041164 | armi | 2051 | -0.65 | 2.47E-07 | -0.05 | 6.22E-01 | 0.00 | 1.00E+00 |
| FBgn0031050 | Arp10 | 640 | 0.68 | 3.14E-04 | 0.00 | 9.77E-01 | 0.00 | 1.00E+00 |
| FBgn0001961 | Arpc1 | 2195 | 0.59 | 5.56E-08 | 0.02 | 8.26E-01 | -0.01 | 1.00E+00 |
| FBgn0284255 | Arpc4 | 1043 | 1.13 | 5.45E-17 | 0.08 | 3.28E-01 | 0.00 | 1.00E+00 |
| FBgn0031437 | Arpc5 | 1308 | 0.77 | 7.24E-08 | 0.00 | 9.90E-01 | -0.01 | 9.73E-01 |
| FBgn0033062 | Ars2 | 4686 | -0.64 | 7.16E-18 | -0.01 | 9.72E-01 | 0.00 | 1.00E+00 |
| FBgn0039908 | Asator | 1415 | -0.76 | 7.99E-10 | -0.04 | 7.13E-01 | 0.00 | 1.00E+00 |
| FBgn0270926 | AsnS | 394 | 1.20 | 2.78E-06 | 0.16 | 1.49E-01 | 0.17 | 1.29E-01 |
| FBgn0034793 | asrij | 688 | 0.86 | 1.78E-07 | 0.36 | 3.09E-03 | 0.00 | 1.00E+00 |
| FBgn0017424 | asRNA:CR11538 | 422 | 0.58 | 1.49E-02 | -0.04 | 7.33E-01 | 0.01 | 1.00E+00 |
| FBgn0265295 | asRNA:CR42871 | 56 | -0.73 | 4.32E-03 | 0.11 | 3.22E-01 | 0.01 | 9.91E-01 |
| FBgn0263344 | asRNA:CR43425 | 78 | -0.50 | 3.31E-02 | 0.80 | 9.37E-03 | 0.00 | 1.00E+00 |
| FBgn0263345 | asRNA:CR43426 | 51 | 0.67 | 1.10E-02 | 0.01 | 9.32E-01 | -0.01 | 1.00E+00 |
| FBgn0263445 | asRNA:CR43468 | 99 | -0.59 | 7.51E-03 | 0.27 | 4.57E-02 | 0.02 | 7.54E-01 |
| FBgn0264822 | asRNA:CR44030 | 121 | -0.10 | 7.14E-01 | 0.86 | 2.95E-04 | 1.64 | 7.16E-13 |
| FBgn0264823 | asRNA:CR44031 | 97 | 0.54 | 1.87E-02 | 1.28 | 2.34E-03 | 0.02 | 7.92E-01 |

| | | | | | | | | |
|---|---|---|---|---|---|---|---|---|
| FBgn0264837 | asRNA:CR44045 | 82 | -0.61 | 1.50E-02 | -0.01 | 9.70E-01 | 0.01 | 1.00E+00 |
| FBgn0265499 | asRNA:CR44368 | 337 | 0.00 | 9.87E-01 | 1.03 | 2.46E-15 | -0.60 | 2.20E-04 |
| FBgn0265613 | asRNA:CR44431 | 123 | 0.83 | 5.37E-04 | -0.03 | 7.85E-01 | 0.01 | 1.00E+00 |
| FBgn0266619 | asRNA:CR45126 | 267 | -0.96 | 7.58E-07 | 2.40 | 2.95E-15 | 0.95 | 3.69E-03 |
| FBgn0266633 | asRNA:CR45140 | 31 | 0.34 | 1.70E-01 | 4.88 | 2.30E-13 | -0.02 | 8.78E-01 |
| FBgn0266681 | asRNA:CR45171 | 27 | -1.00 | 3.76E-03 | 0.34 | 9.99E-02 | 0.05 | 5.27E-01 |
| FBgn0266692 | asRNA:CR45182 | 29 | -0.64 | 1.70E-02 | -0.03 | 7.96E-01 | 0.01 | 1.00E+00 |
| FBgn0266705 | asRNA:CR45195 | 68 | 0.10 | 7.19E-01 | 7.75 | 1.00E-27 | 0.00 | 1.00E+00 |
| FBgn0267022 | asRNA:CR45466 | 32 | -0.97 | 1.72E-03 | 0.00 | 9.72E-01 | 0.00 | 1.00E+00 |
| FBgn0267160 | asRNA:CR45600 | 111 | -5.01 | 2.08E-12 | 5.59 | 1.04E-17 | 5.92 | 1.38E-19 |
| FBgn0267223 | asRNA:CR45663 | 34 | 1.49 | 3.22E-04 | 0.02 | 9.09E-01 | 0.00 | 1.00E+00 |
| FBgn0267302 | asRNA:CR45738 | 32 | 0.61 | 2.24E-02 | -0.08 | 4.91E-01 | -0.01 | 1.00E+00 |
| FBgn0267439 | asRNA:CR45789 | 35 | -0.71 | 1.03E-02 | -0.03 | 7.98E-01 | 0.00 | 1.00E+00 |
| FBgn0267485 | asRNA:CR45835 | 898 | 0.61 | 4.33E-07 | -0.03 | 7.65E-01 | 0.00 | 1.00E+00 |
| FBgn0267550 | asRNA:CR45890 | 135 | 0.74 | 2.39E-03 | -0.09 | 4.80E-01 | 0.00 | 1.00E+00 |
| FBgn0267758 | asRNA:CR46089 | 20 | 0.02 | 9.50E-01 | 6.93 | 9.49E-07 | 0.01 | 1.00E+00 |
| FBgn0015591 | AstA | 15 | -0.47 | 4.28E-02 | -0.85 | 2.71E-02 | -0.03 | 7.69E-01 |
| FBgn0033010 | Atf6 | 1360 | -0.98 | 3.85E-07 | 0.00 | 9.95E-01 | 0.00 | 1.00E+00 |
| FBgn0037363 | Atg17 | 1079 | -0.92 | 3.17E-07 | 0.03 | 7.45E-01 | 0.00 | 1.00E+00 |
| FBgn0032422 | atilla | 38 | 0.68 | 1.38E-02 | -0.01 | 9.56E-01 | 0.00 | 1.00E+00 |
| FBgn0016120 | ATPsynD | 4001 | 0.89 | 1.85E-13 | -0.03 | 7.42E-01 | 0.00 | 1.00E+00 |
| FBgn0028342 | ATPsyndelta | 2959 | 1.17 | 4.25E-42 | 0.05 | 5.61E-01 | 0.00 | 1.00E+00 |
| FBgn0038224 | ATPsynE | 2430 | 0.91 | 4.10E-08 | -0.01 | 9.18E-01 | 0.00 | 1.00E+00 |
| FBgn0035032 | ATPsynF | 2536 | 0.99 | 9.61E-19 | 0.00 | 9.76E-01 | 0.00 | 1.00E+00 |
| FBgn0020235 | ATPsyngamma | 5712 | 0.73 | 9.57E-18 | 0.01 | 9.13E-01 | -0.01 | 1.00E+00 |
| FBgn0016691 | ATPsynO | 3667 | 0.75 | 2.40E-11 | 0.01 | 9.10E-01 | 0.00 | 1.00E+00 |
| FBgn0000150 | awd | 22227 | 0.68 | 1.45E-06 | 0.00 | 9.85E-01 | 0.00 | 1.00E+00 |
| FBgn0013751 | Awh | 61 | -0.10 | 7.33E-01 | -1.11 | 2.95E-03 | 0.00 | 1.00E+00 |
| FBgn0004870 | bab1 | 47 | 0.12 | 6.72E-01 | -1.35 | 7.56E-05 | 0.00 | 1.00E+00 |
| FBgn0011300 | babo | 2914 | -0.80 | 1.14E-09 | -0.04 | 7.06E-01 | 0.00 | 1.00E+00 |
| FBgn0031453 | Bacc | 14814 | -1.10 | 3.05E-25 | 0.05 | 5.22E-01 | 0.01 | 9.02E-01 |
| FBgn0031255 | BBS8 | 105 | 0.44 | 6.81E-02 | 0.88 | 5.53E-05 | 0.01 | 1.00E+00 |
| FBgn0052594 | be | 704 | -0.59 | 7.31E-05 | -0.07 | 4.14E-01 | 0.00 | 1.00E+00 |
| FBgn0000173 | ben | 4467 | -0.60 | 5.15E-06 | -0.04 | 7.26E-01 | 0.00 | 1.00E+00 |
| FBgn0260860 | Bet5 | 305 | 0.66 | 1.09E-03 | 0.02 | 8.60E-01 | 0.00 | 1.00E+00 |
| FBgn0284243 | betaTub56D | 87388 | 0.65 | 3.71E-15 | -0.02 | 8.83E-01 | -0.01 | 1.00E+00 |
| FBgn0035871 | Bl-1 | 1623 | -0.69 | 7.33E-12 | 0.30 | 5.15E-05 | 0.00 | 1.00E+00 |
| FBgn0000183 | BicD | 2135 | -0.68 | 7.68E-09 | 0.06 | 4.83E-01 | 0.00 | 1.00E+00 |
| FBgn0024491 | Bin1 | 1697 | 0.32 | 1.04E-01 | 2.12 | 1.19E-36 | 0.00 | 1.00E+00 |
| FBgn0026262 | bip2 | 3652 | -1.02 | 1.89E-14 | -0.07 | 3.85E-01 | 0.01 | 8.48E-01 |
| FBgn0085284 | Blos3 | 282 | 0.75 | 3.15E-04 | 0.04 | 6.56E-01 | 0.00 | 1.00E+00 |
| FBgn0036449 | bmm | 105 | 0.20 | 4.43E-01 | -1.35 | 1.15E-03 | -0.01 | 1.00E+00 |
| FBgn0037007 | BNIP3 | 2102 | -0.67 | 5.46E-06 | 0.02 | 8.57E-01 | 0.00 | 1.00E+00 |
| FBgn0014135 | bnl | 117 | -0.63 | 1.26E-02 | 0.02 | 8.64E-01 | 0.00 | 1.00E+00 |
| FBgn0261284 | bou | 1262 | 0.81 | 1.17E-08 | 0.09 | 2.46E-01 | 0.00 | 1.00E+00 |
| FBgn0050169 | Brca2 | 341 | -0.64 | 8.41E-05 | 0.00 | 9.81E-01 | 0.00 | 1.00E+00 |
| FBgn0000216 | Brd | 509 | 0.53 | 2.95E-03 | 1.17 | 7.95E-18 | 0.01 | 1.00E+00 |
| FBgn0264001 | bru3 | 51 | 0.05 | 8.22E-01 | 4.90 | 3.37E-15 | -0.01 | 1.00E+00 |
| FBgn0261822 | Bsg | 2702 | -0.77 | 1.00E-06 | 0.00 | 9.76E-01 | 0.00 | 1.00E+00 |
| FBgn0000229 | bsk | 1275 | -0.72 | 5.53E-12 | 0.03 | 7.89E-01 | 0.00 | 1.00E+00 |
| FBgn0000233 | btd | 63 | 0.21 | 3.76E-01 | -2.19 | 6.57E-06 | 0.01 | 1.00E+00 |
| FBgn0023096 | btv | 49 | -0.11 | 6.87E-01 | 0.58 | 6.50E-02 | 0.03 | 7.38E-01 |
| FBgn0259176 | bun | 5539 | -0.70 | 3.32E-07 | -0.41 | 7.00E-04 | 0.01 | 8.66E-01 |
| FBgn0000246 | c(3)G | 214 | -0.86 | 2.99E-05 | 0.06 | 5.01E-01 | 0.01 | 1.00E+00 |
| FBgn0004863 | C15 | 235 | 0.36 | 4.16E-02 | -0.61 | 5.84E-06 | 0.00 | 1.00E+00 |
| FBgn0263111 | cac | 124 | 0.37 | 1.37E-01 | 3.85 | 3.50E-29 | -0.02 | 9.52E-01 |
| FBgn0053653 | Cadps | 187 | 0.69 | 9.19E-03 | 1.58 | 2.50E-09 | 0.01 | 1.00E+00 |
| FBgn0030054 | Caf1-180 | 1658 | -0.58 | 2.62E-04 | 0.04 | 6.21E-01 | 0.01 | 1.00E+00 |
| FBgn0039928 | Cals | 4732 | -0.77 | 1.60E-05 | -0.03 | 8.06E-01 | 0.00 | 1.00E+00 |
| FBgn0000253 | Cam | 12315 | -1.48 | 3.62E-35 | 0.03 | 7.66E-01 | 0.00 | 1.00E+00 |
| FBgn0016126 | CaMKI | 3031 | -0.81 | 2.36E-10 | 0.00 | 9.77E-01 | 0.01 | 1.00E+00 |
| FBgn0264607 | CaMKII | 2738 | -0.98 | 1.88E-12 | 0.13 | 1.18E-01 | 0.08 | 2.06E-01 |
| FBgn0267912 | CanA-14F | 856 | -0.61 | 1.02E-04 | 0.04 | 6.99E-01 | 0.00 | 1.00E+00 |
| FBgn0037831 | Cap-H2 | 1038 | -0.59 | 1.70E-07 | 0.03 | 7.58E-01 | 0.07 | 2.13E-01 |
| FBgn0004878 | cas | 19 | 0.61 | 1.87E-02 | 0.38 | 8.83E-02 | 0.00 | 1.00E+00 |
| FBgn0285954 | caz | 3727 | -0.58 | 1.17E-03 | 0.06 | 5.27E-01 | 0.00 | 1.00E+00 |
| FBgn0031148 | Cbs | 2322 | 0.62 | 2.00E-17 | 0.05 | 5.13E-01 | 0.00 | 1.00E+00 |
| FBgn0030954 | CCKLR-17D3 | 380 | 0.67 | 3.15E-04 | 0.06 | 4.84E-01 | 0.00 | 1.00E+00 |
| FBgn0010621 | CCT5 | 11598 | 0.65 | 5.89E-18 | 0.05 | 5.39E-01 | -0.04 | 3.73E-01 |
| FBgn0052499 | Cda4 | 4755 | 0.63 | 7.21E-13 | 0.03 | 7.32E-01 | 0.02 | 6.49E-01 |
| FBgn0027491 | Cdk5alpha | 35 | -0.75 | 9.09E-03 | -0.09 | 3.69E-01 | 0.00 | 1.00E+00 |
| FBgn0034437 | CG10051 | 15 | -0.03 | 8.98E-01 | -2.23 | 8.63E-03 | -0.38 | 9.09E-02 |
| FBgn0036369 | CG10089 | 215 | -0.17 | 4.09E-01 | 0.84 | 4.59E-09 | -0.35 | 2.02E-02 |
| FBgn0032800 | CG10137 | 20 | 0.26 | 2.83E-01 | 0.65 | 3.90E-02 | 0.00 | 1.00E+00 |
| FBgn0032793 | CG10189 | 280 | 0.09 | 6.85E-01 | 0.61 | 7.79E-06 | 0.00 | 1.00E+00 |
| FBgn0038454 | CG10324 | 235 | -0.83 | 9.43E-06 | 0.06 | 4.80E-01 | 0.00 | 1.00E+00 |
| FBgn0032805 | CG10337 | 141 | -0.44 | 3.08E-02 | 0.94 | 4.60E-08 | 0.00 | 1.00E+00 |
| FBgn0034972 | CG10339 | 12 | 0.07 | 6.39E-01 | 2.10 | 6.08E-04 | 1.55 | 5.81E-03 |
| FBgn0034729 | CG10344 | 43 | -0.05 | 8.66E-01 | 1.22 | 1.10E-03 | 0.05 | 4.80E-01 |

| | | | | | | | | |
|---|---|---|---|---|---|---|---|---|
| FBgn0033019 | CG10395 | 486 | -0.99 | 3.91E-12 | 0.15 | 5.06E-02 | 0.00 | 1.00E+00 |
| FBgn0033021 | CG10417 | 4149 | -0.87 | 9.78E-43 | -0.05 | 4.73E-01 | -0.02 | 6.07E-01 |
| FBgn0036277 | CG10418 | 472 | 0.97 | 6.82E-07 | 0.03 | 8.02E-01 | 0.00 | 1.00E+00 |
| FBgn0037531 | CG10445 | 247 | -0.36 | 7.85E-02 | 0.81 | 6.20E-08 | 0.02 | 6.24E-01 |
| FBgn0033017 | CG10465 | 1501 | -0.72 | 2.95E-08 | -0.06 | 4.92E-01 | -0.01 | 1.00E+00 |
| FBgn0039312 | CG10514 | 39 | -1.49 | 7.99E-04 | 0.02 | 8.51E-01 | 0.00 | 1.00E+00 |
| FBgn0039323 | CG10559 | 50 | 0.14 | 5.95E-01 | 1.51 | 1.14E-08 | 0.00 | 1.00E+00 |
| FBgn0032717 | CG10600 | 1849 | -0.73 | 1.91E-05 | 0.01 | 9.49E-01 | 0.00 | 1.00E+00 |
| FBgn0036290 | CG10638 | 2684 | 0.14 | 2.79E-01 | 1.24 | 2.06E-62 | 0.00 | 1.00E+00 |
| FBgn0029666 | CG10803 | 1895 | -0.74 | 7.57E-06 | -0.03 | 7.80E-01 | 0.00 | 1.00E+00 |
| FBgn0027552 | CG10863 | 209 | 0.62 | 2.02E-02 | -0.03 | 7.37E-01 | 0.01 | 1.00E+00 |
| FBgn0034312 | CG10916 | 405 | -0.21 | 2.73E-01 | 1.32 | 2.49E-27 | 0.00 | 1.00E+00 |
| FBgn0033149 | CG11060 | 14 | 0.19 | 1.86E-01 | 0.63 | 4.56E-02 | 0.00 | 1.00E+00 |
| FBgn0039930 | CG11077 | 321 | 0.68 | 3.96E-04 | 0.04 | 6.86E-01 | 0.00 | 1.00E+00 |
| FBgn0030519 | CG11151 | 1004 | 0.66 | 1.09E-04 | -0.04 | 6.70E-01 | 0.00 | 1.00E+00 |
| FBgn0039927 | CG11155 | 89 | -0.20 | 4.49E-01 | 0.59 | 1.37E-02 | -0.01 | 1.00E+00 |
| FBgn0030511 | CG11158 | 89 | -0.09 | 7.52E-01 | 1.15 | 9.94E-05 | 0.00 | 1.00E+00 |
| FBgn0034528 | CG11180 | 768 | -1.31 | 7.99E-12 | 0.06 | 4.93E-01 | 0.00 | 1.00E+00 |
| FBgn0037115 | CG11249 | 31 | -0.27 | 2.75E-01 | 0.66 | 1.81E-02 | 0.00 | 1.00E+00 |
| FBgn0036334 | CG11267 | 2756 | 0.80 | 2.32E-09 | -0.01 | 9.69E-01 | 0.00 | 1.00E+00 |
| FBgn0035552 | CG11350 | 22 | 1.08 | 2.47E-03 | 0.01 | 9.10E-01 | 0.07 | 4.10E-01 |
| FBgn0039920 | CG11360 | 3374 | -0.59 | 2.82E-03 | 0.08 | 3.43E-01 | 0.01 | 1.00E+00 |
| FBgn0037181 | CG11370 | 1620 | 0.27 | 2.29E-01 | 0.76 | 6.42E-04 | 0.14 | 1.74E-01 |
| FBgn0031224 | CG11454 | 503 | 0.73 | 1.74E-06 | 0.04 | 6.54E-01 | 0.00 | 1.00E+00 |
| FBgn0037396 | CG11459 | 40 | 0.02 | 9.37E-01 | -3.59 | 5.30E-04 | -0.36 | 9.07E-02 |
| FBgn0031244 | CG11601 | 288 | -1.03 | 8.59E-12 | 0.02 | 8.70E-01 | 0.00 | 1.00E+00 |
| FBgn0036196 | CG11658 | 72 | 0.19 | 4.82E-01 | 0.80 | 4.86E-04 | 0.00 | 1.00E+00 |
| FBgn0030311 | CG11699 | 609 | 0.68 | 5.97E-04 | 0.03 | 7.59E-01 | 0.00 | 1.00E+00 |
| FBgn0030292 | CG11752 | 421 | 0.85 | 9.10E-06 | -0.02 | 8.49E-01 | 0.00 | 1.00E+00 |
| FBgn0037603 | CG11753 | 305 | 0.71 | 1.85E-04 | 0.00 | 9.72E-01 | 0.00 | 1.00E+00 |
| FBgn0031264 | CG11835 | 652 | -0.12 | 5.99E-01 | -0.96 | 6.46E-07 | -0.07 | 2.96E-01 |
| FBgn0039299 | CG11854 | 130 | -0.09 | 5.51E-01 | -0.89 | 3.53E-02 | -0.01 | 1.00E+00 |
| FBgn0014427 | CG11899 | 2687 | 0.58 | 2.36E-04 | -0.04 | 6.96E-01 | -0.04 | 4.40E-01 |
| FBgn0037312 | CG11999 | 699 | 0.71 | 3.22E-04 | 0.09 | 3.31E-01 | 0.00 | 1.00E+00 |
| FBgn0039831 | CG12054 | 2044 | -0.78 | 9.35E-06 | -0.01 | 9.62E-01 | 0.00 | 1.00E+00 |
| FBgn0030098 | CG12057 | 41 | 0.04 | 6.75E-01 | -3.13 | 4.56E-03 | -1.77 | 1.09E-02 |
| FBgn0037386 | CG1208 | 54 | 3.54 | 4.06E-08 | 1.32 | 3.43E-03 | 0.00 | 1.00E+00 |
| FBgn0030048 | CG12112 | 271 | -0.85 | 7.80E-06 | 0.12 | 1.48E-01 | 0.00 | 1.00E+00 |
| FBgn0030097 | CG12115 | 11 | -0.08 | 5.01E-01 | -5.99 | 2.31E-05 | -3.05 | 1.15E-03 |
| FBgn0037356 | CG12170 | 204 | 0.70 | 6.12E-04 | 0.03 | 8.04E-01 | 0.01 | 1.00E+00 |
| FBgn0037354 | CG12171 | 763 | 1.03 | 5.36E-13 | 0.75 | 1.51E-13 | 0.00 | 1.00E+00 |
| FBgn0030510 | CG12177 | 62 | 0.10 | 7.32E-01 | 0.76 | 2.59E-03 | 0.00 | 1.00E+00 |
| FBgn0031022 | CG12204 | 290 | 1.03 | 3.91E-07 | 0.43 | 5.96E-03 | 0.01 | 9.42E-01 |
| FBgn0037974 | CG12224 | 199 | -0.04 | 9.02E-01 | 2.01 | 3.75E-30 | 0.00 | 1.00E+00 |
| FBgn0036514 | CG12301 | 779 | -0.59 | 1.10E-03 | 0.09 | 2.87E-01 | 0.00 | 1.00E+00 |
| FBgn0038590 | CG12320 | 383 | 0.61 | 4.18E-04 | 0.06 | 4.68E-01 | 0.00 | 1.00E+00 |
| FBgn0033558 | CG12344 | 18 | -0.73 | 1.06E-02 | -0.07 | 5.45E-01 | 0.00 | 1.00E+00 |
| FBgn0029657 | CG12535 | 259 | -2.49 | 3.99E-31 | 0.04 | 6.98E-01 | 0.00 | 1.00E+00 |
| FBgn0250830 | CG12547 | 2267 | -0.82 | 2.09E-21 | 0.05 | 5.86E-01 | 0.00 | 1.00E+00 |
| FBgn0037811 | CG12592 | 84 | -5.00 | 3.37E-05 | -0.04 | 6.97E-01 | -0.01 | 1.00E+00 |
| FBgn0037796 | CG12814 | 1934 | -0.66 | 1.10E-11 | 0.05 | 5.48E-01 | 0.03 | 5.19E-01 |
| FBgn0033145 | CG12828 | 151 | 0.87 | 1.14E-04 | -0.05 | 6.52E-01 | -0.02 | 8.92E-01 |
| FBgn0033945 | CG12868 | 154 | -0.36 | 1.17E-01 | 1.52 | 3.95E-16 | 0.00 | 1.00E+00 |
| FBgn0033547 | CG12935 | 370 | 1.10 | 1.08E-10 | -0.02 | 8.88E-01 | 0.00 | 1.00E+00 |
| FBgn0037753 | CG12947 | 202 | -0.18 | 4.28E-01 | -0.63 | 3.27E-04 | 0.00 | 1.00E+00 |
| FBgn0035501 | CG1299 | 414 | 0.26 | 2.45E-01 | 1.01 | 2.72E-08 | 0.00 | 1.00E+00 |
| FBgn0030859 | CG12990 | 39 | -1.81 | 3.29E-05 | 0.02 | 8.45E-01 | 0.00 | 1.00E+00 |
| FBgn0036677 | CG13023 | 385 | 2.05 | 7.22E-11 | -0.32 | 5.69E-02 | -0.14 | 1.79E-01 |
| FBgn0036670 | CG13029 | 41 | 0.35 | 1.31E-01 | 2.92 | 1.84E-18 | 0.12 | 2.40E-01 |
| FBgn0036605 | CG13041 | 54 | 1.02 | 3.83E-03 | -0.07 | 4.61E-01 | -0.40 | 3.96E-02 |
| FBgn0036599 | CG13044 | 661 | 0.68 | 4.80E-03 | 0.05 | 5.92E-01 | 0.02 | 8.05E-01 |
| FBgn0036596 | CG13045 | 48 | -0.59 | 2.66E-02 | -0.06 | 5.40E-01 | -0.01 | 1.00E+00 |
| FBgn0036660 | CG13045 | 470 | -0.83 | 4.04E-09 | 0.18 | 4.71E-02 | 0.13 | 1.00E-01 |
| FBgn0040801 | CG13053 | 118 | 0.97 | 1.90E-03 | -0.96 | 9.37E-03 | -1.91 | 1.24E-04 |
| FBgn0040796 | CG13064 | 52 | 0.81 | 5.07E-03 | 0.05 | 6.12E-01 | 0.00 | 1.00E+00 |
| FBgn0036589 | CG13067 | 78 | 0.87 | 4.69E-03 | 0.06 | 5.01E-01 | 1.27 | 1.22E-05 |
| FBgn0032803 | CG13082 | 663 | 0.62 | 4.19E-03 | -0.07 | 4.59E-01 | 0.00 | 1.00E+00 |
| FBgn0032051 | CG13097 | 1071 | -0.63 | 3.97E-06 | 0.08 | 3.18E-01 | 0.00 | 1.00E+00 |
| FBgn0033608 | CG13220 | 1046 | 0.59 | 1.39E-06 | 0.05 | 5.80E-01 | 0.00 | 1.00E+00 |
| FBgn0033781 | CG13319 | 429 | 0.85 | 1.59E-05 | 0.03 | 7.98E-01 | 0.00 | 1.00E+00 |
| FBgn0029531 | CG13362 | 69 | -0.74 | 3.92E-03 | 0.03 | 7.39E-01 | -0.04 | 6.33E-01 |
| FBgn0026879 | CG13364 | 1664 | 0.76 | 6.87E-06 | 0.00 | 9.93E-01 | 0.00 | 1.00E+00 |
| FBgn0025640 | CG13369 | 355 | 0.63 | 8.83E-04 | -0.01 | 9.12E-01 | 0.00 | 1.00E+00 |
| FBgn0040658 | CG13516 | 84 | 0.91 | 8.75E-04 | -2.22 | 4.86E-15 | -3.17 | 1.48E-27 |
| FBgn0035010 | CG13579 | 278 | -0.03 | 8.28E-01 | 11.51 | 4.85E-23 | 0.00 | 1.00E+00 |
| FBgn0035020 | CG13585 | 575 | 0.95 | 4.94E-11 | 0.13 | 1.02E-01 | 0.02 | 7.52E-01 |
| FBgn0039219 | CG13630 | 1091 | 0.64 | 1.67E-09 | 0.10 | 1.19E-01 | 0.00 | 1.00E+00 |
| FBgn0035859 | CG13678 | 134 | 1.06 | 1.15E-03 | 0.02 | 8.64E-01 | 0.09 | 3.13E-01 |
| FBgn0030539 | CG1368 | 104 | 0.83 | 4.40E-03 | -1.49 | 1.89E-04 | -0.01 | 1.00E+00 |

| | | | | | | | | |
|---|---|---|---|---|---|---|---|---|
| FBgn0031254 | CG13692 | 54 | 0.04 | 8.97E-01 | 1.31 | 2.65E-05 | 0.01 | 1.00E+00 |
| FBgn0035553 | CG13722 | 58 | 0.93 | 4.69E-03 | -0.23 | 1.74E-01 | -0.13 | 2.10E-01 |
| FBgn0036382 | CG13737 | 57 | 0.62 | 1.26E-02 | 0.04 | 6.72E-01 | 0.00 | 1.00E+00 |
| FBgn0033340 | CG13751 | 510 | 1.29 | 9.93E-08 | 0.03 | 7.26E-01 | 0.00 | 1.00E+00 |
| FBgn0031834 | CG13766 | 269 | -1.05 | 7.32E-11 | -0.06 | 5.06E-01 | 0.00 | 1.00E+00 |
| FBgn0035176 | CG13905 | 32 | 1.11 | 1.36E-03 | 0.01 | 9.54E-01 | 0.00 | 1.00E+00 |
| FBgn0040786 | CG14104 | 193 | 1.28 | 2.01E-07 | -0.08 | 3.84E-01 | -0.01 | 1.00E+00 |
| FBgn0040817 | CG14132 | 340 | 0.11 | 6.73E-01 | -0.62 | 1.97E-03 | -0.03 | 5.64E-01 |
| FBgn0038658 | CG14292 | 69 | -1.15 | 1.15E-03 | 0.06 | 6.08E-01 | 0.00 | 1.00E+00 |
| FBgn0038647 | CG14302 | 16 | -0.04 | 7.60E-01 | -1.37 | 2.08E-02 | -0.01 | 1.00E+00 |
| FBgn0038629 | CG14304 | 297 | 0.08 | 7.08E-01 | 0.62 | 1.63E-05 | 0.00 | 1.00E+00 |
| FBgn0038581 | CG14314 | 75 | -0.47 | 5.04E-02 | 1.19 | 4.26E-05 | 0.00 | 1.00E+00 |
| FBgn0038148 | CG14377 | 80 | -1.02 | 2.52E-04 | -0.09 | 4.73E-01 | -0.01 | 1.00E+00 |
| FBgn0032900 | CG14401 | 26 | -0.99 | 2.51E-03 | -0.02 | 8.90E-01 | -0.01 | 1.00E+00 |
| FBgn0030584 | CG14407 | 427 | 0.78 | 1.12E-04 | 0.00 | 9.86E-01 | 0.00 | 1.00E+00 |
| FBgn0029894 | CG14440 | 648 | -0.66 | 2.53E-05 | -0.01 | 9.13E-01 | 0.00 | 1.00E+00 |
| FBgn0033000 | CG14464 | 898 | -0.68 | 2.86E-05 | 0.04 | 7.03E-01 | 0.00 | 1.00E+00 |
| FBgn0037127 | CG14566 | 326 | 1.00 | 4.07E-05 | -0.02 | 8.40E-01 | 0.52 | 1.64E-02 |
| FBgn0037503 | CG14598 | 331 | 1.24 | 1.79E-12 | -0.18 | 6.31E-02 | 0.00 | 1.00E+00 |
| FBgn0031184 | CG14615 | 61 | -0.84 | 2.10E-03 | -0.02 | 8.61E-01 | -0.03 | 6.96E-01 |
| FBgn0037850 | CG14695 | 97 | 0.09 | 5.32E-01 | 7.33 | 1.55E-43 | 0.03 | 7.36E-01 |
| FBgn0037930 | CG14715 | 686 | 0.87 | 8.24E-08 | -0.03 | 7.95E-01 | -0.01 | 1.00E+00 |
| FBgn0033243 | CG14763 | 94 | -0.69 | 4.53E-03 | -0.06 | 6.21E-01 | 0.00 | 1.00E+00 |
| FBgn0038455 | CG14907 | 184 | -0.29 | 2.10E-01 | 0.97 | 2.91E-09 | -0.13 | 1.54E-01 |
| FBgn0032335 | CG14915 | 24 | 0.28 | 2.37E-01 | 0.64 | 2.32E-02 | 0.02 | 9.05E-01 |
| FBgn0035409 | CG14963 | 27 | 0.00 | 9.87E-01 | 0.01 | 9.11E-01 | 0.69 | 4.79E-02 |
| FBgn0035414 | CG14965 | 340 | -0.68 | 9.14E-05 | -0.02 | 8.87E-01 | 0.00 | 1.00E+00 |
| FBgn0035415 | CG14966 | 318 | -0.70 | 7.73E-05 | -0.03 | 7.86E-01 | -0.01 | 9.03E-01 |
| FBgn0035469 | CG14977 | 355 | 0.66 | 3.70E-04 | 0.01 | 9.69E-01 | -0.01 | 1.00E+00 |
| FBgn0035480 | CG14984 | 277 | 0.75 | 9.39E-04 | 0.23 | 5.87E-02 | 0.01 | 1.00E+00 |
| FBgn0034399 | CG15083 | 584 | 0.79 | 1.63E-04 | 0.02 | 8.17E-01 | 0.00 | 1.00E+00 |
| FBgn0032733 | CG15170 | 18 | 0.02 | 9.12E-01 | -2.23 | 8.48E-03 | -0.18 | 1.79E-01 |
| FBgn0030234 | CG15211 | 42 | 0.60 | 2.38E-02 | 0.00 | 9.86E-01 | 0.01 | 1.00E+00 |
| FBgn0033104 | CG15237 | 463 | 0.71 | 3.29E-05 | 0.03 | 8.00E-01 | 0.00 | 1.00E+00 |
| FBgn0030183 | CG15309 | 348 | -1.15 | 2.11E-07 | 0.03 | 7.91E-01 | 0.00 | 1.00E+00 |
| FBgn0030029 | CG15343 | 65 | -0.08 | 7.69E-01 | -1.24 | 6.05E-03 | -0.01 | 1.00E+00 |
| FBgn0030040 | CG15347 | 54 | 4.74 | 6.59E-09 | 0.00 | 9.83E-01 | -0.03 | 7.54E-01 |
| FBgn0040718 | CG15353 | 3826 | 0.82 | 1.11E-05 | 0.03 | 7.62E-01 | 0.61 | 2.02E-03 |
| FBgn0031393 | CG15382 | 81 | -0.62 | 8.94E-03 | -0.11 | 2.88E-01 | -0.01 | 1.00E+00 |
| FBgn0039828 | CG1542 | 1295 | 0.67 | 1.31E-06 | 0.06 | 5.20E-01 | -0.01 | 1.00E+00 |
| FBgn0029719 | CG15473 | 26 | 0.74 | 8.74E-03 | 0.00 | 9.77E-01 | 0.00 | 1.00E+00 |
| FBgn0039807 | CG15546 | 179 | 0.73 | 3.79E-04 | 0.75 | 4.93E-04 | 0.03 | 6.76E-01 |
| FBgn0031632 | CG15628 | 830 | -0.69 | 6.10E-05 | -0.02 | 8.10E-01 | 0.02 | 6.86E-01 |
| FBgn0030309 | CG1572 | 778 | 0.65 | 7.98E-06 | -0.10 | 1.88E-01 | 0.00 | 1.00E+00 |
| FBgn0029766 | CG15784 | 8033 | 0.41 | 1.73E-02 | 5.32 | 0.00E+00 | 0.61 | 6.92E-05 |
| FBgn0030246 | CG1582 | 1397 | 0.03 | 8.89E-01 | 0.74 | 2.03E-15 | 0.01 | 1.00E+00 |
| FBgn0038814 | CG15923 | 256 | -1.32 | 5.53E-08 | 0.07 | 5.34E-01 | 0.01 | 1.00E+00 |
| FBgn0033191 | CG1598 | 1089 | 0.60 | 9.56E-06 | 0.05 | 5.99E-01 | 0.00 | 1.00E+00 |
| FBgn0039602 | CG1647 | 659 | -0.61 | 1.64E-05 | 0.03 | 7.46E-01 | 0.01 | 1.00E+00 |
| FBgn0038719 | CG16727 | 101 | -13.72 | 9.08E-07 | 0.00 | 9.77E-01 | 0.00 | 1.00E+00 |
| FBgn0032488 | CG16812 | 796 | -0.63 | 2.20E-08 | 0.05 | 5.27E-01 | 0.00 | 1.00E+00 |
| FBgn0032495 | CG16820 | 141 | -0.22 | 3.58E-01 | 1.11 | 4.35E-10 | -0.63 | 3.41E-03 |
| FBgn0030321 | CG1703 | 2733 | -0.59 | 3.11E-05 | 0.17 | 5.48E-02 | 0.00 | 1.00E+00 |
| FBgn0040496 | CG17104 | 334 | -0.16 | 5.23E-01 | 1.06 | 2.36E-07 | 0.00 | 1.00E+00 |
| FBgn0036440 | CG17177 | 11 | -0.03 | 8.46E-01 | 6.67 | 3.10E-06 | 0.00 | 1.00E+00 |
| FBgn0038043 | CG17202 | 1061 | 0.69 | 2.76E-04 | 0.01 | 9.18E-01 | 0.00 | 1.00E+00 |
| FBgn0039977 | CG17454 | 820 | -0.64 | 7.83E-09 | 0.13 | 4.25E-02 | 0.00 | 1.00E+00 |
| FBgn0032997 | CG17486 | 1168 | -0.59 | 1.85E-06 | 0.00 | 9.88E-01 | 0.00 | 1.00E+00 |
| FBgn0039959 | CG17514 | 7525 | -0.60 | 2.64E-04 | 0.03 | 7.60E-01 | 0.00 | 1.00E+00 |
| FBgn0261387 | CG17528 | 2579 | -0.62 | 4.04E-09 | -0.01 | 9.46E-01 | 0.00 | 1.00E+00 |
| FBgn0033777 | CG17574 | 551 | 0.29 | 8.95E-02 | 0.91 | 1.94E-17 | -0.67 | 1.94E-08 |
| FBgn0031597 | CG17612 | 367 | -0.72 | 9.48E-06 | 0.04 | 7.15E-01 | 0.01 | 1.00E+00 |
| FBgn0263780 | CG17684 | 127 | -0.31 | 1.98E-01 | 1.48 | 1.26E-18 | -0.11 | 1.79E-01 |
| FBgn0040056 | CG17698 | 1191 | -0.96 | 1.17E-14 | -0.06 | 4.28E-01 | 0.00 | 1.00E+00 |
| FBgn0033802 | CG17724 | 203 | -1.43 | 3.95E-06 | 0.06 | 4.74E-01 | 0.00 | 1.00E+00 |
| FBgn0038718 | CG17752 | 65 | -2.13 | 4.83E-04 | 0.02 | 8.61E-01 | 0.00 | 1.00E+00 |
| FBgn0040899 | CG17776 | 374 | 1.06 | 1.51E-06 | 0.02 | 8.65E-01 | 0.00 | 1.00E+00 |
| FBgn0040005 | CG17883 | 1756 | -0.62 | 7.07E-07 | 0.09 | 2.41E-01 | 0.00 | 1.00E+00 |
| FBgn0028856 | CG18063 | 238 | -0.03 | 8.28E-01 | 8.80 | 7.19E-73 | -0.03 | 7.66E-01 |
| FBgn0038470 | CG18213 | 611 | -0.08 | 7.21E-01 | 0.82 | 1.62E-08 | 0.00 | 1.00E+00 |
| FBgn0033431 | CG1827 | 277 | 1.05 | 3.97E-06 | -0.04 | 6.56E-01 | 0.00 | 1.00E+00 |
| FBgn0030351 | CG1840 | 156 | -0.65 | 5.29E-03 | -0.01 | 9.48E-01 | 0.00 | 1.00E+00 |
| FBgn0037973 | CG18547 | 919 | 0.26 | 1.35E-01 | 1.90 | 8.11E-70 | -0.01 | 1.00E+00 |
| FBgn0031929 | CG18585 | 13 | -0.02 | 8.82E-01 | -1.09 | 2.80E-02 | -0.09 | 3.38E-01 |
| FBgn0033421 | CG1888 | 455 | 0.63 | 3.47E-04 | 0.01 | 9.03E-01 | 0.00 | 1.00E+00 |
| FBgn0039911 | CG1909 | 92 | 0.51 | 4.75E-02 | 3.31 | 4.98E-37 | 0.04 | 5.75E-01 |
| FBgn0022349 | CG1910 | 5653 | -0.77 | 1.49E-09 | -0.01 | 9.26E-01 | 0.00 | 1.00E+00 |
| FBgn0039881 | CG1971 | 10 | -0.03 | 9.12E-01 | 0.70 | 3.73E-02 | 0.00 | 1.00E+00 |
| FBgn0039886 | CG2003 | 66 | 0.01 | 9.87E-01 | -0.65 | 4.84E-03 | -0.09 | 2.74E-01 |

| | | | | | | | | |
|---|---|---|---|---|---|---|---|---|
| FBgn0039664 | CG2006 | 513 | 0.76 | 3.00E-05 | 0.01 | 9.46E-01 | 0.00 | 1.00E+00 |
| FBgn0035271 | CG2021 | 304 | 0.65 | 5.50E-04 | 0.02 | 8.14E-01 | 0.00 | 1.00E+00 |
| FBgn0033205 | CG2064 | 103 | -0.13 | 6.35E-01 | 1.83 | 2.85E-21 | 0.00 | 1.00E+00 |
| FBgn0017448 | CG2187 | 56 | 0.76 | 7.19E-03 | 1.48 | 2.14E-05 | 1.25 | 4.85E-04 |
| FBgn0030447 | CG2200 | 465 | 0.66 | 5.82E-05 | -0.04 | 7.06E-01 | -0.01 | 1.00E+00 |
| FBgn0039665 | CG2310 | 1991 | 0.82 | 6.19E-05 | 0.02 | 8.88E-01 | 0.00 | 1.00E+00 |
| FBgn0034931 | CG2812 | 789 | 0.77 | 1.36E-07 | 0.55 | 9.49E-07 | 0.52 | 2.63E-05 |
| FBgn0023526 | CG2865 | 935 | -0.93 | 1.42E-06 | 0.02 | 8.50E-01 | 0.01 | 1.00E+00 |
| FBgn0029679 | CG2901 | 32 | 1.17 | 2.64E-03 | 0.06 | 5.87E-01 | -0.01 | 1.00E+00 |
| FBgn0030189 | CG2909 | 129 | 0.00 | 9.91E-01 | 1.71 | 1.07E-19 | 0.01 | 1.00E+00 |
| FBgn0023528 | CG2924 | 2315 | -0.73 | 1.93E-12 | -0.05 | 5.33E-01 | 0.00 | 1.00E+00 |
| FBgn0031643 | CG3008 | 1831 | -0.27 | 1.20E-02 | 1.75 | 8.62E-147 | 0.01 | 9.62E-01 |
| FBgn0050187 | CG30187 | 266 | -0.12 | 6.49E-01 | 1.68 | 1.49E-24 | 0.00 | 1.00E+00 |
| FBgn0050196 | CG30196 | 60 | 0.19 | 2.15E-01 | 2.12 | 1.75E-11 | 0.08 | 3.26E-01 |
| FBgn0050197 | CG30197 | 45 | 0.25 | 3.20E-01 | -2.87 | 5.43E-10 | 0.00 | 1.00E+00 |
| FBgn0031645 | CG3036 | 831 | 0.87 | 2.51E-06 | 0.65 | 6.15E-05 | -0.01 | 8.88E-01 |
| FBgn0050424 | CG30424 | 15 | -0.20 | 3.78E-01 | -1.39 | 6.47E-03 | -0.01 | 1.00E+00 |
| FBgn0050428 | CG30428 | 190 | -0.09 | 7.38E-01 | 0.65 | 6.52E-04 | 0.86 | 1.95E-05 |
| FBgn0050440 | CG30440 | 2603 | -0.80 | 1.00E-14 | -0.07 | 3.70E-01 | 0.00 | 1.00E+00 |
| FBgn0034816 | CG3085 | 164 | 0.45 | 2.53E-02 | 0.73 | 1.11E-06 | 0.00 | 1.00E+00 |
| FBgn0051030 | CG31030 | 110 | -0.64 | 1.41E-02 | -0.16 | 1.63E-01 | 0.00 | 1.00E+00 |
| FBgn0051055 | CG31055 | 196 | 0.70 | 1.15E-03 | -0.04 | 7.60E-01 | 0.00 | 1.00E+00 |
| FBgn0033005 | CG3107 | 2743 | -0.71 | 1.56E-07 | -0.01 | 9.10E-01 | 0.00 | 1.00E+00 |
| FBgn0051076 | CG31076 | 39 | 0.06 | 8.28E-01 | -0.05 | 5.92E-01 | -2.80 | 1.70E-05 |
| FBgn0051126 | CG31126 | 212 | 0.76 | 3.03E-04 | 0.02 | 8.97E-01 | 0.00 | 1.00E+00 |
| FBgn0047114 | CG31142 | 452 | 0.13 | 5.07E-01 | 0.59 | 2.76E-05 | -0.01 | 9.42E-01 |
| FBgn0051156 | CG31156 | 386 | -0.80 | 1.01E-06 | 0.03 | 7.46E-01 | 0.00 | 1.00E+00 |
| FBgn0051157 | CG31157 | 19 | 4.04 | 1.52E-04 | 0.08 | 5.01E-01 | 0.01 | 1.00E+00 |
| FBgn0051391 | CG31391 | 23 | -0.69 | 1.21E-02 | 0.00 | 9.88E-01 | 0.00 | 1.00E+00 |
| FBgn0051431 | CG31431 | 21 | -0.13 | 5.83E-01 | -0.81 | 2.70E-02 | 0.00 | 1.00E+00 |
| FBgn0051467 | CG31467 | 117 | -0.64 | 7.42E-03 | 0.00 | 9.83E-01 | 0.00 | 1.00E+00 |
| FBgn0051510 | CG31510 | 942 | -1.10 | 4.98E-15 | -0.09 | 2.64E-01 | 0.00 | 1.00E+00 |
| FBgn0051548 | CG31548 | 553 | 0.68 | 1.35E-05 | 0.02 | 8.45E-01 | 0.00 | 1.00E+00 |
| FBgn0051549 | CG31549 | 582 | 0.18 | 2.97E-01 | 0.93 | 1.89E-23 | 0.00 | 1.00E+00 |
| FBgn0051559 | CG31559 | 89 | 0.07 | 8.26E-01 | -0.59 | 2.16E-02 | -0.02 | 8.81E-01 |
| FBgn0051633 | CG31633 | 109 | 0.66 | 6.96E-03 | 0.14 | 1.73E-01 | 0.01 | 1.00E+00 |
| FBgn0029896 | CG3168 | 768 | 0.19 | 4.55E-01 | 1.17 | 3.76E-06 | 0.00 | 1.00E+00 |
| FBgn0051898 | CG31898 | 120 | -0.71 | 1.78E-03 | 0.04 | 7.04E-01 | 0.01 | 1.00E+00 |
| FBgn0031360 | CG31937 | 567 | 0.78 | 1.14E-07 | 0.01 | 9.51E-01 | 0.00 | 1.00E+00 |
| FBgn0051955 | CG31955 | 41 | 0.02 | 9.57E-01 | -0.16 | 2.03E-01 | -1.28 | 1.29E-03 |
| FBgn0051997 | CG31997 | 2776 | 0.64 | 9.55E-06 | 0.08 | 3.37E-01 | -0.14 | 9.82E-02 |
| FBgn0051998 | CG31998 | 3283 | -0.73 | 1.38E-04 | 0.00 | 9.83E-01 | 0.01 | 1.00E+00 |
| FBgn0051999 | CG31999 | 642 | 0.48 | 4.40E-04 | 0.94 | 7.82E-23 | -0.52 | 3.45E-06 |
| FBgn0052017 | CG32017 | 50 | -0.13 | 6.30E-01 | 1.88 | 8.72E-06 | 0.03 | 6.95E-01 |
| FBgn0052055 | CG32055 | 76 | 0.88 | 1.45E-03 | -1.68 | 1.88E-07 | -2.31 | 1.15E-12 |
| FBgn0052069 | CG32069 | 283 | 0.66 | 8.71E-04 | 0.00 | 9.89E-01 | -0.01 | 1.00E+00 |
| FBgn0052095 | CG32095 | 635 | -0.64 | 3.32E-07 | 0.01 | 9.67E-01 | 0.00 | 1.00E+00 |
| FBgn0052163 | CG32163 | 213 | 0.66 | 1.33E-03 | 0.01 | 9.06E-01 | 0.00 | 1.00E+00 |
| FBgn0052176 | CG32176 | 526 | -0.83 | 1.30E-07 | 0.02 | 8.39E-01 | 0.00 | 1.00E+00 |
| FBgn0052243 | CG32243 | 1236 | -0.76 | 1.70E-15 | 0.01 | 9.47E-01 | -0.01 | 1.00E+00 |
| FBgn0052354 | CG32354 | 120 | 0.18 | 4.83E-01 | 0.68 | 4.05E-03 | 0.73 | 5.80E-03 |
| FBgn0041706 | CG3253 | 101 | 0.07 | 8.05E-01 | 1.71 | 4.60E-19 | -0.01 | 9.62E-01 |
| FBgn0052532 | CG32532 | 10 | -0.17 | 4.17E-01 | 5.19 | 1.02E-04 | 4.29 | 1.22E-03 |
| FBgn0052544 | CG32544 | 30 | -0.69 | 1.34E-02 | 0.00 | 9.85E-01 | -0.01 | 1.00E+00 |
| FBgn0052549 | CG32549 | 668 | 0.19 | 4.29E-01 | 1.12 | 1.39E-07 | -0.02 | 8.49E-01 |
| FBgn0032986 | CG3262 | 1798 | -0.61 | 1.50E-16 | 0.23 | 1.11E-05 | 0.02 | 5.82E-01 |
| FBgn0052625 | CG32625 | 10 | -0.04 | 8.29E-01 | 2.22 | 9.92E-04 | 0.00 | 1.00E+00 |
| FBgn0052694 | CG32694 | 95 | 0.22 | 3.56E-01 | -0.84 | 2.94E-04 | 0.00 | 1.00E+00 |
| FBgn0052850 | CG32850 | 772 | -1.46 | 4.07E-37 | 0.05 | 5.76E-01 | 0.00 | 1.00E+00 |
| FBgn0053225 | CG33225 | 21 | 0.60 | 1.61E-02 | 0.15 | 2.63E-01 | 0.00 | 1.00E+00 |
| FBgn0053281 | CG33281 | 14 | -10.96 | 7.84E-06 | 0.05 | 6.42E-01 | 0.01 | 1.00E+00 |
| FBgn0053509 | CG33509 | 743 | -0.64 | 6.34E-04 | 1.82 | 4.17E-46 | 0.19 | 6.27E-02 |
| FBgn0031619 | CG3355 | 63 | 0.66 | 1.35E-02 | -0.04 | 6.88E-01 | 0.00 | 1.00E+00 |
| FBgn0052831 | CG33695 | 761 | 0.62 | 7.88E-05 | -0.02 | 8.24E-01 | 0.00 | 1.00E+00 |
| FBgn0262731 | CG33941 | 610 | -1.36 | 5.20E-10 | 0.05 | 5.38E-01 | 0.00 | 1.00E+00 |
| FBgn0053978 | CG33978 | 6959 | -0.69 | 2.06E-04 | 0.11 | 2.26E-01 | 0.04 | 4.82E-01 |
| FBgn0085192 | CG34163 | 279 | -0.70 | 1.49E-03 | 0.00 | 9.76E-01 | 0.00 | 1.00E+00 |
| FBgn0033100 | CG3420 | 534 | 0.60 | 6.60E-04 | 0.03 | 8.04E-01 | 0.00 | 1.00E+00 |
| FBgn0085236 | CG34207 | 212 | -0.84 | 3.04E-05 | -0.02 | 8.28E-01 | 0.00 | 1.00E+00 |
| FBgn0085276 | CG34247 | 87 | 4.23 | 8.81E-18 | 0.05 | 5.80E-01 | -1.88 | 7.16E-04 |
| FBgn0085411 | CG34382 | 59 | 0.85 | 1.89E-03 | -1.48 | 1.98E-05 | -1.22 | 2.36E-04 |
| FBgn0085452 | CG34423 | 18 | 0.32 | 1.61E-01 | -0.96 | 1.15E-02 | -1.98 | 2.34E-04 |
| FBgn0035996 | CG3448 | 164 | -0.30 | 1.51E-01 | 1.45 | 4.49E-18 | -0.01 | 1.00E+00 |
| FBgn0024984 | CG3457 | 47 | 1.34 | 1.07E-04 | 1.75 | 3.05E-08 | 0.00 | 1.00E+00 |
| FBgn0035007 | CG3492 | 22 | -0.03 | 8.41E-01 | 7.08 | 6.14E-09 | -0.01 | 1.00E+00 |
| FBgn0035008 | CG3494 | 13 | 0.00 | 1.00E+00 | 7.85 | 3.38E-08 | 0.00 | 1.00E+00 |
| FBgn0034849 | CG3500 | 293 | 0.62 | 1.41E-03 | 0.01 | 9.05E-01 | 0.00 | 1.00E+00 |
| FBgn0035063 | CG3594 | 504 | 0.59 | 1.16E-03 | 0.06 | 4.53E-01 | 0.00 | 1.00E+00 |
| FBgn0029824 | CG3726 | 143 | 0.27 | 2.31E-01 | 0.78 | 5.21E-05 | 0.03 | 5.79E-01 |

| | | | | | | | | |
|---|---|---|---|---|---|---|---|---|
| FBgn0035085 | CG3770 | 365 | 0.18 | 3.02E-01 | -0.79 | 4.13E-11 | -0.03 | 5.47E-01 |
| FBgn0036422 | CG3868 | 17 | -0.06 | 6.16E-01 | -1.60 | 1.44E-02 | -0.28 | 1.22E-01 |
| FBgn0058160 | CG40160 | 3127 | -0.80 | 2.78E-16 | -0.05 | 5.08E-01 | -0.01 | 9.40E-01 |
| FBgn0058191 | CG40191 | 1684 | -0.60 | 5.47E-08 | -0.06 | 4.68E-01 | 0.00 | 1.00E+00 |
| FBgn0035986 | CG4022 | 1651 | -0.83 | 7.91E-10 | -0.06 | 4.64E-01 | -0.01 | 9.41E-01 |
| FBgn0063670 | CG40228 | 993 | -1.01 | 2.80E-10 | 0.02 | 8.24E-01 | 0.00 | 1.00E+00 |
| FBgn0039955 | CG41099 | 2624 | -0.75 | 2.20E-08 | -0.07 | 4.01E-01 | 0.00 | 1.00E+00 |
| FBgn0087011 | CG41520 | 382 | -0.08 | 7.78E-01 | 0.72 | 3.67E-04 | -0.02 | 7.95E-01 |
| FBgn0038302 | CG4210 | 304 | 0.06 | 8.08E-01 | 0.84 | 6.58E-11 | 0.01 | 9.40E-01 |
| FBgn0250862 | CG42237 | 12 | -0.60 | 2.25E-02 | -0.05 | 6.20E-01 | 0.00 | 1.00E+00 |
| FBgn0259143 | CG42258 | 765 | -1.18 | 1.27E-20 | 0.02 | 8.81E-01 | 0.00 | 1.00E+00 |
| FBgn0266569 | CG42259 | 221 | -0.77 | 2.05E-03 | 0.02 | 8.22E-01 | 0.00 | 1.00E+00 |
| FBgn0259222 | CG42322 | 26 | -0.66 | 1.61E-02 | 0.03 | 8.02E-01 | 0.01 | 1.00E+00 |
| FBgn0259742 | CG42360 | 459 | -0.79 | 9.71E-08 | 0.07 | 4.15E-01 | 0.00 | 1.00E+00 |
| FBgn0259711 | CG42365 | 141 | -0.21 | 3.83E-01 | 1.18 | 2.56E-11 | 0.00 | 1.00E+00 |
| FBgn0034761 | CG4250 | 49 | 0.70 | 1.17E-02 | 0.09 | 3.74E-01 | 0.01 | 1.00E+00 |
| FBgn0260768 | CG42566 | 16 | 0.14 | 5.66E-01 | -1.69 | 2.81E-03 | -0.84 | 1.90E-02 |
| FBgn0034741 | CG4269 | 32 | -1.33 | 1.99E-03 | -0.02 | 8.60E-01 | 0.01 | 1.00E+00 |
| FBgn0261584 | CG42694 | 19 | -0.16 | 5.02E-01 | -0.86 | 2.34E-02 | 0.00 | 1.00E+00 |
| FBgn0261641 | CG42724 | 4450 | -0.87 | 2.78E-09 | 0.02 | 8.50E-01 | 0.00 | 1.00E+00 |
| FBgn0261995 | CG42813 | 479 | 0.60 | 7.35E-04 | 0.01 | 9.46E-01 | 0.00 | 1.00E+00 |
| FBgn0034742 | CG4294 | 1630 | -0.72 | 3.08E-08 | 0.18 | 3.61E-02 | 0.02 | 7.92E-01 |
| FBgn0038784 | CG4362 | 57 | 0.04 | 8.38E-01 | -1.24 | 1.81E-02 | -0.59 | 5.15E-02 |
| FBgn0263993 | CG43736 | 3291 | -0.60 | 2.45E-04 | -0.03 | 7.64E-01 | 0.00 | 1.00E+00 |
| FBgn0032132 | CG4382 | 1610 | 1.20 | 1.55E-11 | -0.43 | 3.97E-03 | -0.64 | 3.44E-04 |
| FBgn0038771 | CG4390 | 2352 | 0.60 | 4.28E-06 | -0.01 | 9.12E-01 | -0.01 | 1.00E+00 |
| FBgn0264748 | CG44006 | 16 | -0.19 | 3.72E-01 | 0.85 | 1.86E-02 | 0.00 | 1.00E+00 |
| FBgn0264911 | CG44102 | 35 | 0.02 | 9.40E-01 | 2.66 | 5.21E-15 | 0.00 | 1.00E+00 |
| FBgn0265413 | CG44325 | 1479 | 0.60 | 1.75E-04 | -0.03 | 7.71E-01 | 0.00 | 1.00E+00 |
| FBgn0266101 | CG44838 | 1266 | -0.58 | 3.40E-05 | 0.00 | 9.80E-01 | 0.00 | 1.00E+00 |
| FBgn0031896 | CG4502 | 1190 | -0.65 | 9.37E-07 | -0.07 | 3.45E-01 | 0.00 | 1.00E+00 |
| FBgn0266410 | CG45050 | 4461 | -1.41 | 2.07E-17 | 0.01 | 9.46E-01 | 0.00 | 1.00E+00 |
| FBgn0035006 | CG4563 | 11 | -0.02 | 8.74E-01 | 0.61 | 6.06E-02 | 0.00 | 1.00E+00 |
| FBgn0037844 | CG4570 | 35 | 1.56 | 1.77E-04 | 0.00 | 9.85E-01 | 0.01 | 1.00E+00 |
| FBgn0035016 | CG4612 | 3093 | -1.01 | 2.21E-13 | -0.03 | 7.29E-01 | 0.00 | 1.00E+00 |
| FBgn0031299 | CG4629 | 51 | -0.05 | 8.82E-01 | 0.67 | 2.58E-02 | 0.00 | 1.00E+00 |
| FBgn0030776 | CG4653 | 10 | -0.02 | 8.87E-01 | -1.22 | 2.33E-02 | -0.09 | 3.33E-01 |
| FBgn0038739 | CG4686 | 1110 | 0.67 | 8.46E-09 | -0.08 | 2.57E-01 | 0.00 | 1.00E+00 |
| FBgn0037992 | CG4702 | 49 | 0.17 | 5.07E-01 | -1.32 | 4.95E-04 | -0.36 | 5.98E-02 |
| FBgn0033820 | CG4716 | 80 | 0.10 | 4.21E-01 | -2.55 | 6.25E-03 | 0.02 | 9.19E-01 |
| FBgn0030790 | CG4768 | 1064 | -0.63 | 3.80E-07 | -0.05 | 5.59E-01 | 0.00 | 1.00E+00 |
| FBgn0030792 | CG4789 | 1285 | 0.59 | 2.29E-05 | -0.02 | 8.66E-01 | 0.00 | 1.00E+00 |
| FBgn0039013 | CG4813 | 494 | 0.59 | 4.01E-05 | -0.03 | 8.17E-01 | 0.01 | 1.00E+00 |
| FBgn0037999 | CG4860 | 453 | 0.67 | 2.87E-04 | -0.04 | 7.15E-01 | 0.00 | 1.00E+00 |
| FBgn0030803 | CG4880 | 886 | -0.73 | 1.11E-10 | -0.08 | 2.48E-01 | 0.00 | 1.00E+00 |
| FBgn0035959 | CG4911 | 926 | -0.72 | 6.09E-06 | 0.00 | 9.75E-01 | 0.00 | 1.00E+00 |
| FBgn0036587 | CG4950 | 63 | -0.37 | 1.42E-01 | -0.05 | 5.76E-01 | -1.06 | 3.07E-03 |
| FBgn0036579 | CG5027 | 367 | 0.81 | 2.00E-08 | -0.02 | 8.81E-01 | -0.01 | 1.00E+00 |
| FBgn0036576 | CG5151 | 885 | -0.67 | 4.99E-04 | -0.08 | 3.45E-01 | 0.00 | 1.00E+00 |
| FBgn0038943 | CG5391 | 400 | -0.78 | 3.04E-03 | 3.62 | 7.63E-72 | 0.01 | 9.15E-01 |
| FBgn0032429 | CG5446 | 1410 | 0.62 | 1.05E-06 | 0.00 | 9.99E-01 | 0.00 | 1.00E+00 |
| FBgn0039430 | CG5455 | 141 | -0.01 | 9.85E-01 | -0.87 | 5.09E-05 | -2.03 | 7.80E-21 |
| FBgn0036760 | CG5567 | 232 | 0.60 | 1.79E-03 | 0.46 | 6.67E-04 | 0.03 | 4.91E-01 |
| FBgn0036975 | CG5618 | 404 | 0.94 | 3.40E-05 | -0.02 | 8.64E-01 | 0.00 | 1.00E+00 |
| FBgn0036254 | CG5645 | 551 | -1.20 | 2.36E-10 | 0.36 | 8.57E-03 | 0.01 | 9.37E-01 |
| FBgn0037082 | CG5664 | 538 | -0.61 | 3.00E-05 | 0.46 | 6.71E-05 | 0.01 | 1.00E+00 |
| FBgn0032171 | CG5846 | 201 | 0.69 | 1.24E-03 | 0.03 | 7.79E-01 | 0.00 | 1.00E+00 |
| FBgn0025700 | CG5885 | 3676 | 0.60 | 7.64E-10 | -0.05 | 5.96E-01 | -0.01 | 1.00E+00 |
| FBgn0038400 | CG5903 | 2203 | 0.58 | 1.78E-07 | 0.13 | 7.05E-02 | 0.00 | 1.00E+00 |
| FBgn0036997 | CG5955 | 8 | -0.23 | 1.55E-01 | 3.40 | 5.82E-05 | -0.01 | 1.00E+00 |
| FBgn0034725 | CG6044 | 626 | 0.16 | 3.62E-01 | 0.96 | 1.39E-17 | -0.01 | 1.00E+00 |
| FBgn0031918 | CG6055 | 37 | 0.14 | 5.96E-01 | -1.12 | 5.37E-03 | -0.01 | 1.00E+00 |
| FBgn0036186 | CG6071 | 137 | -0.03 | 9.16E-01 | 0.73 | 1.03E-03 | 0.83 | 7.55E-04 |
| FBgn0032453 | CG6180 | 4013 | 0.63 | 5.57E-09 | -0.03 | 8.04E-01 | 0.00 | 1.00E+00 |
| FBgn0032343 | CG6201 | 37 | -0.25 | 3.33E-01 | -0.81 | 1.07E-02 | -0.01 | 1.00E+00 |
| FBgn0038321 | CG6218 | 206 | -0.62 | 1.72E-03 | -0.04 | 6.35E-01 | -0.02 | 7.96E-01 |
| FBgn0033866 | CG6280 | 188 | -0.32 | 9.19E-02 | -0.99 | 2.70E-08 | -0.01 | 1.00E+00 |
| FBgn0037807 | CG6293 | 15 | -0.15 | 5.13E-01 | -1.83 | 4.48E-03 | -0.02 | 7.77E-01 |
| FBgn0036121 | CG6310 | 302 | 0.31 | 1.04E-01 | -0.53 | 1.07E-04 | -1.17 | 6.76E-19 |
| FBgn0039464 | CG6330 | 158 | 0.41 | 8.87E-02 | 0.88 | 6.45E-04 | -0.20 | 1.22E-01 |
| FBgn0033875 | CG6357 | 463 | -0.71 | 3.51E-04 | -1.63 | 2.20E-19 | 0.00 | 1.00E+00 |
| FBgn0029690 | CG6414 | 96 | 0.63 | 1.99E-02 | -1.28 | 1.82E-03 | 0.00 | 1.00E+00 |
| FBgn0036702 | CG6512 | 2297 | 0.13 | 2.43E-01 | 0.63 | 1.96E-19 | 0.01 | 8.51E-01 |
| FBgn0032388 | CG6686 | 1995 | -0.61 | 1.94E-05 | 0.22 | 2.61E-02 | 0.00 | 1.00E+00 |
| FBgn0032305 | CG6700 | 2874 | -0.81 | 6.27E-07 | 0.05 | 5.82E-01 | 0.00 | 1.00E+00 |
| FBgn0037915 | CG6790 | 9 | 0.18 | 2.26E-01 | 3.70 | 8.75E-06 | 0.00 | 1.00E+00 |
| FBgn0038290 | CG6912 | 99 | -5.89 | 1.77E-07 | 4.27 | 3.46E-14 | 4.50 | 3.37E-15 |
| FBgn0036240 | CG6928 | 2325 | -0.21 | 2.85E-01 | 3.92 | 1.78E-156 | 2.87 | 1.32E-82 |
| FBgn0038972 | CG7054 | 111 | -0.57 | 2.87E-02 | 2.11 | 1.30E-26 | 0.33 | 4.67E-02 |

| | | | | | | | | |
|---|---|---|---|---|---|---|---|---|
| FBgn0038941 | CG7080 | 40 | 0.15 | 4.58E-01 | 4.03 | 3.25E-20 | 0.00 | 1.00E+00 |
| FBgn0037099 | CG7173 | 192 | 0.15 | 5.00E-01 | -0.65 | 5.38E-04 | -0.52 | 8.92E-03 |
| FBgn0035861 | CG7213 | 27 | 0.29 | 2.45E-01 | 7.51 | 2.14E-09 | 0.00 | 1.00E+00 |
| FBgn0031976 | CG7367 | 156 | -0.70 | 2.69E-03 | -0.17 | 1.49E-01 | -0.02 | 7.39E-01 |
| FBgn0037135 | CG7414 | 11131 | 0.59 | 7.82E-19 | 0.01 | 9.52E-01 | 0.00 | 1.00E+00 |
| FBgn0035833 | CG7565 | 175 | -0.69 | 2.66E-03 | -2.28 | 1.43E-19 | -0.04 | 5.16E-01 |
| FBgn0040793 | CG7630 | 1562 | 1.04 | 1.98E-14 | 0.05 | 5.86E-01 | 0.00 | 1.00E+00 |
| FBgn0033548 | CG7637 | 973 | 1.92 | 5.72E-24 | 0.02 | 8.99E-01 | 0.00 | 1.00E+00 |
| FBgn0036926 | CG7646 | 134 | -1.04 | 3.22E-05 | 0.02 | 8.47E-01 | 0.00 | 1.00E+00 |
| FBgn0036929 | CG7668 | 1484 | -0.86 | 1.89E-13 | -0.02 | 8.26E-01 | -0.01 | 9.40E-01 |
| FBgn0038619 | CG7685 | 164 | 0.63 | 6.57E-03 | 0.04 | 6.43E-01 | 0.00 | 1.00E+00 |
| FBgn0036714 | CG7692 | 656 | -0.87 | 8.80E-12 | 0.04 | 6.82E-01 | 0.03 | 5.49E-01 |
| FBgn0036509 | CG7739 | 1153 | -0.64 | 3.23E-11 | -0.05 | 6.08E-01 | -0.01 | 1.00E+00 |
| FBgn0036124 | CG7839 | 1429 | -0.80 | 6.12E-07 | 0.09 | 2.63E-01 | 0.00 | 1.00E+00 |
| FBgn0037548 | CG7900 | 41 | 4.55 | 2.26E-05 | 2.11 | 6.72E-04 | 0.00 | 1.00E+00 |
| FBgn0028534 | CG7916 | 16 | 0.13 | 3.16E-01 | 0.04 | 7.30E-01 | -1.26 | 2.16E-02 |
| FBgn0039737 | CG7920 | 3784 | 0.65 | 1.36E-09 | -0.26 | 2.06E-03 | 0.00 | 1.00E+00 |
| FBgn0028533 | CG7953 | 31 | -0.07 | 5.70E-01 | -3.18 | 4.57E-04 | -0.52 | 5.46E-02 |
| FBgn0038115 | CG7966 | 139 | 0.86 | 3.03E-03 | -0.07 | 4.85E-01 | 0.01 | 1.00E+00 |
| FBgn0035260 | CG7991 | 93 | 0.16 | 5.45E-01 | -1.20 | 1.07E-04 | 0.01 | 1.00E+00 |
| FBgn0037612 | CG8112 | 68 | 0.33 | 1.87E-01 | -0.92 | 1.04E-02 | -0.03 | 7.02E-01 |
| FBgn0034010 | CG8157 | 91 | 0.13 | 5.33E-01 | -3.51 | 4.35E-08 | 0.00 | 1.00E+00 |
| FBgn0034011 | CG8160 | 19 | 0.19 | 1.39E-01 | -0.66 | 5.32E-02 | 0.00 | 1.00E+00 |
| FBgn0034030 | CG8192 | 167 | 0.76 | 5.57E-04 | -0.26 | 6.59E-02 | 0.00 | 1.00E+00 |
| FBgn0034033 | CG8204 | 166 | 0.65 | 4.51E-03 | 0.01 | 9.09E-01 | 0.00 | 1.00E+00 |
| FBgn0034143 | CG8303 | 208 | 0.46 | 3.10E-02 | 0.69 | 2.01E-04 | 0.00 | 1.00E+00 |
| FBgn0032001 | CG8360 | 132 | 0.71 | 2.84E-03 | 0.03 | 7.61E-01 | 0.00 | 1.00E+00 |
| FBgn0034067 | CG8399 | 2181 | 0.73 | 4.88E-15 | 0.00 | 9.75E-01 | 0.00 | 1.00E+00 |
| FBgn0037754 | CG8500 | 73 | -0.69 | 8.60E-03 | -1.78 | 1.52E-04 | -0.03 | 6.54E-01 |
| FBgn0035791 | CG8539 | 9 | -0.06 | 7.27E-01 | 0.83 | 2.63E-02 | 0.00 | 1.00E+00 |
| FBgn0035714 | CG8549 | 532 | -0.44 | 1.74E-02 | 0.60 | 6.84E-05 | 0.00 | 1.00E+00 |
| FBgn0036386 | CG8833 | 390 | -0.60 | 2.98E-04 | 0.03 | 8.06E-01 | 0.01 | 9.13E-01 |
| FBgn0038404 | CG8925 | 35 | -0.18 | 4.46E-01 | 0.59 | 3.32E-02 | 0.00 | 1.00E+00 |
| FBgn0038405 | CG8927 | 52 | -0.12 | 6.41E-01 | 2.02 | 1.81E-04 | 0.74 | 2.16E-02 |
| FBgn0028920 | CG8997 | 28 | -0.10 | 5.51E-01 | -5.54 | 2.35E-06 | -0.77 | 3.60E-02 |
| FBgn0040931 | CG9034 | 454 | 1.08 | 6.35E-06 | 0.05 | 5.91E-01 | 0.00 | 1.00E+00 |
| FBgn0030610 | CG9065 | 223 | 1.09 | 7.84E-06 | 0.38 | 2.24E-02 | 0.00 | 1.00E+00 |
| FBgn0030716 | CG9170 | 1167 | 0.00 | 9.77E-01 | 0.67 | 9.06E-26 | 0.01 | 9.27E-01 |
| FBgn0035181 | CG9205 | 889 | 1.09 | 7.19E-17 | 0.07 | 3.38E-01 | 0.00 | 1.00E+00 |
| FBgn0030669 | CG9240 | 519 | 0.58 | 4.27E-05 | 0.07 | 3.97E-01 | 0.00 | 1.00E+00 |
| FBgn0032925 | CG9246 | 1988 | -0.61 | 5.99E-06 | 0.06 | 4.73E-01 | 0.00 | 1.00E+00 |
| FBgn0038179 | CG9312 | 150 | 0.53 | 2.53E-02 | -0.70 | 4.84E-03 | -0.21 | 1.14E-01 |
| FBgn0032889 | CG9331 | 2348 | 0.64 | 4.33E-13 | -0.05 | 5.25E-01 | -0.07 | 1.62E-01 |
| FBgn0032897 | CG9336 | 285 | 1.72 | 2.22E-08 | -0.01 | 8.95E-01 | 0.01 | 1.00E+00 |
| FBgn0032899 | CG9338 | 208 | 1.66 | 1.58E-07 | -0.12 | 2.34E-01 | 0.00 | 1.00E+00 |
| FBgn0030569 | CG9411 | 168 | -0.21 | 4.11E-01 | 0.90 | 2.91E-04 | 0.00 | 1.00E+00 |
| FBgn0037721 | CG9427 | 1394 | 0.88 | 9.72E-07 | 0.00 | 9.73E-01 | 0.00 | 1.00E+00 |
| FBgn0037749 | CG9471 | 643 | 0.73 | 3.50E-06 | 0.10 | 2.20E-01 | 0.00 | 1.00E+00 |
| FBgn0030588 | CG9521 | 33 | -0.09 | 7.33E-01 | 1.59 | 1.95E-04 | 0.33 | 7.06E-02 |
| FBgn0031824 | CG9547 | 243 | 0.77 | 3.20E-04 | 0.04 | 7.14E-01 | 0.00 | 1.00E+00 |
| FBgn0034184 | CG9646 | 708 | -0.15 | 4.09E-01 | 0.62 | 8.27E-08 | 0.00 | 1.00E+00 |
| FBgn0029939 | CG9650 | 337 | -0.32 | 1.09E-01 | -0.61 | 9.83E-04 | 0.00 | 1.00E+00 |
| FBgn0030160 | CG9691 | 853 | 0.65 | 4.41E-04 | 0.03 | 7.54E-01 | -0.01 | 1.00E+00 |
| FBgn0039754 | CG9747 | 717 | 1.09 | 4.41E-12 | -0.51 | 1.11E-04 | -0.01 | 1.00E+00 |
| FBgn0024248 | chico | 1054 | -0.76 | 1.68E-13 | -0.08 | 2.97E-01 | 0.01 | 1.00E+00 |
| FBgn0000307 | chif | 2908 | -0.58 | 8.33E-04 | 0.04 | 6.23E-01 | 0.00 | 1.00E+00 |
| FBgn0086758 | chinmo | 519 | 0.32 | 1.69E-01 | 0.72 | 1.56E-03 | 0.35 | 4.97E-02 |
| FBgn0045761 | CHKov1 | 1347 | -0.44 | 8.28E-04 | 0.59 | 8.77E-09 | 0.00 | 1.00E+00 |
| FBgn0039328 | CHKov2 | 325 | -0.17 | 4.44E-01 | 1.30 | 1.30E-19 | 0.00 | 1.00E+00 |
| FBgn0028387 | chm | 727 | -0.61 | 1.26E-07 | -0.03 | 7.48E-01 | 0.00 | 1.00E+00 |
| FBgn0035589 | CHMP2B | 224 | 0.81 | 1.72E-04 | 0.03 | 7.93E-01 | 0.00 | 1.00E+00 |
| FBgn0043002 | Chrac-14 | 373 | 0.65 | 1.59E-03 | 0.07 | 4.13E-01 | 0.00 | 1.00E+00 |
| FBgn0036165 | chrb | 3380 | -0.90 | 1.39E-05 | -0.05 | 5.80E-01 | -0.01 | 1.00E+00 |
| FBgn0022702 | Cht2 | 3503 | 0.66 | 5.55E-09 | -0.06 | 4.38E-01 | -0.01 | 8.73E-01 |
| FBgn0038180 | Cht5 | 1359 | 0.60 | 1.59E-03 | 0.04 | 6.63E-01 | 0.00 | 1.00E+00 |
| FBgn0035398 | Cht7 | 1066 | 0.58 | 3.91E-03 | -0.05 | 5.99E-01 | 0.01 | 9.27E-01 |
| FBgn0004859 | ci | 8944 | -1.01 | 3.99E-10 | -0.05 | 5.37E-01 | 0.00 | 1.00E+00 |
| FBgn0001977 | CIAPIN1 | 1526 | 0.20 | 1.04E-01 | 0.71 | 1.61E-18 | 0.00 | 1.00E+00 |
| FBgn0026084 | cib | 15779 | -0.64 | 2.76E-06 | 0.06 | 4.84E-01 | -0.01 | 1.00E+00 |
| FBgn0015024 | CkIalpha | 6805 | -1.20 | 3.97E-18 | -0.03 | 7.82E-01 | 0.00 | 1.00E+00 |
| FBgn0000259 | CkIIbeta | 4366 | -0.71 | 2.16E-16 | 0.01 | 9.62E-01 | 0.00 | 1.00E+00 |
| FBgn0000318 | cl | 1966 | 1.07 | 9.57E-13 | 0.11 | 1.85E-01 | -0.01 | 1.00E+00 |
| FBgn0052251 | Claspin | 1098 | -0.97 | 1.62E-05 | 0.02 | 8.37E-01 | 0.00 | 1.00E+00 |
| FBgn0259152 | Clbn | 1163 | -0.65 | 2.13E-07 | 0.52 | 2.71E-08 | 0.00 | 1.00E+00 |
| FBgn0020503 | CLIP-190 | 1351 | -0.94 | 5.34E-09 | 0.01 | 9.54E-01 | 0.00 | 1.00E+00 |
| FBgn0035767 | Cln7 | 1086 | 0.63 | 1.01E-04 | -0.09 | 2.90E-01 | 0.00 | 1.00E+00 |
| FBgn0026255 | clumsy | 768 | -0.05 | 8.43E-01 | 1.88 | 6.29E-37 | 0.10 | 1.79E-01 |
| FBgn0034802 | CNBP | 42330 | -0.99 | 2.65E-17 | 0.03 | 7.89E-01 | 0.00 | 1.00E+00 |
| FBgn0264077 | Cnx14D | 13 | 0.08 | 7.33E-01 | 6.73 | 7.83E-07 | 0.00 | 1.00E+00 |

| | | | | | | | | |
|---|---|---|---|---|---|---|---|---|
| FBgn0010105 | comm | 299 | 0.84 | 5.61E-06 | -0.03 | 7.35E-01 | 0.00 | 1.00E+00 |
| FBgn0039994 | conu | 1969 | -0.92 | 1.92E-15 | 0.03 | 7.39E-01 | 0.00 | 1.00E+00 |
| FBgn0033192 | Corin | 24 | -0.10 | 7.04E-01 | -1.49 | 7.09E-03 | 0.00 | 1.00E+00 |
| FBgn0030028 | Corp | 23 | 0.18 | 4.74E-01 | 0.74 | 1.54E-02 | -0.01 | 1.00E+00 |
| FBgn0032833 | COX4 | 2941 | 0.86 | 5.76E-08 | 0.12 | 1.56E-01 | 0.00 | 1.00E+00 |
| FBgn0019624 | COX5A | 2606 | 1.17 | 1.83E-09 | 0.05 | 5.56E-01 | 0.00 | 1.00E+00 |
| FBgn0031830 | COX5B | 1591 | 0.59 | 1.91E-06 | 0.00 | 9.88E-01 | -0.02 | 7.52E-01 |
| FBgn0031066 | COX6B | 2997 | 0.78 | 3.12E-11 | -0.06 | 5.05E-01 | -0.01 | 1.00E+00 |
| FBgn0040529 | COX7A | 3160 | 1.11 | 4.94E-14 | -0.01 | 9.48E-01 | 0.00 | 1.00E+00 |
| FBgn0263911 | COX8 | 1195 | 0.59 | 1.97E-04 | 0.02 | 8.62E-01 | 0.00 | 1.00E+00 |
| FBgn0053302 | Cpr31A | 46 | 1.71 | 3.35E-06 | 0.02 | 8.71E-01 | 0.00 | 1.00E+00 |
| FBgn0028871 | Cpr35B | 60 | 1.28 | 2.13E-04 | 0.00 | 9.89E-01 | -0.01 | 1.00E+00 |
| FBgn0033598 | Cpr47Eb | 67 | 1.16 | 1.59E-03 | 0.05 | 6.06E-01 | 0.00 | 1.00E+00 |
| FBgn0033602 | Cpr47Ee | 24 | -0.03 | 8.88E-01 | -1.69 | 4.84E-03 | 0.00 | 1.00E+00 |
| FBgn0033730 | Cpr49Ag | 381 | 0.92 | 1.89E-07 | -0.07 | 3.83E-01 | 0.00 | 1.00E+00 |
| FBgn0033731 | Cpr49Ah | 539 | 0.81 | 1.36E-03 | 1.02 | 1.13E-04 | 0.16 | 1.61E-01 |
| FBgn0033942 | Cpr51A | 266 | 0.96 | 8.10E-05 | -0.51 | 9.48E-03 | 0.00 | 1.00E+00 |
| FBgn0034499 | Cpr56F | 33 | -0.17 | 4.91E-01 | -2.07 | 2.64E-04 | -0.01 | 1.00E+00 |
| FBgn0035737 | Cpr65Ec | 171 | 0.83 | 1.01E-03 | -0.33 | 4.57E-02 | -0.21 | 1.14E-01 |
| FBgn0052029 | Cpr66D | 478 | 1.87 | 5.31E-25 | -0.03 | 8.00E-01 | 0.00 | 1.00E+00 |
| FBgn0036109 | Cpr67Fa2 | 86 | 0.27 | 2.67E-01 | -0.09 | 3.18E-01 | -0.66 | 5.64E-03 |
| FBgn0036617 | Cpr72Ea | 13 | 0.83 | 6.43E-03 | 0.03 | 7.54E-01 | 0.00 | 1.00E+00 |
| FBgn0037114 | Cpr78E | 355 | 0.71 | 9.83E-03 | -0.59 | 2.57E-02 | -0.02 | 8.40E-01 |
| FBgn0042701 | CR12628 | 15 | 0.32 | 1.58E-01 | 1.70 | 3.97E-03 | 0.03 | 7.10E-01 |
| FBgn0040928 | CR15345 | 118 | -0.66 | 4.31E-03 | 0.00 | 9.77E-01 | 0.00 | 1.00E+00 |
| FBgn0262532 | CR43086 | 36 | 0.01 | 9.46E-01 | 5.51 | 5.98E-22 | 3.08 | 3.23E-06 |
| FBgn0263093 | CR43361 | 426 | -0.03 | 8.89E-01 | -1.74 | 1.53E-30 | 0.00 | 1.00E+00 |
| FBgn0263744 | CR43669 | 21 | -0.86 | 5.34E-03 | 0.04 | 7.25E-01 | 0.00 | 1.00E+00 |
| FBgn0035636 | Cralbp | 80 | 0.73 | 3.28E-03 | -0.10 | 3.25E-01 | 0.00 | 1.00E+00 |
| FBgn0004396 | CrebA | 1973 | -0.77 | 1.03E-05 | 0.00 | 9.83E-01 | 0.00 | 1.00E+00 |
| FBgn0023023 | CRMP | 259 | -0.13 | 5.66E-01 | -1.04 | 7.11E-10 | 0.00 | 1.00E+00 |
| FBgn0015924 | crq | 883 | -0.78 | 1.85E-12 | -0.06 | 4.91E-01 | 0.00 | 1.00E+00 |
| FBgn0036746 | Crtc | 1137 | -0.88 | 5.35E-06 | 0.04 | 7.12E-01 | 0.00 | 1.00E+00 |
| FBgn0000384 | cta | 945 | -0.64 | 6.04E-04 | 0.02 | 8.88E-01 | 0.00 | 1.00E+00 |
| FBgn0062412 | Ctr1B | 34 | 0.02 | 8.78E-01 | -0.82 | 3.92E-02 | -0.07 | 4.03E-01 |
| FBgn0260932 | cuff | 181 | -0.59 | 9.91E-04 | 0.34 | 1.92E-02 | 0.00 | 1.00E+00 |
| FBgn0032956 | Cul2 | 6827 | -0.48 | 5.97E-05 | 3.49 | 0.00E+00 | 0.01 | 1.00E+00 |
| FBgn0261268 | Cul3 | 3133 | -0.62 | 1.96E-10 | -0.02 | 9.07E-01 | 0.00 | 1.00E+00 |
| FBgn0035880 | Culd | 2104 | -0.15 | 5.49E-01 | 3.94 | 7.19E-150 | 5.04 | 1.82E-247 |
| FBgn0031689 | Cyp28d1 | 91 | 1.86 | 1.03E-03 | -0.16 | 2.56E-01 | 0.00 | 1.00E+00 |
| FBgn0001992 | Cyp303a1 | 68 | -0.99 | 1.46E-03 | 0.02 | 8.88E-01 | 0.02 | 9.40E-01 |
| FBgn0034756 | Cyp6d2 | 381 | 0.01 | 9.79E-01 | 4.75 | 2.83E-216 | 0.02 | 7.02E-01 |
| FBgn0039006 | Cyp6d4 | 137 | 0.36 | 1.38E-01 | 1.42 | 6.67E-10 | 0.00 | 1.00E+00 |
| FBgn0025454 | Cyp6g1 | 147 | 0.24 | 3.07E-01 | -1.83 | 1.59E-04 | -0.01 | 1.00E+00 |
| FBgn0031126 | Cyp6v1 | 1037 | -1.19 | 1.46E-27 | -0.18 | 5.37E-03 | 0.00 | 1.00E+00 |
| FBgn0000411 | D | 14 | 0.19 | 2.73E-01 | -0.62 | 4.04E-02 | 0.01 | 1.00E+00 |
| FBgn0005677 | dac | 806 | -0.13 | 4.87E-01 | -0.95 | 7.73E-13 | 0.03 | 5.06E-01 |
| FBgn0263852 | Dad1 | 1589 | 1.13 | 4.02E-14 | 0.00 | 9.86E-01 | -0.01 | 1.00E+00 |
| FBgn0263930 | dally | 6391 | -0.67 | 2.37E-07 | 0.06 | 4.68E-01 | 0.00 | 1.00E+00 |
| FBgn0262636 | dati | 43 | -0.07 | 7.98E-01 | 3.80 | 3.07E-10 | 5.11 | 4.48E-18 |
| FBgn0031820 | Daxx | 2065 | -0.59 | 3.50E-06 | -0.02 | 8.83E-01 | 0.01 | 1.00E+00 |
| FBgn0067779 | dbr | 1696 | -0.84 | 1.08E-11 | 0.01 | 9.73E-01 | 0.00 | 1.00E+00 |
| FBgn0002413 | dco | 3330 | -0.72 | 1.41E-14 | -0.01 | 9.24E-01 | 0.00 | 1.00E+00 |
| FBgn0036534 | DCP2 | 6445 | -0.99 | 5.36E-08 | 0.03 | 7.18E-01 | 0.00 | 1.00E+00 |
| FBgn0029067 | Dd | 965 | -0.87 | 1.81E-10 | -0.02 | 8.60E-01 | 0.00 | 1.00E+00 |
| FBgn0086251 | del | 642 | -0.93 | 2.75E-08 | -0.05 | 5.91E-01 | 0.00 | 1.00E+00 |
| FBgn0267972 | Der-1 | 1024 | 0.64 | 1.65E-07 | 0.03 | 7.60E-01 | 0.00 | 1.00E+00 |
| FBgn0022893 | Df31 | 48354 | -0.63 | 6.61E-10 | 0.02 | 8.64E-01 | 0.00 | 1.00E+00 |
| FBgn0013812 | Dhc93AB | 47 | -0.10 | 7.32E-01 | 0.83 | 4.04E-03 | 0.00 | 1.00E+00 |
| FBgn0000449 | dib | 86 | -0.60 | 1.66E-02 | -1.67 | 3.76E-06 | -0.04 | 5.19E-01 |
| FBgn0011274 | Dif | 323 | -0.45 | 3.54E-02 | 2.57 | 3.24E-38 | 0.00 | 1.00E+00 |
| FBgn0000459 | disco | 168 | -0.02 | 9.57E-01 | -0.73 | 2.21E-03 | 0.02 | 8.78E-01 |
| FBgn0285879 | disco-r | 318 | 0.11 | 6.79E-01 | -1.45 | 2.90E-11 | 0.04 | 4.80E-01 |
| FBgn0263106 | DnaJ-1 | 4644 | 0.13 | 5.57E-01 | -0.72 | 2.47E-05 | 0.00 | 1.00E+00 |
| FBgn0286075 | DNAlig3 | 384 | -0.23 | 1.73E-01 | 0.73 | 5.92E-14 | 0.00 | 1.00E+00 |
| FBgn0030506 | DNAlig4 | 262 | -0.19 | 3.68E-01 | 0.74 | 2.66E-08 | 0.00 | 1.00E+00 |
| FBgn0002891 | DNApol-zeta | 1022 | -0.31 | 3.75E-02 | 0.88 | 1.75E-15 | 0.02 | 6.30E-01 |
| FBgn0035542 | DOR | 485 | -0.62 | 6.59E-03 | 0.00 | 9.77E-01 | 0.00 | 1.00E+00 |
| FBgn0024558 | Dph5 | 1749 | 0.65 | 2.00E-07 | 0.09 | 2.15E-01 | 0.00 | 1.00E+00 |
| FBgn0037295 | dpr16 | 116 | 2.16 | 3.85E-14 | 0.00 | 9.71E-01 | -0.01 | 9.39E-01 |
| FBgn0053196 | dpy | 6720 | 1.12 | 5.51E-07 | 0.63 | 1.77E-03 | 0.01 | 1.00E+00 |
| FBgn0032293 | Dpy-30L1 | 556 | 0.88 | 3.69E-06 | -0.01 | 9.49E-01 | 0.00 | 1.00E+00 |
| FBgn0260006 | drd | 87 | 0.15 | 5.72E-01 | 1.73 | 4.49E-05 | 0.01 | 1.00E+00 |
| FBgn0004638 | drk | 3220 | -0.61 | 7.50E-08 | -0.02 | 8.51E-01 | 0.00 | 1.00E+00 |
| FBgn0024244 | drm | 2325 | -0.71 | 1.13E-07 | -0.15 | 8.06E-02 | -0.01 | 1.00E+00 |
| FBgn0278608 | Dsp1 | 6983 | -0.58 | 7.34E-05 | 0.00 | 9.82E-01 | 0.00 | 1.00E+00 |
| FBgn0261799 | dsx-c73A | 101 | 1.06 | 6.38E-06 | 0.37 | 2.38E-02 | 0.00 | 1.00E+00 |
| FBgn0027101 | Dyrk3 | 2604 | -0.88 | 1.38E-09 | -0.03 | 7.75E-01 | 0.00 | 1.00E+00 |
| FBgn0002592 | E(spl)m2-BFM | 1109 | 0.65 | 6.60E-06 | -0.12 | 1.52E-01 | 0.00 | 1.00E+00 |

| | | | | | | | | |
|---|---|---|---|---|---|---|---|---|
| FBgn0002629 | E(spl)m4-BFM | 800 | 0.72 | 3.01E-07 | 0.05 | 5.64E-01 | 0.09 | 1.63E-01 |
| FBgn0002732 | E(spl)malpha-BFM | 2631 | 0.76 | 1.39E-05 | -0.01 | 9.10E-01 | 0.00 | 1.00E+00 |
| FBgn0002735 | E(spl)mgamma-HLH | 148 | 0.65 | 4.39E-03 | 0.05 | 5.80E-01 | 0.01 | 8.87E-01 |
| FBgn0011766 | E2f1 | 4380 | -0.84 | 3.21E-10 | 0.01 | 9.30E-01 | 0.01 | 1.00E+00 |
| FBgn0069242 | eca | 731 | 0.68 | 1.04E-03 | 0.00 | 9.98E-01 | -0.01 | 1.00E+00 |
| FBgn0033879 | Echs1 | 3518 | 0.92 | 3.48E-26 | -0.02 | 8.64E-01 | 0.00 | 1.00E+00 |
| FBgn0260746 | Ect3 | 339 | -1.92 | 2.07E-19 | 0.04 | 7.15E-01 | -0.01 | 1.00E+00 |
| FBgn0028737 | eEF1beta | 17684 | 0.95 | 5.64E-23 | -0.07 | 3.37E-01 | 0.00 | 1.00E+00 |
| FBgn0029176 | eEF1gamma | 47067 | 0.69 | 1.56E-30 | -0.04 | 6.50E-01 | 0.00 | 1.00E+00 |
| FBgn0034883 | Eglp2 | 10 | 0.03 | 8.59E-01 | 1.95 | 2.90E-03 | 0.19 | 1.44E-01 |
| FBgn0037270 | eIF3f1 | 6584 | 0.80 | 1.53E-14 | -0.04 | 7.25E-01 | 0.00 | 1.00E+00 |
| FBgn0022023 | eIF3h | 7150 | 0.90 | 6.93E-21 | 0.00 | 9.80E-01 | 0.00 | 1.00E+00 |
| FBgn0020660 | eIF4B | 389 | -0.77 | 7.87E-03 | -0.12 | 2.19E-01 | -0.07 | 3.37E-01 |
| FBgn0053100 | eIF4EHP | 1127 | -1.07 | 3.14E-18 | 0.03 | 7.43E-01 | 0.00 | 1.00E+00 |
| FBgn0023213 | eIF4G1 | 11399 | -0.77 | 1.42E-10 | -0.03 | 7.88E-01 | 0.00 | 1.00E+00 |
| FBgn0262734 | eIF4H1 | 7393 | -1.47 | 7.70E-14 | 0.01 | 9.18E-01 | 0.00 | 1.00E+00 |
| FBgn0030719 | eIF5 | 87 | -2.37 | 1.38E-05 | 0.03 | 7.39E-01 | 0.00 | 1.00E+00 |
| FBgn0034915 | eIF6 | 2129 | 0.36 | 8.36E-03 | 0.83 | 2.03E-15 | 0.00 | 1.00E+00 |
| FBgn0004858 | elB | 1567 | -0.77 | 1.52E-09 | 0.04 | 6.25E-01 | 0.00 | 1.00E+00 |
| FBgn0037534 | ELOVL | 41 | 0.60 | 2.45E-02 | -0.09 | 4.12E-01 | 0.00 | 1.00E+00 |
| FBgn0062440 | EMRE | 456 | 0.87 | 1.66E-05 | 0.01 | 9.36E-01 | 0.00 | 1.00E+00 |
| FBgn0000579 | Eno | 10736 | 0.65 | 1.05E-11 | -0.03 | 7.79E-01 | -0.01 | 1.00E+00 |
| FBgn0264693 | ens | 3968 | -0.63 | 2.15E-06 | 0.01 | 9.18E-01 | 0.00 | 1.00E+00 |
| FBgn0036319 | Ent3 | 19 | -0.14 | 4.83E-01 | -0.83 | 1.87E-02 | -0.01 | 1.00E+00 |
| FBgn0025936 | Eph | 4729 | -0.81 | 1.92E-08 | -0.07 | 4.30E-01 | 0.00 | 1.00E+00 |
| FBgn0040324 | Ephrin | 2393 | -0.62 | 2.73E-03 | -0.01 | 9.53E-01 | 0.00 | 1.00E+00 |
| FBgn0005660 | Ets21C | 87 | 0.15 | 5.10E-01 | 2.51 | 3.68E-04 | 0.00 | 1.00E+00 |
| FBgn0033668 | exp | 166 | -0.65 | 1.72E-03 | -0.06 | 4.78E-01 | 0.00 | 1.00E+00 |
| FBgn0034529 | FAM21 | 494 | -0.58 | 1.42E-05 | 0.02 | 8.90E-01 | 0.01 | 1.00E+00 |
| FBgn0032428 | Fam92 | 20 | 0.07 | 7.33E-01 | 3.79 | 5.35E-11 | 0.00 | 1.00E+00 |
| FBgn0042627 | FASN2 | 15 | 0.06 | 6.87E-01 | 0.67 | 4.83E-02 | 0.10 | 2.91E-01 |
| FBgn0034999 | Fatp3 | 93 | 0.25 | 3.03E-01 | -1.49 | 1.09E-03 | -0.09 | 3.00E-01 |
| FBgn0014163 | fax | 30703 | -0.92 | 3.44E-16 | -0.10 | 1.77E-01 | 0.00 | 1.00E+00 |
| FBgn0000639 | Fbp1 | 20462 | 0.58 | 1.65E-02 | -0.05 | 6.80E-01 | 0.00 | 1.00E+00 |
| FBgn0004897 | fd96Ca | 39 | 0.18 | 2.26E-01 | -2.52 | 9.80E-08 | 0.00 | 1.00E+00 |
| FBgn0004898 | fd96Cb | 154 | 1.83 | 1.09E-03 | -1.96 | 8.37E-05 | 0.01 | 1.00E+00 |
| FBgn0011768 | Fdh | 3119 | 1.45 | 1.89E-32 | 0.12 | 9.98E-02 | 0.02 | 6.08E-01 |
| FBgn0015221 | Fer2LCH | 4994 | 0.62 | 1.05E-16 | 0.11 | 6.21E-02 | 0.00 | 1.00E+00 |
| FBgn0039969 | Fis1 | 1106 | -0.59 | 9.95E-05 | -0.02 | 8.95E-01 | 0.00 | 1.00E+00 |
| FBgn0000658 | fj | 5209 | -0.61 | 3.71E-11 | 0.11 | 9.65E-02 | 0.00 | 1.00E+00 |
| FBgn0013954 | Fkbp12 | 9074 | 1.03 | 6.70E-36 | 0.08 | 2.51E-01 | 0.00 | 1.00E+00 |
| FBgn0033806 | FLASH | 452 | -0.84 | 2.02E-07 | 0.04 | 6.52E-01 | 0.00 | 1.00E+00 |
| FBgn0028734 | Fmr1 | 5727 | -0.78 | 5.68E-09 | 0.05 | 5.79E-01 | 0.00 | 1.00E+00 |
| FBgn0286222 | Fum1 | 4193 | 0.72 | 1.52E-10 | -0.04 | 7.12E-01 | 0.00 | 1.00E+00 |
| FBgn0039932 | fuss | 37 | -0.01 | 9.66E-01 | 2.22 | 1.05E-05 | 0.00 | 1.00E+00 |
| FBgn0001085 | fz | 2807 | -0.62 | 1.53E-06 | -0.08 | 3.18E-01 | 0.00 | 1.00E+00 |
| FBgn0031275 | GABA-B-R3 | 12 | 0.31 | 1.46E-01 | 3.04 | 4.22E-05 | 0.00 | 1.00E+00 |
| FBgn0033153 | Gadd45 | 924 | -0.03 | 8.97E-01 | 1.85 | 2.07E-41 | 0.00 | 1.00E+00 |
| FBgn0001122 | Galphao | 883 | -0.93 | 4.45E-10 | -0.02 | 8.19E-01 | 0.00 | 1.00E+00 |
| FBgn0004435 | Galphaq | 656 | -1.67 | 8.33E-15 | 0.00 | 9.86E-01 | 0.00 | 1.00E+00 |
| FBgn0001123 | Galphas | 2681 | -0.69 | 1.58E-16 | -0.03 | 8.08E-01 | 0.00 | 1.00E+00 |
| FBgn0001091 | Gapdh1 | 5452 | 0.68 | 3.17E-08 | 0.03 | 7.91E-01 | 0.00 | 1.00E+00 |
| FBgn0001092 | Gapdh2 | 16172 | 0.72 | 1.33E-21 | 0.08 | 2.03E-01 | 0.00 | 1.00E+00 |
| FBgn0032223 | GATAd | 1049 | -0.75 | 4.92E-07 | -0.05 | 6.16E-01 | 0.00 | 1.00E+00 |
| FBgn0001105 | Gbeta13F | 6116 | -1.18 | 2.14E-13 | -0.02 | 8.45E-01 | 0.00 | 1.00E+00 |
| FBgn0040319 | Gclc | 466 | -0.10 | 6.52E-01 | 0.86 | 6.23E-10 | 0.00 | 1.00E+00 |
| FBgn0020388 | Gcn5 | 1584 | 0.01 | 9.38E-01 | 1.05 | 8.06E-33 | 0.01 | 1.00E+00 |
| FBgn0000808 | gd | 21 | 0.13 | 5.32E-01 | 3.47 | 7.24E-13 | 0.00 | 1.00E+00 |
| FBgn0033081 | geminin | 1895 | 0.66 | 1.64E-06 | -0.02 | 9.04E-01 | 0.00 | 1.00E+00 |
| FBgn0250823 | gish | 4407 | -0.94 | 4.34E-13 | 0.03 | 7.82E-01 | 0.00 | 1.00E+00 |
| FBgn0015229 | glec | 1116 | -0.61 | 3.40E-05 | 0.02 | 8.88E-01 | -0.01 | 1.00E+00 |
| FBgn0004913 | Gnf1 | 1884 | -0.64 | 1.25E-06 | -0.01 | 9.54E-01 | 0.01 | 1.00E+00 |
| FBgn0263048 | Gpdh3 | 671 | -0.71 | 1.65E-07 | 0.05 | 5.72E-01 | 0.01 | 1.00E+00 |
| FBgn0260798 | Gprk1 | 1444 | -0.93 | 2.42E-16 | -0.06 | 4.44E-01 | 0.00 | 1.00E+00 |
| FBgn0035167 | Gr61a | 111 | 0.64 | 1.02E-02 | 1.25 | 3.97E-08 | 0.00 | 1.00E+00 |
| FBgn0039494 | grass | 168 | 0.95 | 6.23E-04 | -0.03 | 7.66E-01 | 0.00 | 1.00E+00 |
| FBgn0001137 | grk | 60 | -1.88 | 3.50E-06 | 0.01 | 9.09E-01 | -0.01 | 1.00E+00 |
| FBgn0019982 | Gs1l | 293 | 0.61 | 5.31E-03 | 0.00 | 9.95E-01 | 0.00 | 1.00E+00 |
| FBgn0001148 | gsb | 14 | 0.09 | 6.04E-01 | -1.84 | 2.89E-03 | 0.00 | 1.00E+00 |
| FBgn0001147 | gsb-n | 13 | 0.07 | 6.91E-01 | -1.55 | 4.35E-03 | 0.00 | 1.00E+00 |
| FBgn0001149 | GstD1 | 10655 | 1.18 | 7.26E-22 | 0.08 | 3.21E-01 | -0.01 | 1.00E+00 |
| FBgn0010039 | GstD3 | 959 | -1.17 | 1.90E-10 | 0.63 | 3.50E-07 | 0.00 | 1.00E+00 |
| FBgn0038020 | GstD9 | 723 | -0.53 | 5.19E-03 | 0.89 | 3.62E-10 | 0.02 | 7.52E-01 |
| FBgn0034335 | GstE1 | 1270 | 0.50 | 2.32E-02 | 5.48 | 0.00E+00 | 0.62 | 1.95E-05 |
| FBgn0027590 | GstE12 | 2243 | 0.80 | 1.39E-08 | 0.06 | 5.13E-01 | 0.00 | 1.00E+00 |
| FBgn0063498 | GstE2 | 76 | 0.03 | 8.52E-01 | 6.89 | 9.09E-39 | 0.00 | 1.00E+00 |
| FBgn0063495 | GstE5 | 41 | 0.36 | 1.44E-01 | 1.34 | 9.60E-05 | -0.01 | 1.00E+00 |
| FBgn0063494 | GstE6 | 374 | 0.19 | 4.42E-01 | 4.52 | 2.60E-102 | 0.80 | 1.67E-03 |
| FBgn0063492 | GstE8 | 106 | 0.14 | 5.85E-01 | 1.66 | 7.07E-11 | 0.63 | 1.24E-02 |

| | | | | | | | | |
|---|---|---|---|---|---|---|---|---|
| FBgn0050000 | GstT1 | 515 | -0.02 | 9.28E-01 | 1.05 | 3.81E-19 | 0.00 | 1.00E+00 |
| FBgn0031117 | GstT3 | 109 | 0.98 | 3.51E-04 | 0.02 | 8.80E-01 | -0.03 | 6.08E-01 |
| FBgn0026238 | gus | 2692 | -0.99 | 3.39E-18 | -0.01 | 9.44E-01 | 0.00 | 1.00E+00 |
| FBgn0039936 | Gyf | 5384 | -0.60 | 9.10E-09 | -0.05 | 5.59E-01 | 0.00 | 1.00E+00 |
| FBgn0016660 | H15 | 67 | 0.29 | 1.75E-01 | -1.81 | 1.80E-04 | 0.01 | 1.00E+00 |
| FBgn0032394 | Hacd1 | 670 | 0.71 | 1.68E-07 | -0.22 | 1.37E-02 | 0.00 | 1.00E+00 |
| FBgn0034488 | Hacl | 2789 | 0.59 | 1.23E-11 | -0.03 | 7.67E-01 | 0.00 | 1.00E+00 |
| FBgn0286508 | Had1 | 310 | 0.66 | 8.55E-03 | -0.01 | 9.16E-01 | 0.01 | 1.00E+00 |
| FBgn0033949 | Had2 | 63 | 0.21 | 4.30E-01 | -0.31 | 7.20E-02 | -0.59 | 2.15E-02 |
| FBgn0046706 | Haspin | 1098 | -1.00 | 3.56E-13 | 0.01 | 9.18E-01 | -0.01 | 9.13E-01 |
| FBgn0051119 | HDAC11 | 108 | -0.87 | 4.32E-04 | 0.13 | 3.37E-01 | -0.01 | 1.00E+00 |
| FBgn0011224 | heph | 19761 | -0.80 | 6.76E-07 | -0.01 | 9.33E-01 | 0.00 | 1.00E+00 |
| FBgn0001185 | her | 823 | -0.65 | 6.24E-10 | 0.00 | 9.90E-01 | 0.00 | 1.00E+00 |
| FBgn0031459 | HINT1 | 2526 | 1.22 | 1.99E-18 | 0.04 | 6.85E-01 | 0.00 | 1.00E+00 |
| FBgn0029676 | HIP-R | 1449 | -1.00 | 1.27E-11 | 0.02 | 8.38E-01 | -0.01 | 1.00E+00 |
| FBgn0010228 | HmgZ | 5046 | -1.13 | 3.34E-40 | -0.06 | 4.70E-01 | 0.00 | 1.00E+00 |
| FBgn0035160 | hng3 | 952 | 0.83 | 6.58E-08 | 0.03 | 7.51E-01 | -0.02 | 7.72E-01 |
| FBgn0286786 | hoip | 2078 | 0.67 | 3.30E-05 | 0.06 | 4.66E-01 | 0.00 | 1.00E+00 |
| FBgn0025777 | homer | 1227 | -0.59 | 2.79E-06 | -0.01 | 9.45E-01 | 0.00 | 1.00E+00 |
| FBgn0039019 | HP1c | 924 | 0.74 | 1.49E-08 | 0.02 | 8.82E-01 | -0.01 | 1.00E+00 |
| FBgn0035829 | HP4 | 943 | 0.86 | 1.33E-08 | -0.01 | 9.47E-01 | 0.00 | 1.00E+00 |
| FBgn0036992 | Hpd | 47 | -0.06 | 8.29E-01 | 1.06 | 2.78E-02 | 0.00 | 1.00E+00 |
| FBgn0062928 | hpRNA:CR33940 | 1575 | 0.04 | 8.63E-01 | 0.08 | 3.50E-01 | -0.63 | 6.35E-05 |
| FBgn0261239 | Hr39 | 2570 | -0.62 | 2.60E-04 | 0.03 | 7.29E-01 | 0.00 | 1.00E+00 |
| FBgn0004838 | Hrb27C | 27370 | -0.74 | 5.65E-06 | 0.00 | 9.93E-01 | 0.00 | 1.00E+00 |
| FBgn0015949 | hrg | 4460 | -0.65 | 6.58E-08 | -0.01 | 9.30E-01 | 0.00 | 1.00E+00 |
| FBgn0001217 | Hsc70-2 | 69 | 6.71 | 3.42E-10 | -0.17 | 2.55E-01 | -0.01 | 1.00E+00 |
| FBgn0266599 | Hsc70-4 | 117280 | 0.62 | 2.13E-13 | -0.02 | 8.37E-01 | -0.01 | 1.00E+00 |
| FBgn0001225 | Hsp26 | 9822 | 1.22 | 1.08E-09 | -0.76 | 2.25E-05 | -0.01 | 9.36E-01 |
| FBgn0015245 | Hsp60A | 10641 | 0.72 | 1.47E-15 | 0.03 | 7.32E-01 | 0.00 | 1.00E+00 |
| FBgn0001229 | Hsp67Bc | 154 | 0.75 | 5.60E-03 | -0.25 | 1.02E-01 | 1.19 | 2.82E-04 |
| FBgn0001230 | Hsp68 | 4283 | 2.04 | 3.72E-05 | -0.23 | 1.53E-01 | 0.00 | 1.00E+00 |
| FBgn0013276 | Hsp70Ab | 244 | 1.56 | 5.11E-05 | -0.04 | 6.54E-01 | 0.00 | 1.00E+00 |
| FBgn0013278 | Hsp70Bb | 477 | 3.65 | 6.59E-06 | -0.04 | 6.85E-01 | 0.00 | 1.00E+00 |
| FBgn0013279 | Hsp70Bc | 336 | 0.14 | 5.78E-01 | -0.60 | 2.76E-02 | 0.00 | 1.00E+00 |
| FBgn0061198 | HSPC300 | 346 | 0.67 | 1.15E-03 | 0.03 | 7.98E-01 | 0.00 | 1.00E+00 |
| FBgn0001235 | hth | 8836 | -1.34 | 8.95E-12 | -0.03 | 7.37E-01 | 0.00 | 1.00E+00 |
| FBgn0010389 | htl | 655 | -0.47 | 5.38E-04 | -0.68 | 7.42E-11 | -0.03 | 5.57E-01 |
| FBgn0033968 | hui | 830 | 1.30 | 1.52E-31 | -0.25 | 1.07E-03 | -0.40 | 2.25E-06 |
| FBgn0036556 | hzg | 1250 | -0.84 | 3.30E-07 | -0.04 | 6.88E-01 | 0.00 | 1.00E+00 |
| FBgn0286204 | ich | 15 | -0.15 | 4.49E-01 | 0.76 | 3.76E-02 | 0.82 | 3.10E-02 |
| FBgn0001248 | Idh | 11385 | 0.64 | 1.66E-18 | -0.41 | 1.55E-11 | -0.79 | 3.46E-41 |
| FBgn0036690 | Ilp8 | 61 | 0.10 | 4.72E-01 | 7.14 | 7.50E-17 | 0.00 | 1.00E+00 |
| FBgn0285926 | Imp | 2083 | -0.92 | 7.76E-06 | 0.01 | 9.32E-01 | 0.00 | 1.00E+00 |
| FBgn0001255 | ImpE3 | 1673 | 0.60 | 8.75E-03 | -0.03 | 8.00E-01 | -0.01 | 1.00E+00 |
| FBgn0001256 | ImpL1 | 53 | 0.18 | 4.69E-01 | -2.17 | 5.55E-05 | -0.01 | 1.00E+00 |
| FBgn0001263 | inaD | 152 | -1.09 | 6.41E-06 | -0.33 | 1.27E-01 | -0.05 | 5.16E-01 |
| FBgn0039459 | IntS12 | 266 | 0.59 | 4.56E-03 | 0.12 | 1.91E-01 | 0.84 | 3.22E-06 |
| FBgn0262117 | IntS3 | 2249 | -0.72 | 5.69E-13 | -0.03 | 8.06E-01 | 0.00 | 1.00E+00 |
| FBgn0027106 | Inx7 | 34 | 1.67 | 7.61E-05 | 0.00 | 9.88E-01 | -0.01 | 1.00E+00 |
| FBgn0259683 | Ir40a | 56 | 0.12 | 4.36E-01 | 5.29 | 3.59E-25 | 0.01 | 1.00E+00 |
| FBgn0050081 | Ir51b | 36 | -0.63 | 1.99E-02 | 0.00 | 9.89E-01 | 0.00 | 1.00E+00 |
| FBgn0011774 | Irbp | 458 | -0.04 | 8.60E-01 | 1.22 | 2.35E-45 | 0.02 | 6.08E-01 |
| FBgn0036126 | Irbp18 | 342 | -0.22 | 2.45E-01 | 1.38 | 2.54E-33 | 0.00 | 1.00E+00 |
| FBgn0037637 | IscU | 1069 | -0.24 | 1.13E-01 | 1.08 | 5.20E-30 | 0.00 | 1.00E+00 |
| FBgn0034005 | ItgaPS4 | 26 | 0.83 | 6.27E-03 | 0.28 | 1.34E-01 | 0.00 | 1.00E+00 |
| FBgn0010053 | Jheh1 | 254 | 0.43 | 2.53E-02 | -0.67 | 4.97E-05 | 0.00 | 1.00E+00 |
| FBgn0028424 | JhI-26 | 62 | -0.06 | 8.21E-01 | 1.56 | 1.78E-03 | 0.00 | 1.00E+00 |
| FBgn0020906 | Jon25Bi | 21 | -0.13 | 2.46E-01 | -2.69 | 6.00E-03 | -0.01 | 1.00E+00 |
| FBgn0031654 | Jon25Bii | 14 | -0.12 | 2.89E-01 | -1.07 | 2.92E-02 | -0.01 | 1.00E+00 |
| FBgn0035665 | Jon65Aiii | 69 | -0.02 | 8.87E-01 | -3.27 | 4.09E-03 | -0.02 | 9.06E-01 |
| FBgn0250815 | Jon65Aiv | 95 | -0.01 | 9.39E-01 | -5.49 | 2.64E-04 | -0.02 | 1.00E+00 |
| FBgn0025820 | JTBR | 391 | 0.66 | 1.51E-04 | 0.04 | 6.21E-01 | 0.00 | 1.00E+00 |
| FBgn0001296 | kar | 177 | 0.71 | 1.82E-03 | 0.08 | 3.88E-01 | -0.01 | 1.00E+00 |
| FBgn0015399 | kek1 | 2777 | -0.75 | 1.30E-07 | -0.10 | 1.92E-01 | 0.15 | 8.10E-02 |
| FBgn0039925 | Kif3C | 10 | -0.20 | 3.61E-01 | 0.22 | 1.84E-01 | 1.55 | 9.41E-03 |
| FBgn0001323 | knrl | 445 | -0.98 | 3.52E-07 | 0.18 | 9.48E-02 | 0.00 | 1.00E+00 |
| FBgn0034098 | krimp | 58 | 0.81 | 5.98E-03 | 0.02 | 8.22E-01 | 0.53 | 3.30E-02 |
| FBgn0040206 | krz | 1877 | -0.74 | 4.98E-07 | 0.02 | 8.67E-01 | 0.00 | 1.00E+00 |
| FBgn0040890 | ksh | 432 | 0.80 | 1.06E-04 | 0.01 | 9.07E-01 | 0.00 | 1.00E+00 |
| FBgn0041627 | Ku80 | 500 | -0.25 | 7.23E-02 | 1.10 | 1.85E-41 | 0.02 | 6.49E-01 |
| FBgn0036667 | kud | 749 | 0.74 | 1.15E-04 | 0.01 | 9.30E-01 | 0.00 | 1.00E+00 |
| FBgn0259984 | kuz | 2737 | -0.59 | 1.26E-03 | -0.04 | 6.48E-01 | 0.00 | 1.00E+00 |
| FBgn0040153 | l(1)G0469 | 354 | -0.48 | 2.80E-02 | -0.66 | 2.67E-03 | -0.01 | 1.00E+00 |
| FBgn0284251 | l(2)05287 | 1676 | -0.58 | 6.07E-06 | 0.03 | 8.01E-01 | 0.00 | 1.00E+00 |
| FBgn0002031 | l(2)37Cc | 3950 | 0.63 | 7.33E-12 | 0.12 | 4.75E-02 | 0.00 | 1.00E+00 |
| FBgn0002121 | l(2)gl | 5956 | -0.66 | 9.23E-06 | -0.03 | 7.39E-01 | 0.00 | 1.00E+00 |
| FBgn0284244 | l(2)k05911 | 1160 | -0.03 | 8.80E-01 | 0.95 | 2.58E-13 | -0.05 | 3.65E-01 |
| FBgn0035617 | l(3)psg2 | 1918 | -0.65 | 2.42E-05 | 0.04 | 6.68E-01 | 0.00 | 1.00E+00 |

| | | | | | | | | |
|---|---|---|---|---|---|---|---|---|
| FBgn0008651 | lbl | 44 | 0.20 | 2.70E-01 | -2.03 | 4.28E-05 | 0.00 | 1.00E+00 |
| FBgn0001258 | Ldh | 467 | 0.64 | 4.76E-03 | 1.77 | 2.03E-15 | 0.07 | 3.52E-01 |
| FBgn0016675 | Lectin-galC1 | 18 | 0.05 | 7.07E-01 | 1.81 | 1.00E-02 | 0.97 | 3.10E-02 |
| FBgn0034877 | levy | 2150 | 0.87 | 1.58E-07 | 0.02 | 8.52E-01 | 0.00 | 1.00E+00 |
| FBgn0025692 | Lfg | 29 | -0.11 | 6.55E-01 | -0.99 | 2.30E-02 | -0.02 | 8.66E-01 |
| FBgn0039907 | lgs | 3040 | -0.63 | 1.33E-06 | -0.03 | 7.79E-01 | 0.00 | 1.00E+00 |
| FBgn0041111 | lilli | 1423 | -0.65 | 1.41E-04 | -0.01 | 9.51E-01 | 0.00 | 1.00E+00 |
| FBgn0026411 | Lim1 | 66 | 0.31 | 1.24E-01 | -2.43 | 3.62E-08 | 0.01 | 1.00E+00 |
| FBgn0032253 | LManI | 47 | -0.30 | 2.32E-01 | -0.82 | 3.71E-03 | -0.16 | 1.60E-01 |
| FBgn0027611 | LManII | 658 | 0.16 | 5.25E-01 | -0.98 | 2.09E-04 | -0.03 | 7.07E-01 |
| FBgn0051044 | lncRNA:CR31044 | 109 | -0.62 | 1.51E-02 | -0.06 | 4.99E-01 | 0.00 | 1.00E+00 |
| FBgn0063127 | lncRNA:CR33938 | 820 | 2.12 | 4.16E-22 | 0.14 | 1.56E-01 | 0.00 | 1.00E+00 |
| FBgn0267910 | lncRNA:CR34335 | 7316 | 1.25 | 1.68E-05 | 0.02 | 8.76E-01 | 0.00 | 1.00E+00 |
| FBgn0259993 | lncRNA:CR42491 | 1363 | 0.61 | 4.70E-06 | 0.06 | 4.52E-01 | -0.94 | 5.20E-17 |
| FBgn0261813 | lncRNA:CR42755 | 10 | -0.25 | 2.06E-01 | -0.12 | 3.37E-01 | -2.47 | 2.02E-03 |
| FBgn0262620 | lncRNA:CR43144 | 765 | 0.08 | 7.87E-01 | 5.09 | 1.28E-279 | 0.02 | 7.45E-01 |
| FBgn0262886 | lncRNA:CR43241 | 230 | -1.36 | 3.84E-09 | 0.04 | 7.17E-01 | 0.00 | 1.00E+00 |
| FBgn0286827 | lncRNA:CR43334 | 202 | 0.33 | 1.75E-01 | 0.68 | 2.98E-02 | -0.03 | 7.07E-01 |
| FBgn0263415 | lncRNA:CR43461 | 36 | -0.06 | 8.34E-01 | 0.61 | 3.42E-02 | 0.00 | 1.00E+00 |
| FBgn0263617 | lncRNA:CR43626 | 25 | -0.89 | 4.25E-03 | 0.00 | 9.88E-01 | 0.00 | 1.00E+00 |
| FBgn0263626 | lncRNA:CR43635 | 12 | 0.86 | 5.82E-03 | 0.02 | 8.52E-01 | 0.00 | 1.00E+00 |
| FBgn0264721 | lncRNA:CR43989 | 32 | -0.04 | 9.11E-01 | 0.85 | 7.12E-03 | 0.01 | 1.00E+00 |
| FBgn0264794 | lncRNA:CR44024 | 20 | 0.03 | 9.25E-01 | 2.10 | 2.05E-06 | 0.00 | 1.00E+00 |
| FBgn0264834 | lncRNA:CR44042 | 177 | -0.77 | 1.55E-03 | 0.63 | 1.55E-03 | 0.00 | 1.00E+00 |
| FBgn0264942 | lncRNA:CR44111 | 42 | 0.09 | 5.34E-01 | 5.99 | 1.17E-27 | 0.00 | 1.00E+00 |
| FBgn0264984 | lncRNA:CR44135 | 23 | -0.29 | 1.06E-01 | 1.21 | 3.43E-03 | -0.01 | 1.00E+00 |
| FBgn0265422 | lncRNA:CR44334 | 427 | -0.68 | 3.70E-04 | 0.20 | 6.50E-02 | 0.01 | 8.66E-01 |
| FBgn0265497 | lncRNA:CR44366 | 18 | -0.66 | 1.58E-02 | -0.03 | 7.38E-01 | 0.00 | 1.00E+00 |
| FBgn0265651 | lncRNA:CR44458 | 17 | -0.04 | 8.81E-01 | 1.51 | 3.11E-03 | -0.01 | 1.00E+00 |
| FBgn0266049 | lncRNA:CR44814 | 120 | -1.88 | 1.42E-09 | 0.04 | 7.13E-01 | 0.00 | 1.00E+00 |
| FBgn0266051 | lncRNA:CR44816 | 80 | -1.42 | 2.27E-06 | 0.00 | 9.92E-01 | -0.01 | 1.00E+00 |
| FBgn0266414 | lncRNA:CR45054 | 19 | 0.14 | 3.83E-01 | 0.79 | 3.16E-02 | 0.00 | 1.00E+00 |
| FBgn0266549 | lncRNA:CR45102 | 308 | -2.24 | 1.22E-12 | 0.02 | 8.91E-01 | 0.00 | 1.00E+00 |
| FBgn0266691 | lncRNA:CR45181 | 18 | 0.87 | 5.43E-03 | 0.52 | 4.50E-02 | 0.01 | 9.89E-01 |
| FBgn0267793 | lncRNA:CR45232 | 366 | -0.73 | 3.43E-04 | -0.15 | 1.61E-01 | 0.00 | 1.00E+00 |
| FBgn0266866 | lncRNA:CR45327 | 19 | -0.60 | 2.28E-02 | -0.03 | 8.04E-01 | -0.01 | 1.00E+00 |
| FBgn0266958 | lncRNA:CR45409 | 159 | -1.00 | 1.92E-05 | -0.03 | 7.29E-01 | 0.00 | 1.00E+00 |
| FBgn0267029 | lncRNA:CR45473 | 135 | -0.38 | 7.22E-02 | 0.78 | 1.73E-05 | 0.01 | 1.00E+00 |
| FBgn0267073 | lncRNA:CR45517 | 44 | -0.04 | 8.97E-01 | 2.24 | 6.58E-11 | 0.04 | 5.85E-01 |
| FBgn0267126 | lncRNA:CR45566 | 684 | 0.92 | 7.68E-13 | 0.05 | 6.05E-01 | 0.00 | 1.00E+00 |
| FBgn0267635 | lncRNA:CR45973 | 81 | -0.26 | 3.03E-01 | 1.44 | 1.55E-08 | 0.00 | 1.00E+00 |
| FBgn0278598 | lncRNA:CR46268 | 190 | -1.19 | 1.56E-06 | 0.03 | 7.60E-01 | 0.00 | 1.00E+00 |
| FBgn0267585 | lncRNA:dntRL | 95 | -0.03 | 9.12E-01 | 1.02 | 6.17E-06 | 0.00 | 1.00E+00 |
| FBgn0022238 | lolal | 3178 | -1.02 | 7.82E-19 | 0.05 | 5.82E-01 | 0.00 | 1.00E+00 |
| FBgn0052699 | LPCAT | 956 | -0.61 | 5.17E-07 | 0.00 | 9.93E-01 | 0.00 | 1.00E+00 |
| FBgn0028582 | lqf | 1867 | -0.59 | 1.74E-08 | -0.01 | 9.20E-01 | 0.00 | 1.00E+00 |
| FBgn0267861 | Maf1 | 2161 | -0.95 | 1.80E-17 | 0.00 | 9.83E-01 | 0.00 | 1.00E+00 |
| FBgn0029979 | mahe | 4391 | -0.78 | 1.42E-04 | -0.03 | 7.25E-01 | 0.00 | 1.00E+00 |
| FBgn0032382 | Mal-B2 | 163 | 0.24 | 3.10E-01 | 2.29 | 9.15E-07 | 0.00 | 1.00E+00 |
| FBgn0034282 | Mapmodulin | 8990 | -0.84 | 1.93E-15 | 0.05 | 5.94E-01 | 0.00 | 1.00E+00 |
| FBgn0039972 | Marf1 | 212 | 0.44 | 4.74E-02 | 1.26 | 1.83E-10 | -0.01 | 1.00E+00 |
| FBgn0039914 | mav | 2998 | -0.75 | 1.06E-16 | 0.07 | 3.67E-01 | 0.00 | 1.00E+00 |
| FBgn0035811 | Mcad | 4056 | 0.66 | 1.26E-09 | -0.26 | 2.17E-03 | 0.00 | 1.00E+00 |
| FBgn0032929 | Mcm10 | 613 | -0.83 | 1.75E-08 | 0.06 | 4.99E-01 | 0.02 | 7.84E-01 |
| FBgn0284442 | Mcm3 | 2442 | 0.59 | 1.94E-05 | -0.01 | 9.16E-01 | 0.00 | 1.00E+00 |
| FBgn0032116 | Mco1 | 31 | -0.16 | 5.19E-01 | -0.74 | 2.10E-02 | 0.00 | 1.00E+00 |
| FBgn0262782 | Mdh1 | 2243 | 0.69 | 7.72E-06 | -0.03 | 7.73E-01 | 0.00 | 1.00E+00 |
| FBgn0262559 | Mdh2 | 2464 | 0.61 | 1.94E-06 | 0.05 | 5.86E-01 | 0.00 | 1.00E+00 |
| FBgn0034707 | MED16 | 746 | 0.29 | 2.14E-02 | -0.58 | 1.75E-11 | 0.00 | 1.00E+00 |
| FBgn0039923 | MED26 | 3695 | -0.93 | 2.24E-08 | -0.04 | 6.74E-01 | 0.00 | 1.00E+00 |
| FBgn0039337 | MED28 | 361 | 0.64 | 1.35E-03 | 0.03 | 7.60E-01 | 0.00 | 1.00E+00 |
| FBgn0024556 | mEFTu1 | 2555 | 0.70 | 8.55E-10 | 0.15 | 4.44E-02 | 0.00 | 1.00E+00 |
| FBgn0039851 | mey | 1124 | 1.29 | 3.47E-17 | 1.26 | 1.81E-23 | 1.01 | 3.55E-15 |
| FBgn0031307 | MFS3 | 3325 | 0.69 | 9.89E-07 | 0.00 | 9.93E-01 | 0.00 | 1.00E+00 |
| FBgn0025814 | Mgstl | 739 | 0.87 | 2.43E-05 | -0.03 | 7.47E-01 | 0.00 | 1.00E+00 |
| FBgn0036333 | MICAL-like | 1637 | -0.63 | 2.14E-13 | -0.12 | 3.63E-02 | 0.00 | 1.00E+00 |
| FBgn0261963 | mid | 203 | 0.43 | 4.48E-02 | -0.58 | 3.07E-03 | 0.01 | 1.00E+00 |
| FBgn0039250 | Mink | 2302 | -0.60 | 9.89E-07 | -0.06 | 4.88E-01 | 0.00 | 1.00E+00 |
| FBgn0026061 | Mipp1 | 38 | 1.35 | 5.54E-04 | 0.00 | 9.98E-01 | 0.00 | 1.00E+00 |
| FBgn0263112 | Mitf | 2077 | -0.73 | 4.40E-07 | 0.00 | 9.90E-01 | 0.04 | 4.11E-01 |
| FBgn0024326 | Mkk4 | 1661 | -0.65 | 1.15E-11 | -0.02 | 9.00E-01 | 0.00 | 1.00E+00 |
| FBgn0002774 | mle | 2065 | -0.68 | 3.04E-14 | -0.08 | 2.61E-01 | 0.00 | 1.00E+00 |
| FBgn0259168 | mnb | 2631 | -0.67 | 4.00E-06 | -0.05 | 6.05E-01 | 0.00 | 1.00E+00 |
| FBgn0263241 | Mocs1 | 299 | 0.02 | 9.55E-01 | 1.00 | 6.99E-12 | -0.02 | 7.07E-01 |
| FBgn0026409 | Mpcp2 | 7899 | 0.69 | 8.14E-23 | 0.05 | 5.60E-01 | 0.00 | 1.00E+00 |
| FBgn0020270 | mre11 | 1699 | 0.34 | 1.30E-03 | 1.11 | 7.59E-75 | 0.35 | 3.50E-07 |
| FBgn0034091 | mrj | 966 | -0.79 | 8.03E-13 | -0.01 | 9.45E-01 | 0.00 | 1.00E+00 |
| FBgn0032456 | MRP | 3772 | 0.06 | 7.35E-01 | 1.55 | 5.28E-48 | 0.00 | 1.00E+00 |
| FBgn0011787 | mRpL12 | 969 | 0.91 | 4.58E-07 | 0.02 | 8.61E-01 | 0.00 | 1.00E+00 |

| | | | | | | | | |
|---|---|---|---|---|---|---|---|---|
| FBgn0035122 | mRpL17 | 461 | 0.68 | 2.30E-05 | 0.04 | 6.74E-01 | 0.00 | 1.00E+00 |
| FBgn0036135 | mRpL2 | 767 | 0.65 | 2.40E-05 | 0.01 | 9.37E-01 | 0.00 | 1.00E+00 |
| FBgn0040907 | mRpL33 | 284 | 1.11 | 1.42E-07 | 0.00 | 9.86E-01 | -0.01 | 1.00E+00 |
| FBgn0038923 | mRpL35 | 302 | 0.61 | 9.42E-04 | -0.01 | 9.37E-01 | 0.00 | 1.00E+00 |
| FBgn0030552 | mRpL38 | 817 | 0.65 | 1.45E-05 | 0.02 | 8.73E-01 | 0.00 | 1.00E+00 |
| FBgn0037330 | mRpL44 | 725 | 0.66 | 2.81E-05 | 0.02 | 8.52E-01 | 0.00 | 1.00E+00 |
| FBgn0031357 | mRpL48 | 312 | 0.58 | 5.52E-03 | 0.01 | 9.18E-01 | 0.00 | 1.00E+00 |
| FBgn0030433 | mRpL49 | 450 | 0.84 | 2.67E-05 | 0.03 | 8.00E-01 | 0.00 | 1.00E+00 |
| FBgn0028648 | mRpL50 | 407 | 0.69 | 2.41E-04 | 0.01 | 9.24E-01 | 0.00 | 1.00E+00 |
| FBgn0032053 | mRpL51 | 425 | 0.96 | 1.82E-08 | 0.01 | 9.68E-01 | -0.01 | 1.00E+00 |
| FBgn0033208 | mRpL52 | 367 | 0.95 | 2.88E-07 | 0.03 | 7.47E-01 | 0.00 | 1.00E+00 |
| FBgn0038678 | mRpL55 | 253 | 0.72 | 4.36E-04 | 0.00 | 9.72E-01 | 0.00 | 1.00E+00 |
| FBgn0044030 | mRpS14 | 193 | 1.02 | 3.28E-05 | 0.07 | 4.31E-01 | 0.00 | 1.00E+00 |
| FBgn0030572 | mRpS25 | 392 | 0.59 | 8.81E-04 | 0.03 | 7.83E-01 | 0.00 | 1.00E+00 |
| FBgn0044510 | mRpS5 | 493 | -0.93 | 9.94E-07 | -0.01 | 9.26E-01 | 0.00 | 1.00E+00 |
| FBgn0011666 | msi | 2622 | -0.62 | 3.63E-04 | 0.01 | 9.71E-01 | 0.00 | 1.00E+00 |
| FBgn0027948 | msps | 6613 | -0.62 | 3.50E-06 | -0.01 | 9.70E-01 | 0.00 | 1.00E+00 |
| FBgn0013672 | mt:ATPase6 | 7880 | -3.31 | 5.37E-41 | -0.02 | 8.94E-01 | 0.00 | 1.00E+00 |
| FBgn0013673 | mt:ATPase8 | 1600 | -3.84 | 3.78E-230 | 0.02 | 9.06E-01 | 0.02 | 9.34E-01 |
| FBgn0013674 | mt:CoI | 59337 | -0.61 | 1.51E-02 | 0.01 | 8.92E-01 | 0.00 | 1.00E+00 |
| FBgn0013675 | mt:CoII | 18791 | -1.47 | 1.12E-09 | 0.00 | 9.75E-01 | -0.01 | 1.00E+00 |
| FBgn0013686 | mt:lrRNA | 420131 | -2.84 | 3.45E-10 | 0.02 | 8.37E-01 | 0.00 | 1.00E+00 |
| FBgn0013679 | mt:ND1 | 4891 | -3.62 | 7.13E-14 | 0.02 | 8.91E-01 | 0.00 | 1.00E+00 |
| FBgn0013680 | mt:ND2 | 1185 | -4.38 | 3.24E-10 | -0.03 | 7.42E-01 | 0.00 | 1.00E+00 |
| FBgn0013681 | mt:ND3 | 583 | -3.85 | 7.22E-11 | 0.02 | 8.24E-01 | 0.01 | 1.00E+00 |
| FBgn0262952 | mt:ND4 | 3538 | -2.38 | 7.67E-11 | 0.04 | 7.12E-01 | 0.00 | 1.00E+00 |
| FBgn0013684 | mt:ND5 | 6582 | -2.51 | 1.53E-58 | 0.00 | 9.88E-01 | 0.03 | 4.80E-01 |
| FBgn0013685 | mt:ND6 | 440 | -0.91 | 5.02E-03 | -0.02 | 8.68E-01 | 0.00 | 1.00E+00 |
| FBgn0013688 | mt:srRNA | 534 | -0.37 | 6.27E-02 | 1.14 | 2.01E-02 | 0.00 | 1.00E+00 |
| FBgn0013698 | mt:tRNA:Leu-TAG | 25 | -2.56 | 3.95E-05 | 0.02 | 9.10E-01 | 0.00 | 1.00E+00 |
| FBgn0013710 | mt:tRNA:Tyr-GTA | 24 | -0.61 | 2.14E-02 | -0.01 | 9.71E-01 | 0.00 | 1.00E+00 |
| FBgn0028956 | mthl3 | 3240 | -0.66 | 8.47E-16 | 0.05 | 5.44E-01 | 0.00 | 1.00E+00 |
| FBgn0034219 | mthl4 | 587 | -0.91 | 1.26E-09 | -0.02 | 8.83E-01 | 0.00 | 1.00E+00 |
| FBgn0052475 | mthl8 | 493 | -0.52 | 7.63E-04 | -0.06 | 6.13E-01 | 10.83 | 1.59E-36 |
| FBgn0025352 | Mtpbeta | 4531 | 0.62 | 3.72E-12 | -0.09 | 1.74E-01 | 0.00 | 1.00E+00 |
| FBgn0262737 | mub | 5736 | -0.69 | 5.78E-07 | -0.05 | 5.79E-01 | 0.00 | 1.00E+00 |
| FBgn0002887 | mus201 | 555 | -0.84 | 2.67E-06 | 0.57 | 8.19E-05 | 0.00 | 1.00E+00 |
| FBgn0264272 | mwh | 95 | -0.49 | 2.83E-02 | -1.28 | 1.73E-08 | 0.00 | 1.00E+00 |
| FBgn0262656 | Myc | 3026 | -1.25 | 4.29E-08 | 0.01 | 9.30E-01 | 0.00 | 1.00E+00 |
| FBgn0026199 | myo | 1852 | -1.04 | 1.52E-11 | 0.02 | 8.64E-01 | 0.00 | 1.00E+00 |
| FBgn0039157 | Myo95E | 879 | -0.68 | 4.80E-07 | 0.00 | 9.96E-01 | 0.00 | 1.00E+00 |
| FBgn0051216 | Naam | 156 | -0.21 | 3.64E-01 | 0.62 | 4.56E-04 | 0.01 | 1.00E+00 |
| FBgn0086904 | Nacalpha | 17191 | 0.66 | 9.52E-13 | -0.05 | 5.39E-01 | 0.00 | 1.00E+00 |
| FBgn0260795 | NaPi-III | 1252 | -0.59 | 1.45E-05 | -0.07 | 3.45E-01 | -0.04 | 4.03E-01 |
| FBgn0085417 | natalisin | 246 | 1.08 | 1.55E-09 | 0.12 | 1.55E-01 | 0.02 | 6.27E-01 |
| FBgn0013303 | Nca | 871 | -0.82 | 8.10E-09 | -0.03 | 8.10E-01 | 0.00 | 1.00E+00 |
| FBgn0086707 | ncm | 2162 | -1.09 | 8.23E-10 | 0.03 | 7.48E-01 | 0.00 | 1.00E+00 |
| FBgn0031021 | ND-18 | 707 | 0.84 | 1.01E-05 | 0.08 | 3.70E-01 | 0.00 | 1.00E+00 |
| FBgn0266582 | ND-30 | 1362 | 0.67 | 1.79E-09 | 0.04 | 6.37E-01 | 0.00 | 1.00E+00 |
| FBgn0037001 | ND-39 | 2350 | 0.93 | 2.39E-21 | 0.03 | 7.96E-01 | -0.01 | 1.00E+00 |
| FBgn0058002 | ND-AGGG | 176 | -1.41 | 4.06E-09 | 0.04 | 6.30E-01 | 0.00 | 1.00E+00 |
| FBgn0034645 | ND-B12 | 751 | 0.88 | 5.97E-07 | -0.06 | 4.68E-01 | 0.00 | 1.00E+00 |
| FBgn0025839 | ND-B14.5A | 357 | 0.75 | 3.67E-05 | 0.06 | 5.10E-01 | 0.00 | 1.00E+00 |
| FBgn0034576 | ND-B14.7 | 900 | 0.81 | 1.14E-06 | 0.04 | 6.74E-01 | 0.00 | 1.00E+00 |
| FBgn0033961 | ND-B15 | 514 | 0.94 | 2.40E-06 | 0.03 | 7.82E-01 | 0.00 | 1.00E+00 |
| FBgn0029868 | ND-B16.6 | 1491 | 0.95 | 1.26E-13 | 0.02 | 9.07E-01 | 0.00 | 1.00E+00 |
| FBgn0001989 | ND-B17 | 829 | 0.73 | 7.01E-07 | -0.01 | 9.18E-01 | 0.00 | 1.00E+00 |
| FBgn0031436 | ND-B17.2 | 939 | 1.38 | 2.39E-21 | -0.01 | 9.33E-01 | 0.00 | 1.00E+00 |
| FBgn0030605 | ND-B18 | 697 | 0.72 | 1.67E-08 | 0.07 | 4.19E-01 | 0.00 | 1.00E+00 |
| FBgn0085468 | ND-MWFE | 607 | 0.80 | 3.40E-05 | 0.00 | 9.82E-01 | 0.00 | 1.00E+00 |
| FBgn0021967 | ND-PDSW | 1240 | 1.05 | 1.19E-06 | 0.10 | 2.58E-01 | 0.00 | 1.00E+00 |
| FBgn0017430 | Nelf-E | 566 | 0.19 | 2.49E-01 | 0.51 | 4.14E-07 | 0.90 | 5.05E-21 |
| FBgn0027570 | Nep2 | 1864 | 0.24 | 1.37E-01 | -1.19 | 2.14E-22 | -0.72 | 4.22E-08 |
| FBgn0039564 | Nep7 | 118 | 0.99 | 2.44E-04 | 0.02 | 8.45E-01 | 0.10 | 2.75E-01 |
| FBgn0032393 | Nfs1 | 892 | 0.16 | 3.80E-01 | 1.00 | 1.30E-19 | 0.00 | 1.00E+00 |
| FBgn0036101 | NijA | 1536 | -0.22 | 1.07E-01 | -0.38 | 2.26E-04 | -0.67 | 1.96E-10 |
| FBgn0038079 | NijC | 67 | 0.83 | 2.32E-03 | 0.01 | 9.16E-01 | -0.01 | 1.00E+00 |
| FBgn0259896 | NimC1 | 57 | 0.18 | 4.82E-01 | -2.27 | 4.02E-06 | -0.01 | 1.00E+00 |
| FBgn0026401 | Nipped-B | 5024 | -1.01 | 7.71E-10 | 0.01 | 9.69E-01 | 0.00 | 1.00E+00 |
| FBgn0024321 | NK7.1 | 331 | -0.67 | 2.02E-04 | -0.02 | 8.22E-01 | 0.03 | 5.71E-01 |
| FBgn0051547 | NKCC | 84 | 0.03 | 9.36E-01 | 1.28 | 1.77E-03 | -0.01 | 1.00E+00 |
| FBgn0031866 | Nlg2 | 15 | 2.40 | 3.35E-04 | -0.03 | 7.75E-01 | 0.00 | 1.00E+00 |
| FBgn0011817 | nmo | 3519 | -0.72 | 9.06E-07 | 0.04 | 6.54E-01 | 0.00 | 1.00E+00 |
| FBgn0037617 | nom | 365 | -0.63 | 4.56E-04 | 0.05 | 6.11E-01 | 0.01 | 1.00E+00 |
| FBgn0015520 | nonA-l | 647 | -0.65 | 1.76E-06 | -0.11 | 1.51E-01 | 0.00 | 1.00E+00 |
| FBgn0033029 | Not3 | 5518 | -0.69 | 2.88E-13 | -0.05 | 5.72E-01 | 0.00 | 1.00E+00 |
| FBgn0027785 | NP15.6 | 1071 | 1.36 | 1.53E-13 | 0.02 | 8.85E-01 | 0.00 | 1.00E+00 |
| FBgn0031381 | Npc2a | 201 | 0.59 | 1.30E-02 | -0.01 | 9.10E-01 | 0.00 | 1.00E+00 |
| FBgn0035092 | Nplp1 | 42 | -0.16 | 5.61E-01 | -1.43 | 6.65E-05 | 0.00 | 1.00E+00 |

| | | | | | | | | |
|---|---|---|---|---|---|---|---|---|
| FBgn0040717 | Nplp4 | 686 | 1.01 | 6.07E-04 | -0.62 | 1.63E-02 | 0.96 | 4.01E-03 |
| FBgn0261526 | NT1 | 183 | -0.07 | 8.23E-01 | -0.80 | 1.26E-03 | 0.00 | 1.00E+00 |
| FBgn0028411 | Nxt1 | 883 | 0.59 | 7.17E-04 | 0.13 | 1.56E-01 | 0.00 | 1.00E+00 |
| FBgn0027791 | O-fut2 | 218 | 0.96 | 2.28E-05 | 0.00 | 9.76E-01 | 0.00 | 1.00E+00 |
| FBgn0022774 | Oat | 220 | -0.06 | 8.35E-01 | -0.69 | 2.98E-02 | -0.01 | 1.00E+00 |
| FBgn0038344 | obe | 1568 | -0.17 | 2.99E-01 | 0.87 | 8.91E-14 | 0.01 | 1.00E+00 |
| FBgn0046875 | Obp83g | 76 | 1.39 | 1.93E-05 | -0.07 | 4.76E-01 | 0.00 | 1.00E+00 |
| FBgn0039678 | Obp99a | 345 | 2.49 | 2.05E-15 | -2.20 | 1.17E-27 | 0.00 | 1.00E+00 |
| FBgn0027600 | obst-B | 1761 | 0.83 | 9.71E-08 | -0.04 | 6.25E-01 | 0.00 | 1.00E+00 |
| FBgn0028996 | onecut | 28 | -0.17 | 4.64E-01 | 2.85 | 7.82E-12 | 0.04 | 6.14E-01 |
| FBgn0026393 | Or43b | 242 | -0.70 | 3.46E-05 | 0.04 | 7.13E-01 | 0.00 | 1.00E+00 |
| FBgn0039551 | Or98a | 41 | -0.03 | 8.31E-01 | 8.76 | 7.97E-13 | 0.00 | 1.00E+00 |
| FBgn0040279 | Osi14 | 132 | 0.95 | 6.82E-04 | 1.56 | 3.99E-09 | 0.18 | 1.42E-01 |
| FBgn0032197 | ova | 1116 | -0.77 | 3.32E-07 | 0.00 | 9.86E-01 | 0.00 | 1.00E+00 |
| FBgn0011227 | ox | 799 | 2.14 | 2.56E-51 | 0.02 | 8.38E-01 | 0.00 | 1.00E+00 |
| FBgn0260799 | p120ctn | 3379 | -1.11 | 1.81E-16 | 0.00 | 9.86E-01 | 0.00 | 1.00E+00 |
| FBgn0265297 | pAbp | 44968 | -0.66 | 4.08E-04 | 0.02 | 8.24E-01 | 0.00 | 1.00E+00 |
| FBgn0005648 | Pabp2 | 4791 | -0.80 | 3.13E-09 | 0.15 | 7.73E-02 | 0.00 | 1.00E+00 |
| FBgn0060296 | pain | 1156 | -0.10 | 5.67E-01 | -0.63 | 8.12E-08 | -0.03 | 4.82E-01 |
| FBgn0038100 | Paip2 | 1311 | -0.80 | 1.58E-07 | 0.00 | 1.00E+00 | 0.00 | 1.00E+00 |
| FBgn0085432 | pan | 2131 | -0.59 | 6.42E-03 | 0.00 | 9.77E-01 | 0.00 | 1.00E+00 |
| FBgn0035397 | PAN3 | 3490 | -0.71 | 4.02E-11 | -0.10 | 1.74E-01 | 0.00 | 1.00E+00 |
| FBgn0010247 | Parp | 2656 | -0.94 | 3.12E-08 | 0.38 | 6.00E-03 | 0.00 | 1.00E+00 |
| FBgn0036007 | path | 4128 | 0.03 | 8.74E-01 | 0.03 | 8.10E-01 | 0.80 | 5.05E-13 |
| FBgn0027580 | PCB | 2192 | 0.26 | 2.54E-01 | -0.84 | 2.06E-04 | -0.02 | 7.92E-01 |
| FBgn0024841 | Pcd | 661 | 0.96 | 6.82E-08 | -0.02 | 8.22E-01 | 0.00 | 1.00E+00 |
| FBgn0086768 | Pcmt | 1557 | 0.98 | 5.51E-10 | 0.02 | 8.59E-01 | 0.00 | 1.00E+00 |
| FBgn0005655 | PCNA | 4886 | 0.69 | 2.73E-15 | 0.06 | 3.90E-01 | 0.00 | 1.00E+00 |
| FBgn0036580 | PDCD-5 | 930 | 0.91 | 5.47E-08 | 0.04 | 6.66E-01 | 0.00 | 1.00E+00 |
| FBgn0039635 | Pdhb | 1804 | 0.61 | 4.78E-04 | 0.03 | 7.61E-01 | 0.00 | 1.00E+00 |
| FBgn0016694 | Pdp1 | 373 | -1.14 | 5.67E-06 | 0.00 | 9.98E-01 | 0.00 | 1.00E+00 |
| FBgn0031969 | pes | 393 | -0.80 | 8.68E-06 | 0.00 | 9.86E-01 | 0.00 | 1.00E+00 |
| FBgn0031530 | Pgant2 | 288 | 0.65 | 9.45E-04 | 0.04 | 6.43E-01 | 0.00 | 1.00E+00 |
| FBgn0031681 | Pgant5 | 2854 | -0.59 | 8.73E-06 | -0.05 | 5.36E-01 | 0.00 | 1.00E+00 |
| FBgn0014869 | Pglym78 | 1405 | 0.87 | 6.68E-09 | 0.25 | 1.76E-02 | 0.00 | 1.00E+00 |
| FBgn0035976 | PGRP-LC | 112 | 0.10 | 7.03E-01 | -0.74 | 1.85E-04 | -1.41 | 7.16E-13 |
| FBgn0035438 | PHGPx | 2555 | 0.73 | 1.23E-05 | 0.04 | 7.06E-01 | 0.00 | 1.00E+00 |
| FBgn0035089 | Phk-3 | 2712 | 0.81 | 2.59E-09 | -0.04 | 6.45E-01 | -0.01 | 1.00E+00 |
| FBgn0002521 | pho | 2284 | -1.27 | 2.01E-21 | -0.06 | 5.03E-01 | 0.00 | 1.00E+00 |
| FBgn0036522 | Phs | 2259 | -0.66 | 1.14E-07 | 0.03 | 7.54E-01 | 0.00 | 1.00E+00 |
| FBgn0033479 | PIG-N | 188 | -0.69 | 2.91E-04 | -0.08 | 3.40E-01 | 0.00 | 1.00E+00 |
| FBgn0086448 | PIG-Q | 352 | -0.64 | 5.87E-05 | -0.10 | 2.17E-01 | -0.01 | 1.00E+00 |
| FBgn0038966 | pinta | 68 | -0.07 | 8.23E-01 | 2.59 | 5.85E-18 | 1.24 | 2.00E-04 |
| FBgn0039924 | PIP4K | 2088 | -0.89 | 2.33E-15 | -0.03 | 8.07E-01 | -0.01 | 1.00E+00 |
| FBgn0030400 | Pits | 2206 | -0.74 | 4.02E-07 | 0.01 | 9.52E-01 | 0.00 | 1.00E+00 |
| FBgn0004872 | piwi | 37 | -0.40 | 1.07E-01 | -0.69 | 2.81E-02 | 0.00 | 1.00E+00 |
| FBgn0022382 | Pka-R2 | 572 | -0.62 | 4.35E-03 | 0.01 | 9.63E-01 | 0.00 | 1.00E+00 |
| FBgn0036192 | Pldn | 144 | -0.60 | 3.59E-03 | -0.02 | 8.38E-01 | 0.00 | 1.00E+00 |
| FBgn0259214 | PMCA | 7693 | -0.73 | 5.33E-08 | -0.01 | 9.33E-01 | 0.00 | 1.00E+00 |
| FBgn0037737 | Pnn | 621 | -0.66 | 3.63E-06 | 0.00 | 9.86E-01 | 0.00 | 1.00E+00 |
| FBgn0036696 | Pop5 | 239 | 0.63 | 2.04E-03 | 0.17 | 9.54E-02 | 0.01 | 1.00E+00 |
| FBgn0003130 | Poxn | 231 | 0.63 | 5.12E-04 | 0.64 | 4.29E-06 | 0.05 | 3.88E-01 |
| FBgn0053508 | ppk13 | 1275 | -0.27 | 1.10E-01 | 1.86 | 1.49E-64 | 0.45 | 1.10E-03 |
| FBgn0261363 | PPO3 | 59 | 0.63 | 1.08E-02 | 0.03 | 7.94E-01 | 0.00 | 1.00E+00 |
| FBgn0011474 | PR-Set7 | 2050 | -0.63 | 3.34E-05 | 0.01 | 9.28E-01 | 0.00 | 1.00E+00 |
| FBgn0033635 | Prip | 107 | -0.08 | 7.73E-01 | -0.62 | 4.40E-03 | -0.06 | 3.60E-01 |
| FBgn0086134 | Prosalpha2 | 4106 | 0.70 | 1.72E-16 | -0.03 | 7.26E-01 | -0.03 | 5.00E-01 |
| FBgn0261394 | Prosalpha3 | 3487 | 0.71 | 2.64E-08 | -0.03 | 7.85E-01 | 0.00 | 1.00E+00 |
| FBgn0027587 | Prp4k | 992 | -0.65 | 1.36E-11 | 0.03 | 7.55E-01 | 0.16 | 1.67E-02 |
| FBgn0033518 | Prx2540-2 | 56 | 0.53 | 4.22E-02 | -0.62 | 1.93E-02 | 0.00 | 1.00E+00 |
| FBgn0035770 | pst | 722 | 0.88 | 2.24E-04 | 0.18 | 1.36E-01 | 0.00 | 1.00E+00 |
| FBgn0026379 | Pten | 1333 | -0.83 | 2.36E-09 | -0.05 | 5.30E-01 | 0.00 | 1.00E+00 |
| FBgn0034085 | Ptp52F | 12 | -0.16 | 4.59E-01 | -0.29 | 1.17E-01 | -0.60 | 3.13E-02 |
| FBgn0014007 | Ptp69D | 2086 | -0.89 | 2.12E-13 | -0.14 | 6.28E-02 | 0.00 | 1.00E+00 |
| FBgn0243512 | puc | 688 | -0.86 | 6.08E-06 | 0.00 | 9.80E-01 | -0.01 | 1.00E+00 |
| FBgn0033226 | puml | 453 | 0.27 | 1.11E-01 | 0.65 | 3.52E-09 | 0.01 | 1.00E+00 |
| FBgn0022361 | Pur-alpha | 1438 | -1.00 | 6.23E-08 | 0.01 | 9.30E-01 | 0.00 | 1.00E+00 |
| FBgn0004577 | Pxd | 86 | -0.01 | 9.75E-01 | -0.87 | 3.05E-02 | -0.03 | 6.72E-01 |
| FBgn0033649 | pyr | 313 | -0.85 | 3.77E-07 | -0.05 | 5.43E-01 | 0.00 | 1.00E+00 |
| FBgn0052412 | QC | 164 | 0.44 | 5.93E-02 | -0.01 | 9.12E-01 | -1.14 | 2.11E-06 |
| FBgn0022987 | qkr54B | 1628 | -1.10 | 1.29E-12 | -0.05 | 5.42E-01 | 0.00 | 1.00E+00 |
| FBgn0022986 | qkr58E-1 | 1684 | -0.95 | 7.55E-09 | -0.02 | 8.81E-01 | 0.00 | 1.00E+00 |
| FBgn0022984 | qkr58E-3 | 1887 | -0.97 | 6.58E-09 | 0.06 | 4.61E-01 | 0.00 | 1.00E+00 |
| FBgn0051864 | Qtzl | 82 | -0.35 | 1.62E-01 | 1.34 | 6.16E-08 | -0.01 | 9.19E-01 |
| FBgn0014010 | Rab5 | 3111 | -0.75 | 1.02E-15 | 0.00 | 9.99E-01 | -0.01 | 1.00E+00 |
| FBgn0030200 | RabX2 | 20 | 0.03 | 8.57E-01 | 6.65 | 2.74E-11 | 0.00 | 1.00E+00 |
| FBgn0014011 | Rac2 | 1610 | -0.87 | 1.83E-12 | -0.04 | 6.69E-01 | -0.01 | 1.00E+00 |
| FBgn0020618 | Rack1 | 56381 | 1.07 | 6.74E-36 | -0.32 | 1.48E-05 | 0.00 | 1.00E+00 |
| FBgn0034728 | rad50 | 1403 | -0.96 | 1.79E-10 | 0.87 | 1.40E-13 | 0.01 | 9.73E-01 |

| | | | | | | | | |
|---|---|---|---|---|---|---|---|---|
| FBgn0034646 | Rae1 | 1214 | 0.61 | 3.94E-06 | 0.06 | 4.85E-01 | 0.00 | 1.00E+00 |
| FBgn0003079 | Raf | 1265 | -0.81 | 2.61E-16 | -0.03 | 7.61E-01 | 0.00 | 1.00E+00 |
| FBgn0039110 | RanBP3 | 2099 | 0.62 | 1.98E-08 | -0.02 | 8.88E-01 | -0.01 | 9.85E-01 |
| FBgn0004636 | Rap1 | 4020 | -1.04 | 1.23E-12 | -0.01 | 9.55E-01 | 0.00 | 1.00E+00 |
| FBgn0031745 | rau | 255 | 0.61 | 1.41E-03 | -0.05 | 5.50E-01 | 0.00 | 1.00E+00 |
| FBgn0030479 | Rbp1-like | 2083 | -1.39 | 8.00E-35 | 0.06 | 4.91E-01 | 0.01 | 1.00E+00 |
| FBgn0031047 | Rcd-1 | 3879 | 0.62 | 4.92E-07 | -0.02 | 8.48E-01 | 0.00 | 1.00E+00 |
| FBgn0039644 | rdog | 430 | -0.19 | 4.02E-01 | 2.41 | 3.73E-52 | 0.00 | 1.00E+00 |
| FBgn0264493 | rdx | 1794 | -0.80 | 1.57E-15 | 0.00 | 9.77E-01 | 0.01 | 1.00E+00 |
| FBgn0027375 | RecQ5 | 730 | -1.02 | 3.31E-11 | 0.00 | 1.00E+00 | 0.01 | 1.00E+00 |
| FBgn0021800 | Reph | 522 | -0.83 | 9.85E-06 | 0.07 | 4.30E-01 | 0.01 | 9.70E-01 |
| FBgn0021906 | RFeSP | 1542 | 0.74 | 6.01E-07 | 0.06 | 5.05E-01 | -0.01 | 1.00E+00 |
| FBgn0014020 | Rho1 | 8527 | -0.78 | 9.28E-22 | 0.03 | 7.71E-01 | 0.00 | 1.00E+00 |
| FBgn0265605 | Ric | 315 | -0.61 | 4.46E-04 | -0.02 | 8.59E-01 | 0.00 | 1.00E+00 |
| FBgn0003256 | rl | 1022 | -0.65 | 3.22E-05 | -0.01 | 9.40E-01 | 0.00 | 1.00E+00 |
| FBgn0014022 | Rlb1 | 390 | -0.84 | 1.02E-05 | 0.04 | 6.98E-01 | 0.00 | 1.00E+00 |
| FBgn0262116 | RNASEK | 1235 | -0.70 | 2.47E-07 | 0.09 | 2.58E-01 | -0.03 | 5.42E-01 |
| FBgn0065098 | RNaseMRP:RNA | 80 | 4.59 | 4.42E-10 | 0.57 | 3.60E-02 | -0.03 | 6.87E-01 |
| FBgn0046696 | RNaseP:RNA | 131 | 2.12 | 4.96E-12 | 0.05 | 5.84E-01 | 0.00 | 1.00E+00 |
| FBgn0250838 | roh | 1748 | 0.64 | 8.88E-05 | -0.03 | 7.61E-01 | 0.00 | 1.00E+00 |
| FBgn0005649 | Rox8 | 5628 | -0.89 | 7.39E-27 | -0.13 | 3.05E-02 | 0.00 | 1.00E+00 |
| FBgn0010173 | RpA-70 | 3830 | -0.17 | 1.04E-01 | 0.58 | 8.68E-17 | 0.00 | 1.00E+00 |
| FBgn0262954 | Rpb12 | 562 | 1.01 | 4.13E-07 | 0.06 | 4.95E-01 | -0.01 | 9.40E-01 |
| FBgn0050499 | Rpe | 1477 | 0.82 | 4.26E-11 | -0.02 | 8.91E-01 | 0.00 | 1.00E+00 |
| FBgn0004855 | RpII15 | 1036 | 0.63 | 2.77E-06 | -0.01 | 9.39E-01 | -0.01 | 1.00E+00 |
| FBgn0026373 | RpII33 | 900 | 0.63 | 3.15E-05 | 0.02 | 8.21E-01 | 0.00 | 1.00E+00 |
| FBgn0022981 | rpk | 458 | 0.06 | 7.97E-01 | 0.75 | 1.76E-08 | -0.01 | 1.00E+00 |
| FBgn0036213 | RpL10Ab | 42243 | 0.74 | 3.62E-14 | -0.35 | 3.27E-05 | 0.00 | 1.00E+00 |
| FBgn0013325 | RpL11 | 28172 | 0.92 | 3.13E-18 | -0.02 | 8.26E-01 | 0.00 | 1.00E+00 |
| FBgn0034968 | RpL12 | 29887 | 0.59 | 9.92E-06 | -0.13 | 1.05E-01 | 0.00 | 1.00E+00 |
| FBgn0011272 | RpL13 | 31038 | 1.32 | 7.08E-19 | 0.01 | 9.16E-01 | 0.00 | 1.00E+00 |
| FBgn0037351 | RpL13A | 29649 | 1.58 | 4.20E-41 | -0.03 | 7.96E-01 | 0.00 | 1.00E+00 |
| FBgn0017579 | RpL14 | 24860 | 1.54 | 1.76E-29 | -0.04 | 7.13E-01 | 0.00 | 1.00E+00 |
| FBgn0029897 | RpL17 | 24696 | 1.51 | 2.72E-26 | -0.03 | 7.43E-01 | 0.00 | 1.00E+00 |
| FBgn0035753 | RpL18 | 25719 | 1.26 | 6.13E-19 | -0.04 | 7.01E-01 | 0.00 | 1.00E+00 |
| FBgn0010409 | RpL18A | 33698 | 1.19 | 5.09E-22 | 0.01 | 9.46E-01 | 0.00 | 1.00E+00 |
| FBgn0285950 | RpL19 | 37514 | 0.70 | 2.79E-10 | -0.02 | 8.39E-01 | 0.00 | 1.00E+00 |
| FBgn0034837 | RpL22-like | 14 | 0.31 | 1.65E-01 | -2.96 | 6.20E-04 | 0.00 | 1.00E+00 |
| FBgn0010078 | RpL23 | 29740 | 1.18 | 6.09E-15 | -0.01 | 9.33E-01 | 0.00 | 1.00E+00 |
| FBgn0032518 | RpL24 | 20688 | 1.16 | 1.40E-10 | 0.00 | 9.94E-01 | 0.00 | 1.00E+00 |
| FBgn0036825 | RpL26 | 26262 | 1.18 | 5.38E-19 | -0.09 | 2.57E-01 | 0.00 | 1.00E+00 |
| FBgn0039359 | RpL27 | 30893 | 1.82 | 7.59E-44 | -0.07 | 4.00E-01 | 0.00 | 1.00E+00 |
| FBgn0035422 | RpL28 | 33296 | 0.92 | 3.35E-12 | -0.05 | 5.91E-01 | 0.00 | 1.00E+00 |
| FBgn0016726 | RpL29 | 15027 | 2.47 | 1.56E-58 | -0.03 | 7.67E-01 | 0.00 | 1.00E+00 |
| FBgn0020910 | RpL3 | 84241 | 0.82 | 9.57E-18 | -0.32 | 1.18E-04 | 0.00 | 1.00E+00 |
| FBgn0086710 | RpL30 | 15925 | 0.82 | 5.69E-08 | -0.15 | 9.68E-02 | 0.00 | 1.00E+00 |
| FBgn0285949 | RpL31 | 23945 | 1.87 | 4.08E-31 | -0.07 | 3.96E-01 | 0.00 | 1.00E+00 |
| FBgn0002626 | RpL32 | 26148 | 1.55 | 3.34E-30 | -0.08 | 3.38E-01 | 0.00 | 1.00E+00 |
| FBgn0039406 | RpL34a | 4745 | 0.79 | 2.78E-08 | 0.00 | 9.93E-01 | 0.00 | 1.00E+00 |
| FBgn0037686 | RpL34b | 14511 | 1.00 | 8.70E-10 | 0.00 | 1.00E+00 | 0.00 | 1.00E+00 |
| FBgn0029785 | RpL35 | 22897 | 1.11 | 1.57E-14 | -0.04 | 6.97E-01 | 0.00 | 1.00E+00 |
| FBgn0037328 | RpL35A | 18082 | 1.07 | 1.66E-06 | 0.03 | 7.56E-01 | 0.00 | 1.00E+00 |
| FBgn0002579 | RpL36 | 22750 | 0.63 | 4.03E-05 | -0.05 | 5.93E-01 | 0.00 | 1.00E+00 |
| FBgn0031980 | RpL36A | 20310 | 1.45 | 5.37E-17 | -0.01 | 9.36E-01 | 0.00 | 1.00E+00 |
| FBgn0030616 | RpL37a | 28870 | 1.44 | 2.13E-31 | -0.01 | 9.31E-01 | 0.00 | 1.00E+00 |
| FBgn0261608 | RpL37A | 15974 | 1.17 | 6.51E-15 | -0.05 | 5.40E-01 | 0.00 | 1.00E+00 |
| FBgn0040007 | RpL38 | 10930 | -1.22 | 8.25E-12 | 0.01 | 9.45E-01 | 0.00 | 1.00E+00 |
| FBgn0023170 | RpL39 | 14842 | 1.09 | 6.07E-13 | -0.01 | 9.54E-01 | 0.00 | 1.00E+00 |
| FBgn0003279 | RpL4 | 68623 | 0.74 | 7.97E-10 | -0.01 | 9.31E-01 | 0.00 | 1.00E+00 |
| FBgn0003941 | RpL40 | 28783 | 1.13 | 3.13E-23 | -0.04 | 6.70E-01 | 0.00 | 1.00E+00 |
| FBgn0066084 | RpL41 | 44221 | 1.27 | 1.23E-12 | -0.10 | 2.39E-01 | 0.00 | 1.00E+00 |
| FBgn0039857 | RpL6 | 40858 | 0.66 | 5.68E-09 | -0.02 | 8.96E-01 | 0.00 | 1.00E+00 |
| FBgn0005593 | RpL7 | 46015 | 0.88 | 6.36E-19 | -0.07 | 3.97E-01 | 0.00 | 1.00E+00 |
| FBgn0014026 | RpL7A | 51685 | 0.64 | 1.91E-08 | -0.04 | 6.85E-01 | 0.00 | 1.00E+00 |
| FBgn0261602 | RpL8 | 41973 | 0.93 | 2.53E-13 | -0.03 | 7.65E-01 | 0.00 | 1.00E+00 |
| FBgn0015756 | RpL9 | 33572 | 1.23 | 3.91E-24 | -0.04 | 6.30E-01 | 0.00 | 1.00E+00 |
| FBgn0000100 | RpLP0 | 49121 | 0.88 | 2.69E-10 | -0.02 | 8.64E-01 | 0.00 | 1.00E+00 |
| FBgn0002593 | RpLP1 | 34330 | 1.50 | 2.88E-29 | -0.05 | 6.18E-01 | 0.00 | 1.00E+00 |
| FBgn0003274 | RpLP2 | 29029 | 0.72 | 3.01E-04 | 0.02 | 8.90E-01 | 0.00 | 1.00E+00 |
| FBgn0015283 | Rpn10 | 3237 | 0.60 | 8.41E-07 | -0.04 | 6.60E-01 | -0.01 | 8.51E-01 |
| FBgn0285947 | RpS10b | 22456 | 0.90 | 7.94E-17 | -0.04 | 7.10E-01 | 0.00 | 1.00E+00 |
| FBgn0033699 | RpS11 | 27054 | 1.05 | 7.21E-11 | -0.02 | 8.37E-01 | 0.00 | 1.00E+00 |
| FBgn0286213 | RpS12 | 22585 | 1.49 | 7.12E-26 | -0.27 | 1.12E-02 | 0.00 | 1.00E+00 |
| FBgn0010265 | RpS13 | 36129 | 0.85 | 1.52E-13 | -0.04 | 7.11E-01 | 0.00 | 1.00E+00 |
| FBgn0004403 | RpS14a | 14676 | 0.84 | 1.97E-10 | -0.02 | 8.88E-01 | 0.00 | 1.00E+00 |
| FBgn0004404 | RpS14b | 5724 | 1.39 | 5.22E-15 | 0.00 | 9.94E-01 | 0.00 | 1.00E+00 |
| FBgn0034138 | RpS15 | 32466 | 1.00 | 1.50E-15 | -0.12 | 1.11E-01 | 0.00 | 1.00E+00 |
| FBgn0010198 | RpS15Aa | 22650 | 0.73 | 3.20E-09 | -0.08 | 3.36E-01 | 0.00 | 1.00E+00 |
| FBgn0033555 | RpS15Ab | 2011 | 0.81 | 1.69E-12 | -0.06 | 5.17E-01 | 0.00 | 1.00E+00 |

| | | | | | | | | |
|---|---|---|---|---|---|---|---|---|
| FBgn0005533 | RpS17 | 26903 | 1.72 | 4.39E-37 | -0.06 | 4.59E-01 | 0.00 | 1.00E+00 |
| FBgn0010411 | RpS18 | 32633 | 1.38 | 2.01E-26 | -0.05 | 5.78E-01 | 0.00 | 1.00E+00 |
| FBgn0010412 | RpS19a | 27379 | 1.20 | 1.49E-08 | 0.00 | 9.88E-01 | 0.00 | 1.00E+00 |
| FBgn0004867 | RpS2 | 48438 | 0.78 | 4.36E-11 | -0.12 | 1.17E-01 | 0.00 | 1.00E+00 |
| FBgn0019936 | RpS20 | 22643 | 1.07 | 7.12E-14 | -0.18 | 4.75E-02 | 0.00 | 1.00E+00 |
| FBgn0015521 | RpS21 | 11086 | 0.72 | 1.53E-06 | -0.03 | 8.07E-01 | 0.00 | 1.00E+00 |
| FBgn0033912 | RpS23 | 31632 | 1.84 | 3.62E-35 | -0.04 | 6.97E-01 | 0.00 | 1.00E+00 |
| FBgn0261596 | RpS24 | 32050 | 1.17 | 2.61E-22 | -0.08 | 2.62E-01 | -0.01 | 1.00E+00 |
| FBgn0086472 | RpS25 | 28124 | 0.59 | 1.91E-05 | -0.04 | 6.82E-01 | 0.00 | 1.00E+00 |
| FBgn0261597 | RpS26 | 31285 | 1.52 | 3.40E-35 | -0.02 | 8.47E-01 | 0.00 | 1.00E+00 |
| FBgn0030136 | RpS28b | 20297 | 0.86 | 1.46E-06 | -0.07 | 3.87E-01 | 0.00 | 1.00E+00 |
| FBgn0261599 | RpS29 | 31258 | 0.83 | 1.80E-11 | -0.07 | 4.10E-01 | 0.00 | 1.00E+00 |
| FBgn0002622 | RpS3 | 41096 | 1.00 | 7.80E-28 | -0.10 | 1.22E-01 | 0.00 | 1.00E+00 |
| FBgn0038834 | RpS30 | 22722 | 0.64 | 4.12E-07 | -0.04 | 6.49E-01 | 0.00 | 1.00E+00 |
| FBgn0011284 | RpS4 | 55820 | 1.07 | 4.18E-29 | -0.06 | 4.68E-01 | 0.00 | 1.00E+00 |
| FBgn0002590 | RpS5a | 48391 | 0.92 | 1.61E-23 | -0.07 | 3.37E-01 | 0.00 | 1.00E+00 |
| FBgn0261592 | RpS6 | 46007 | 1.20 | 1.79E-20 | -0.04 | 6.36E-01 | 0.00 | 1.00E+00 |
| FBgn0039757 | RpS7 | 40640 | 0.89 | 5.35E-11 | -0.05 | 5.82E-01 | 0.00 | 1.00E+00 |
| FBgn0039713 | RpS8 | 39353 | 1.28 | 3.54E-25 | -0.04 | 6.64E-01 | 0.00 | 1.00E+00 |
| FBgn0010408 | RpS9 | 31452 | 0.59 | 9.82E-06 | -0.15 | 7.18E-02 | 0.00 | 1.00E+00 |
| FBgn0028686 | Rpt3 | 3750 | 0.69 | 2.73E-15 | -0.02 | 8.50E-01 | -0.01 | 9.19E-01 |
| FBgn0283472 | S6k | 1848 | -1.11 | 1.68E-13 | 0.06 | 4.86E-01 | 0.02 | 7.50E-01 |
| FBgn0262866 | S6kII | 523 | -0.60 | 8.22E-05 | 0.05 | 5.74E-01 | 0.00 | 1.00E+00 |
| FBgn0037672 | sage | 118 | -0.97 | 7.66E-04 | 0.02 | 8.54E-01 | 0.00 | 1.00E+00 |
| FBgn0026371 | SAK | 1348 | -0.58 | 1.68E-07 | -0.05 | 6.10E-01 | 0.01 | 1.00E+00 |
| FBgn0013334 | Sap47 | 1082 | -0.65 | 1.87E-05 | 0.00 | 9.73E-01 | 0.00 | 1.00E+00 |
| FBgn0267378 | sau | 2766 | -0.71 | 2.10E-12 | -0.08 | 2.97E-01 | 0.00 | 1.00E+00 |
| FBgn0051950 | Sbat | 156 | 1.17 | 4.34E-07 | 0.07 | 4.00E-01 | -0.01 | 1.00E+00 |
| FBgn0261872 | scaf6 | 2332 | -0.63 | 8.81E-18 | -0.03 | 7.70E-01 | 0.00 | 1.00E+00 |
| FBgn0038038 | Sccpdh2 | 466 | 1.03 | 4.86E-08 | -0.02 | 8.24E-01 | 0.08 | 2.30E-01 |
| FBgn0020907 | Scp2 | 14 | 0.05 | 7.27E-01 | 0.07 | 5.58E-01 | 0.78 | 4.12E-02 |
| FBgn0021765 | scu | 3185 | 1.00 | 5.91E-14 | -0.11 | 1.65E-01 | 0.00 | 1.00E+00 |
| FBgn0041094 | scyl | 8561 | -1.33 | 8.68E-18 | -0.06 | 4.87E-01 | -0.01 | 1.00E+00 |
| FBgn0003345 | sd | 7654 | -0.68 | 6.92E-06 | -0.01 | 9.09E-01 | 0.00 | 1.00E+00 |
| FBgn0010415 | Sdc | 5389 | -0.69 | 8.04E-06 | -0.01 | 9.30E-01 | 0.00 | 1.00E+00 |
| FBgn0014028 | SdhB | 1621 | 0.63 | 1.89E-06 | 0.00 | 9.76E-01 | 0.00 | 1.00E+00 |
| FBgn0039112 | SdhD | 864 | 0.75 | 4.35E-07 | -0.02 | 8.28E-01 | 0.00 | 1.00E+00 |
| FBgn0053497 | Sdic2 | 15 | -0.38 | 1.03E-01 | 0.06 | 5.87E-01 | 2.96 | 5.80E-04 |
| FBgn0035771 | Sec63 | 3488 | 0.58 | 7.21E-13 | 0.02 | 8.44E-01 | 0.00 | 1.00E+00 |
| FBgn0010414 | SerT | 71 | -0.42 | 8.67E-02 | -1.55 | 5.72E-07 | -0.03 | 6.87E-01 |
| FBgn0040022 | Set1 | 2236 | -1.19 | 1.64E-18 | -0.04 | 6.94E-01 | 0.00 | 1.00E+00 |
| FBgn0003371 | sgg | 3372 | -0.85 | 5.82E-06 | 0.07 | 4.58E-01 | 0.00 | 1.00E+00 |
| FBgn0052423 | shep | 1920 | -0.90 | 1.74E-10 | 0.04 | 6.70E-01 | 0.00 | 1.00E+00 |
| FBgn0003392 | shi | 2787 | -0.60 | 3.04E-06 | -0.03 | 7.56E-01 | 0.00 | 1.00E+00 |
| FBgn0263873 | sick | 3454 | 0.08 | 6.69E-01 | 3.08 | 4.79E-168 | 0.00 | 1.00E+00 |
| FBgn0010762 | simj | 2866 | -0.61 | 1.03E-05 | -0.07 | 4.22E-01 | 0.00 | 1.00E+00 |
| FBgn0037802 | Sirt6 | 173 | -0.63 | 1.58E-03 | 0.03 | 7.51E-01 | 0.00 | 1.00E+00 |
| FBgn0031971 | Sirup | 481 | 0.45 | 4.83E-02 | -0.90 | 1.93E-04 | 0.00 | 1.00E+00 |
| FBgn0031998 | SLC5A11 | 132 | -0.38 | 1.05E-01 | -0.99 | 6.52E-04 | -0.01 | 9.70E-01 |
| FBgn0037810 | sle | 3373 | -0.67 | 2.17E-05 | 0.02 | 8.42E-01 | 0.00 | 1.00E+00 |
| FBgn0037203 | slif | 87 | -0.66 | 1.01E-02 | -0.06 | 5.03E-01 | 0.00 | 1.00E+00 |
| FBgn0040011 | Slmap | 2039 | -0.78 | 5.55E-15 | 0.04 | 7.15E-01 | 0.00 | 1.00E+00 |
| FBgn0040283 | SMC1 | 3037 | -0.64 | 1.85E-06 | -0.02 | 8.19E-01 | 0.00 | 1.00E+00 |
| FBgn0261789 | SmD2 | 1956 | 0.73 | 2.12E-06 | 0.04 | 6.65E-01 | 0.00 | 1.00E+00 |
| FBgn0000426 | SmF | 1111 | 0.73 | 3.23E-05 | 0.01 | 9.46E-01 | 0.00 | 1.00E+00 |
| FBgn0003444 | smo | 2301 | -0.63 | 3.91E-07 | -0.10 | 2.03E-01 | 0.00 | 1.00E+00 |
| FBgn0036282 | Smyd4-2 | 19 | 1.33 | 8.38E-04 | 1.15 | 6.49E-03 | 0.00 | 1.00E+00 |
| FBgn0086129 | snama | 1490 | -0.65 | 8.22E-09 | 0.02 | 8.84E-01 | 0.01 | 1.00E+00 |
| FBgn0011288 | Snap25 | 98 | -0.08 | 7.87E-01 | 4.02 | 5.30E-61 | -0.02 | 8.64E-01 |
| FBgn0030026 | sni | 488 | 0.71 | 2.21E-08 | 0.07 | 3.85E-01 | 0.00 | 1.00E+00 |
| FBgn0083027 | snoRNA:Psi18S-531 | 228 | 0.76 | 7.84E-06 | 0.01 | 9.54E-01 | -0.99 | 6.07E-15 |
| FBgn0065099 | snRNA:7SK | 74 | 0.77 | 8.34E-03 | 0.12 | 3.03E-01 | -0.01 | 1.00E+00 |
| FBgn0041721 | snRNA:U12 | 72 | 0.99 | 6.65E-04 | 0.25 | 9.48E-02 | 0.02 | 9.19E-01 |
| FBgn0052758 | Snx27 | 1056 | -0.64 | 4.13E-06 | 0.00 | 9.90E-01 | 0.00 | 1.00E+00 |
| FBgn0003462 | Sod1 | 4607 | 0.82 | 6.89E-09 | -0.01 | 9.62E-01 | -0.02 | 7.72E-01 |
| FBgn0033631 | Sod3 | 3675 | 0.63 | 1.89E-07 | -0.08 | 3.00E-01 | -0.01 | 1.00E+00 |
| FBgn0042630 | Sox21b | 15 | 0.01 | 9.47E-01 | -0.84 | 2.59E-02 | 0.01 | 1.00E+00 |
| FBgn0020378 | Sp1 | 380 | 0.20 | 4.39E-01 | -2.11 | 5.44E-08 | 0.01 | 1.00E+00 |
| FBgn0035710 | SP1173 | 2433 | 0.10 | 5.98E-01 | 0.11 | 1.53E-01 | 0.66 | 3.68E-07 |
| FBgn0026562 | SPARC | 1433 | 0.81 | 2.04E-06 | -0.34 | 1.04E-02 | 0.00 | 1.00E+00 |
| FBgn0040623 | Spase12 | 1070 | 0.90 | 4.41E-07 | 0.02 | 8.81E-01 | -0.01 | 1.00E+00 |
| FBgn0039172 | Spase22-23 | 2345 | 0.62 | 4.08E-07 | 0.01 | 9.32E-01 | -0.01 | 1.00E+00 |
| FBgn0037025 | Spc105R | 1970 | -0.64 | 3.39E-06 | 0.00 | 9.82E-01 | 0.02 | 6.68E-01 |
| FBgn0031549 | Spindly | 889 | -0.77 | 1.08E-11 | -0.58 | 2.41E-11 | -0.47 | 5.60E-07 |
| FBgn0003483 | spn-E | 262 | 0.05 | 8.37E-01 | 0.95 | 2.55E-08 | 0.01 | 1.00E+00 |
| FBgn0039795 | Spn100A | 2226 | 0.21 | 4.08E-01 | -0.60 | 2.38E-02 | -0.01 | 1.00E+00 |
| FBgn0028990 | Spn27A | 3624 | 0.72 | 4.88E-15 | -0.12 | 5.13E-02 | 0.00 | 1.00E+00 |
| FBgn0033115 | Spn42De | 140 | 1.10 | 3.10E-06 | 0.08 | 3.73E-01 | 0.00 | 1.00E+00 |
| FBgn0024294 | Spn43Aa | 3847 | 0.81 | 1.21E-04 | -0.01 | 9.37E-01 | -0.02 | 8.73E-01 |

| | | | | | | | | |
|---|---|---|---|---|---|---|---|---|
| FBgn0033574 | Spn47C | 40 | -2.01 | 4.50E-05 | 2.30 | 2.05E-07 | 0.00 | 1.00E+00 |
| FBgn0003486 | spo | 14 | -0.61 | 1.57E-02 | 0.01 | 9.14E-01 | 0.00 | 1.00E+00 |
| FBgn0263987 | spoon | 4782 | -0.94 | 1.55E-08 | -0.01 | 9.13E-01 | 0.00 | 1.00E+00 |
| FBgn0031260 | Spp | 2973 | 0.59 | 1.42E-06 | -0.01 | 9.49E-01 | 0.00 | 1.00E+00 |
| FBgn0032362 | spz4 | 38 | -1.26 | 9.87E-05 | -0.02 | 8.49E-01 | 0.00 | 1.00E+00 |
| FBgn0263396 | sqd | 24176 | -1.21 | 2.03E-15 | 0.01 | 9.25E-01 | 0.00 | 1.00E+00 |
| FBgn0037248 | srl | 2653 | -0.64 | 2.17E-06 | -0.05 | 5.95E-01 | 0.00 | 1.00E+00 |
| FBgn0011481 | Ssdp | 3514 | -1.21 | 3.35E-12 | -0.02 | 8.47E-01 | 0.00 | 1.00E+00 |
| FBgn0037665 | St2 | 912 | 0.77 | 6.07E-04 | -0.05 | 5.42E-01 | -0.09 | 2.64E-01 |
| FBgn0265052 | St3 | 62 | 0.34 | 1.70E-01 | 1.40 | 5.50E-08 | 1.21 | 3.44E-06 |
| FBgn0003517 | sta | 62664 | 1.36 | 1.84E-26 | -0.14 | 8.33E-02 | 0.00 | 1.00E+00 |
| FBgn0086779 | step | 1533 | -0.70 | 1.73E-06 | 0.05 | 5.62E-01 | 0.00 | 1.00E+00 |
| FBgn0046692 | Stlk | 960 | -0.69 | 1.41E-04 | 0.03 | 7.68E-01 | 0.00 | 1.00E+00 |
| FBgn0020299 | stumps | 1216 | -0.16 | 1.84E-01 | -0.61 | 1.14E-13 | -0.01 | 1.00E+00 |
| FBgn0086708 | stv | 2256 | 0.16 | 5.24E-01 | -1.48 | 4.91E-10 | 0.01 | 1.00E+00 |
| FBgn0014388 | sty | 1582 | -0.70 | 7.56E-06 | 0.04 | 6.60E-01 | 0.00 | 1.00E+00 |
| FBgn0003567 | su(Hw) | 2513 | -0.63 | 1.09E-08 | -0.01 | 9.69E-01 | 0.00 | 1.00E+00 |
| FBgn0014391 | sun | 1616 | 1.01 | 5.14E-09 | 0.01 | 9.64E-01 | -0.01 | 1.00E+00 |
| FBgn0028675 | Sur | 11 | -0.09 | 6.53E-01 | 0.58 | 4.64E-02 | 0.00 | 1.00E+00 |
| FBgn0261403 | sxc | 3212 | -0.69 | 8.92E-08 | -0.05 | 6.02E-01 | 0.00 | 1.00E+00 |
| FBgn0038826 | Syp | 23681 | -1.10 | 4.58E-08 | -0.03 | 7.82E-01 | 0.00 | 1.00E+00 |
| FBgn0028400 | Syt4 | 264 | 7.49 | 1.96E-06 | 4.89 | 1.96E-08 | 6.07 | 2.24E-12 |
| FBgn0028398 | Taf10 | 549 | 0.66 | 1.19E-04 | -0.01 | 9.18E-01 | 0.00 | 1.00E+00 |
| FBgn0004406 | tam | 490 | 0.29 | 8.95E-02 | 0.72 | 1.51E-09 | 0.01 | 8.66E-01 |
| FBgn0021795 | Tapdelta | 3819 | 0.58 | 3.60E-05 | -0.02 | 8.76E-01 | 0.00 | 1.00E+00 |
| FBgn0034451 | TBCB | 731 | 0.19 | 2.42E-01 | 0.68 | 1.75E-10 | 0.58 | 3.63E-07 |
| FBgn0285892 | tea | 768 | -0.63 | 4.20E-05 | -0.03 | 7.57E-01 | 0.00 | 1.00E+00 |
| FBgn0261953 | TfAP-2 | 180 | 0.34 | 1.52E-01 | -2.03 | 5.20E-06 | 0.01 | 1.00E+00 |
| FBgn0013347 | TfIIA-S | 1676 | 0.94 | 1.18E-09 | 0.02 | 8.85E-01 | 0.00 | 1.00E+00 |
| FBgn0026869 | Thd1 | 3729 | -0.71 | 8.71E-05 | -0.03 | 7.76E-01 | 0.01 | 1.00E+00 |
| FBgn0261560 | Thor | 335 | 1.02 | 1.07E-03 | -0.09 | 4.00E-01 | 0.00 | 1.00E+00 |
| FBgn0032988 | Tif-IA | 2210 | -0.65 | 1.22E-11 | -0.06 | 4.87E-01 | 0.00 | 1.00E+00 |
| FBgn0027359 | Tim8 | 995 | 0.89 | 3.09E-07 | 0.03 | 7.23E-01 | 0.00 | 1.00E+00 |
| FBgn0025879 | Timp | 183 | 1.31 | 4.85E-08 | 0.00 | 9.93E-01 | 0.00 | 1.00E+00 |
| FBgn0004841 | TkR86C | 16 | 2.44 | 5.62E-04 | 6.69 | 3.31E-07 | 6.67 | 9.56E-07 |
| FBgn0003721 | Tm1 | 10585 | -0.87 | 8.91E-13 | -0.01 | 9.30E-01 | 0.00 | 1.00E+00 |
| FBgn0267796 | Tmc | 187 | -1.09 | 6.16E-06 | -0.01 | 9.30E-01 | -0.01 | 9.67E-01 |
| FBgn0026160 | tna | 4864 | -0.86 | 1.63E-08 | -0.01 | 9.53E-01 | 0.00 | 1.00E+00 |
| FBgn0033357 | Tom7 | 1314 | 1.19 | 2.26E-12 | 0.04 | 6.63E-01 | 0.00 | 1.00E+00 |
| FBgn0037751 | topi | 17 | 0.16 | 2.73E-01 | 3.33 | 7.94E-09 | 0.01 | 1.00E+00 |
| FBgn0086355 | Tpi | 3109 | 0.75 | 5.13E-06 | 0.06 | 4.99E-01 | 0.00 | 1.00E+00 |
| FBgn0031692 | TpnC25D | 30 | -0.49 | 4.92E-02 | -0.68 | 4.79E-02 | -0.05 | 5.00E-01 |
| FBgn0086674 | Tpst | 715 | -1.40 | 5.38E-11 | 0.03 | 7.58E-01 | 0.00 | 1.00E+00 |
| FBgn0030748 | Traf-like | 115 | -0.73 | 5.07E-03 | -0.08 | 4.80E-01 | 0.00 | 1.00E+00 |
| FBgn0026319 | Traf4 | 481 | 0.24 | 1.13E-01 | 0.76 | 4.59E-17 | 0.00 | 1.00E+00 |
| FBgn0261793 | Trf2 | 2420 | -0.99 | 3.98E-07 | 0.03 | 7.55E-01 | 0.00 | 1.00E+00 |
| FBgn0013263 | Trl | 369 | -1.26 | 3.45E-08 | -0.01 | 9.05E-01 | 0.00 | 1.00E+00 |
| FBgn0050343 | Tsen15 | 167 | -1.57 | 1.05E-14 | 0.02 | 8.94E-01 | 0.00 | 1.00E+00 |
| FBgn0003866 | tsh | 5545 | -0.66 | 4.17E-20 | -0.16 | 2.69E-03 | 0.00 | 1.00E+00 |
| FBgn0031850 | Tsp | 2665 | 0.69 | 2.98E-10 | -0.34 | 3.00E-04 | 0.00 | 1.00E+00 |
| FBgn0032943 | Tsp39D | 1560 | -0.58 | 8.20E-05 | 0.01 | 9.37E-01 | 0.00 | 1.00E+00 |
| FBgn0029507 | Tsp42Ed | 56 | 0.33 | 1.26E-01 | 1.57 | 7.02E-04 | -0.02 | 8.30E-01 |
| FBgn0033130 | Tsp42Ei | 87 | 0.57 | 2.08E-02 | -0.61 | 1.22E-02 | -0.02 | 7.84E-01 |
| FBgn0043550 | Tsp68C | 24 | 0.92 | 4.23E-03 | 3.43 | 4.52E-12 | 0.04 | 5.83E-01 |
| FBgn0027865 | Tsp96F | 1091 | -0.73 | 1.13E-04 | 0.00 | 9.93E-01 | 0.00 | 1.00E+00 |
| FBgn0032744 | Ttc19 | 220 | -0.65 | 9.39E-04 | 1.15 | 1.18E-14 | 0.01 | 1.00E+00 |
| FBgn0051108 | TTLL5 | 618 | -0.89 | 5.37E-10 | 0.00 | 9.85E-01 | 0.00 | 1.00E+00 |
| FBgn0052364 | tut | 126 | -0.50 | 3.44E-02 | -0.04 | 6.75E-01 | -3.19 | 3.46E-25 |
| FBgn0039434 | TwdlM | 16 | 0.92 | 5.02E-03 | 0.34 | 1.02E-01 | 2.59 | 7.18E-05 |
| FBgn0029170 | TwdlT | 119 | 0.43 | 7.55E-02 | -1.52 | 1.34E-06 | -0.11 | 2.46E-01 |
| FBgn0003900 | twi | 636 | 0.03 | 8.92E-01 | -0.69 | 5.16E-11 | -0.12 | 1.00E-01 |
| FBgn0262801 | twr | 2970 | 0.60 | 6.91E-05 | -0.03 | 8.10E-01 | -0.01 | 1.00E+00 |
| FBgn0034636 | twz | 761 | 0.58 | 1.40E-03 | -0.01 | 9.54E-01 | 0.03 | 6.00E-01 |
| FBgn0029996 | UbcE2H | 1978 | -0.73 | 9.00E-09 | 0.15 | 6.58E-02 | 0.01 | 1.00E+00 |
| FBgn0026076 | UBL3 | 1245 | -0.79 | 6.80E-08 | -0.02 | 8.24E-01 | 0.00 | 1.00E+00 |
| FBgn0003944 | Ubx | 347 | -0.41 | 9.19E-02 | -1.48 | 4.15E-08 | 0.00 | 1.00E+00 |
| FBgn0262124 | uex | 1998 | -0.87 | 4.48E-09 | 0.01 | 9.32E-01 | 0.00 | 1.00E+00 |
| FBgn0040259 | Ugt302C1 | 1204 | -0.23 | 8.13E-02 | 0.70 | 2.85E-15 | 0.00 | 1.00E+00 |
| FBgn0040251 | Ugt302K1 | 820 | -1.93 | 1.49E-26 | 2.29 | 4.84E-76 | 0.00 | 1.00E+00 |
| FBgn0040091 | Ugt317A1 | 411 | 0.43 | 6.49E-03 | -1.04 | 4.52E-19 | -1.12 | 4.52E-23 |
| FBgn0045800 | Uhg1 | 543 | -0.62 | 8.92E-04 | 0.04 | 6.63E-01 | 0.00 | 1.00E+00 |
| FBgn0083124 | Uhg4 | 570 | -0.63 | 2.41E-04 | -0.02 | 8.43E-01 | 0.00 | 1.00E+00 |
| FBgn0025549 | unc-119 | 1224 | -0.71 | 1.70E-13 | -0.09 | 1.90E-01 | 0.00 | 1.00E+00 |
| FBgn0025726 | unc-13 | 8539 | -0.16 | 3.82E-01 | 3.08 | 1.68E-117 | 0.01 | 9.70E-01 |
| FBgn0024184 | unc-4 | 56 | 0.21 | 1.73E-01 | -1.78 | 2.44E-04 | 0.01 | 1.00E+00 |
| FBgn0263352 | Unr | 3986 | -1.51 | 8.25E-12 | -0.01 | 9.12E-01 | 0.00 | 1.00E+00 |
| FBgn0030904 | upd2 | 26 | 0.21 | 1.41E-01 | 2.98 | 6.12E-04 | 0.00 | 1.00E+00 |
| FBgn0053542 | upd3 | 34 | 0.01 | 9.69E-01 | 2.77 | 2.35E-07 | 0.01 | 1.00E+00 |
| FBgn0034245 | UQCR-6.4 | 522 | 0.83 | 2.12E-05 | 0.02 | 8.90E-01 | 0.00 | 1.00E+00 |

| | | | | | | | | |
|---|---|---|---|---|---|---|---|---|
| FBgn0038271 | UQCR-C1 | 4523 | 0.84 | 1.29E-13 | 0.03 | 7.62E-01 | 0.00 | 1.00E+00 |
| FBgn0036728 | UQCR-Q | 1447 | 1.88 | 3.34E-40 | 0.00 | 9.79E-01 | 0.00 | 1.00E+00 |
| FBgn0033428 | Urod | 1510 | 0.64 | 3.48E-06 | 0.02 | 8.25E-01 | 0.00 | 1.00E+00 |
| FBgn0260749 | Utx | 1479 | -0.75 | 1.33E-17 | -0.06 | 4.40E-01 | 0.00 | 1.00E+00 |
| FBgn0050101 | Vajk4 | 36 | 1.18 | 5.97E-04 | 0.14 | 2.34E-01 | 0.00 | 1.00E+00 |
| FBgn0035942 | ValRS-m | 232 | 0.60 | 4.18E-04 | -0.01 | 9.18E-01 | 0.00 | 1.00E+00 |
| FBgn0029687 | Vap33 | 7747 | -0.78 | 2.49E-07 | 0.01 | 9.16E-01 | 0.00 | 1.00E+00 |
| FBgn0053200 | VepD | 48 | -1.91 | 1.02E-06 | 0.11 | 3.01E-01 | 0.00 | 1.00E+00 |
| FBgn0262524 | ver | 296 | -0.22 | 2.25E-01 | 2.11 | 4.76E-65 | 0.00 | 1.00E+00 |
| FBgn0261341 | verm | 9505 | 0.62 | 9.38E-04 | -0.13 | 1.70E-01 | 0.00 | 1.00E+00 |
| FBgn0033911 | VGAT | 134 | -0.09 | 7.49E-01 | 0.64 | 1.29E-03 | -0.63 | 6.34E-03 |
| FBgn0267975 | vib | 1469 | -1.04 | 1.87E-26 | 0.04 | 6.69E-01 | -0.01 | 1.00E+00 |
| FBgn0024183 | vig | 3544 | -0.78 | 1.04E-08 | 0.01 | 9.06E-01 | 0.00 | 1.00E+00 |
| FBgn0259978 | vlc | 2238 | -0.62 | 3.56E-08 | -0.03 | 7.54E-01 | 0.00 | 1.00E+00 |
| FBgn0052350 | Vps11 | 1027 | -0.65 | 2.49E-03 | 0.00 | 9.72E-01 | 0.00 | 1.00E+00 |
| FBgn0260987 | vtd | 2064 | -0.95 | 1.33E-21 | -0.06 | 4.50E-01 | -0.01 | 1.00E+00 |
| FBgn0266848 | wap | 3147 | -0.66 | 1.92E-08 | -0.03 | 7.45E-01 | 0.00 | 1.00E+00 |
| FBgn0027492 | wdb | 5970 | -0.84 | 1.30E-11 | 0.01 | 9.22E-01 | 0.00 | 1.00E+00 |
| FBgn0027499 | wde | 2735 | -0.89 | 1.99E-07 | 0.02 | 8.39E-01 | 0.00 | 1.00E+00 |
| FBgn0011739 | wts | 1411 | -0.66 | 9.62E-07 | 0.02 | 8.38E-01 | 0.00 | 1.00E+00 |
| FBgn0261113 | Xrp1 | 10923 | -0.55 | 6.59E-06 | 1.08 | 1.69E-26 | 0.11 | 1.21E-01 |
| FBgn0041711 | yellow-e | 215 | 0.87 | 5.43E-05 | -0.03 | 7.39E-01 | 0.00 | 1.00E+00 |
| FBgn0041710 | yellow-f | 83 | -0.23 | 3.71E-01 | -0.62 | 3.15E-02 | -0.05 | 5.27E-01 |
| FBgn0039896 | yellow-h | 56 | 0.51 | 4.40E-02 | 2.46 | 1.25E-10 | -0.01 | 1.00E+00 |
| FBgn0040060 | yip7 | 41 | -0.04 | 6.60E-01 | -0.97 | 3.20E-02 | -0.02 | 9.62E-01 |
| FBgn0039261 | Ythdf | 2954 | -0.66 | 9.40E-09 | -0.04 | 6.25E-01 | 0.00 | 1.00E+00 |
| FBgn0004606 | zfh1 | 688 | -0.13 | 5.56E-01 | -0.63 | 2.67E-04 | 0.00 | 1.00E+00 |
| FBgn0266709 | Zmynd10 | 117 | -1.05 | 1.89E-06 | -0.13 | 2.22E-01 | 0.00 | 1.00E+00 |
| FBgn0011642 | Zyx | 2299 | -1.25 | 1.45E-35 | 0.00 | 9.77E-01 | 0.00 | 1.00E+00 |

**Supplementary Table 4: Ontology of genes deregulated in *corto*$^{L1}$*/corto*$^{420}$ *and uL11*$^{K3A}$ mutants**

CC: Cellular Component; BP: Biological Process; MF: Molecular Function

| *Expression in* corto$^{L1}$/corto$^{420}$ | | *GO term* | *Count* | *p-value* | *Benjamini* |
|---|---|---|---|---|---|
| up-regulated | BP-GO:0002181 | cytoplasmic translation | 66 | 6.90E-72 | 4.20E-69 |
| up-regulated | CC-GO:0005840 | ribosome | 61 | 1.00E-61 | 2.00E-59 |
| up-regulated | MF-GO:0003735 | structural constituent of ribosome | 82 | 5.40E-47 | 1.70E-44 |
| up-regulated | CC-GO:0022625 | cytosolic large ribosomal subunit | 38 | 3.90E-39 | 3.90E-37 |
| up-regulated | CC-GO:0005840 | cytosolic small ribosomal subunit | 28 | 4.10E-28 | 2.70E-26 |
| up-regulated | CC-GO:0005840 | cytosolic ribosome | 41 | 2.10E-23 | 1.00E-21 |
| up-regulated | BP-GO:0006412 | translation | 74 | 2.80E-22 | 8.50E-20 |
| up-regulated | CC-GO:0005840 | mitochondrial respiratory chain complex I | 15 | 6.40E-10 | 1.80E-08 |
| up-regulated | MF-GO:0003954 | NADH dehydrogenase activity | 10 | 1.50E-06 | 1.60E-04 |
| up-regulated | BP-GO:0002181 | mitochondrial electron transport, NADH to ubiquinone | 9 | 2.80E-05 | 2.90E-03 |
| up-regulated | MF-GO:0003954 | NADH dehydrogenase (ubiquinone) activity | 8 | 4.10E-04 | 1.80E-02 |
| up-regulated | CC-GO:0005751 | mitochondrial respiratory chain complex IV | 6 | 1.10E-04 | 1.70E-03 |
| up-regulated | MF-GO:0004129 | cytochrome-c oxidase activity | 7 | 1.80E-04 | 1.20E-02 |
| up-regulated | BP-GO:0006123 | mitochondrial electron transport, cytochrome c to oxygen | 5 | 5.40E-04 | 3.70E-02 |
| up-regulated | CC-GO:0005750 | mitochondrial respiratory chain complex III | 5 | 5.10E-04 | 7.20E-03 |
| up-regulated | MF-GO:0008121 | ubiquinol-cytochrome-c reductase activity | 5 | 7.10E-04 | 2.50E-02 |
| up-regulated | BP-GO:0006122 | mitochondrial electron transport, ubiquinol to cytochrome c | 5 | 1.80E-03 | 9.00E-02 |
| up-regulated | BP-GO:0006465 | signal peptide processing | 5 | 8.40E-05 | 7.40E-03 |
| up-regulated | CC-GO:0005787 | signal peptidase complex | 3 | 7.90E-03 | 9.20E-02 |
| up-regulated | BP-GO:1902600 | proton transport | 7 | 2.00E-04 | 1.50E-02 |
| up-regulated | MF-GO:0046933 | proton-transporting ATP synthase activity, rotational mechanism | 6 | 3.80E-04 | 1.80E-02 |
| up-regulated | BP-GO:0040003 | chitin-based cuticle development | 15 | 1.10E-03 | 6.70E-02 |
| up-regulated | MF-GO:0009055 | electron carrier activity | 8 | 2.20E-03 | 7.00E-02 |
| down-regulated | BP-GO:0006465 | protein phosphorylation | 30 | 7.60E-09 | 9.30E-06 |
| down-regulated | MF-GO:0004674 | protein serine/threonine kinase activity | 23 | 1.20E-08 | 2.10E-06 |
| down-regulated | MF-GO:0004672 | protein kinase activity | 19 | 4.00E-06 | 4.60E-04 |
| down-regulated | MF-GO:0005524 | ATP binding | 45 | 2.30E-04 | 1.30E-02 |
| down-regulated | BP-GO:0045475 | locomotor rhythm | 9 | 9.60E-04 | 6.20E-02 |
| down-regulated | CC-GO:0005834 | heterotrimeric G-protein complex | 5 | 7.10E-04 | 3.00E-02 |
| down-regulated | MF-GO:0004871 | signal transducer activity | 7 | 1.40E-03 | 5.30E-02 |
| down-regulated | MF-GO:0031683 | G-protein beta/gamma-subunit complex binding | 4 | 1.90E-03 | 5.50E-02 |
| down-regulated | BP-GO:0030307 | positive regulation of cell growth | 7 | 2.70E-04 | 2.40E-02 |

| *Expression in* uL11$^{K3A}$ | | *GO term* | *Count* | *p-value* | *Benjamini* |
|---|---|---|---|---|---|
| up-regulated | BP~GO:0006749 | glutathione metabolic process | 9 | 3.10E-07 | 1.30E-04 |
| up-regulated | MF~GO:0004364 | glutathione transferase activity | 8 | 5.00E-06 | 8.90E-04 |
| up-regulated | MF~GO:0004602 | glutathione peroxidase activity | 7 | 8.10E-06 | 8.90E-04 |
| up-regulated | BP~GO:0000723 | telomere maintenance | 6 | 1.30E-05 | 2.60E-03 |
| up-regulated | BP~GO:0006310 | DNA recombination | 5 | 6.70E-04 | 7.60E-02 |
| down-regulated | MF~GO:0043565 | sequence-specific DNA binding | 13 | 2.90E-05 | 4.10E-03 |
| down-regulated | BP~GO:0006355 | regulation of transcription, DNA-templated | 14 | 1.20E-04 | 9.30E-03 |
| down-regulated | BP~GO:0006351 | transcription, DNA-templated | 13 | 2.60E-04 | 1.50E-02 |
| down-regulated | MF~GO:0003700 | transcription factor activity, sequence-specific DNA binding | 11 | 8.50E-04 | 4.40E-02 |
| down-regulated | MF~GO:0004035 | alkaline phosphatase activity | 4 | 9.30E-04 | 4.40E-02 |

**Supplementary Table 5: Enrichment for Mad sites in the *cis*-regulatory sequences of genes deregulated in *uL11*$^{K3Y}$ mutants.**

21 nucleotide-long motifs shared by 39 of the 84 *uL11*$^{K3Y}$ deregulated genes (E-value: 2.1E-23).

dnv: does not vary

| Gene Id | Gene symbol | Expression in *uL11*$^{K3Y}$ versus *w*$^{1118}$ | Expression in *uL11*$^{K3A}$ versus *w*$^{1118}$ | Strand | Distance from the TSS | *p*-value | | Sites | |
|---|---|---|---|---|---|---|---|---|---|
| *FBgn0267160* | *asRNA:CR45600* | up | up | - | -1367 | 1.24E-06 | CAGCGAAGGA | GACGCCGTCCAAGCCGTCACC | ACTTTGTGAG |
| *FBgn0034972* | *CG10339* | up | up | - | -1148 | 2.21E-07 | CATTACCACT | GCCACCGCCGCCGATGTGGCC | GTTCTCTTCG |
| *FBgn0036589* | *CG13067* | up | dnv | + | -1578 | 3.18E-10 | TGCCTACACC | GCTCCAGTTGCCGCCGCTGCC | TATACCGCTC |
| *FBgn0040658* | *CG13516* | down | down | + | 621 | 1.65E-06 | GGCTCCAGCG | GTTTCCGCACCAGCAATCGCC | CCAGCACGAC |
| *FBgn0037127* | *CG14566* | up | dnv | + | 257 | 5.47E-09 | CTTCGACGAG | GCTGCTGCTCCCTCGGCCGCC | GGGCCCACAC |
| *FBgn0035409* | *CG14963* | up | dnv | - | -1497 | 6.83E-08 | AGTTTCCGCT | GCTGCCTTAGCAGCCGCTCGC | TTCAGTTCAT |
| *FBgn0033777* | *CG17574* | down | up | - | -1371 | 1.19E-07 | CAGCAAGAGC | GGCGCCGGCGGAGCAGCCGGC | TCTTCAGCCG |
| *FBgn0017448* | *CG2187* | up | up | - | -179 | 1.54E-06 | ATATTCATCG | GTTGCAGTTCAAGCTATTGCC | GATAACCCAA |
| *FBgn0034931* | *CG2812* | up | up | - | 836 | 1.35E-08 | AGATCTTGCA | GTCCCCGGCGCCGCCGCTCGC | CAGCAGCCGG |
| *FBgn0052354* | *CG32354* | up | up | + | -51 | 4.64E-08 | GCTCTGCTCT | GCCGTCGCCGACTGCGCTGCC | TTTCACCCAT |
| *FBgn0052532* | *CG32532* | up | up | + | 655 | 1.30E-07 | GCAGGACAGC | GCTGCTTCTGCCGGCGTCGAC | GCGTCCTGCT |
| *FBgn0085276* | *CG34247* | down | dnv | - | -1572 | 3.44E-08 | TCAAGACTCG | GCCGCCAGAGCACCTGTCGCC | CGAAGCTCAA |
| *FBgn0085411* | *CG34382* | down | down | - | 374 | 3.64E-07 | GACTACGATG | GTCGCATCTGCTGCATCTGCA | TCGAGCATTG |
| *FBgn0085452* | *CG34423* | down | down | - | 265 | 1.57E-12 | GGATGGATCC | GCCGCCGCCGCCGCCGTTGCC | GGCTCCACTG |
| *FBgn0260768* | *CG42566* | down | down | + | -1898 | 1.09E-07 | TGATTCTCCA | GCTGCTGGCCCTTCATCTGCC | TTCCCTACTT |
| *FBgn0038784* | *CG4362* | down | down | + | 57 | 2.03E-06 | GCATGACCTT | GCTGCTGGCCATCGGGCTGCA | GCAGATCGAC |
| *FBgn0036587* | *CG4950* | down | dnv | - | -1566 | 3.18E-10 | TGCCTACACC | GCTCCAGTTGCCGCCGCTGCC | TATACCGCTC |
| *FBgn0039430* | *CG5455* | down | down | - | -829 | 1.70E-07 | AGGACGAGGA | GCTGGAGGAGCTGCTGTCCCC | GGAACTGGTG |
| *FBgn0036121* | *CG6310* | down | down | + | -670 | 1.67E-08 | ATATGGACGA | GCTGCCAGCGCAGCTTTCGCC | AAAGCTTTTG |
| *FBgn0028533* | *CG7953* | down | down | - | 105 | 2.61E-07 | TGATCCGCCA | GCTCCAACAGACGCTGCTCCA | CCTCGTTGGG |
| *FBgn0038405* | *CG8927* | up | up | - | -1482 | 8.24E-08 | ACTTGGATCC | GGTGCATCTGCCTCCACTGCC | TTCGACTCTA |
| *FBgn0036109* | *Cpr67Fa2* | down | dnv | + | -1471 | 8.61E-07 | CGCTATCCTG | GCCTGCGCCTACGGTGCCGCC | ACCTACAACC |
| *FBgn0262636* | *dati* | up | up | - | -1275 | 6.86E-07 | GGTAACGTTA | GCCGCCACCTCTGCTGCTCAA | TCGATTTTGG |
| *FBgn0062928* | *hpRNA:CR33940* | down | dnv | + | -834 | 2.61E-07 | GTGGGCGGTA | GGTGGTGGTGCATCTGCTGCA | CTCGTTGGCA |
| *FBgn0286204* | *ich* | up | up | - | -1332 | 1.33E-06 | GGTTGATATG | GGTACTTTCCCCGCCGCCTCC | GCTGTATCCG |
| *FBgn0039459* | *IntS12* | up | dnv | + | 108 | 1.20E-11 | CGCAAATATA | GCCGCCGCTGCTGCAGCTGCC | CAAGAAGTGG |
| *FBgn0034098* | *krimp* | up | dnv | + | -747 | 4.85E-09 | TGGTGGATCT | GTCGCCGGTGACCCTGCTGCC | GTCAGTGAAG |
| *FBgn0259993* | *lncRNA:CR42491* | down | dnv | + | -1111 | 1.73E-11 | GAAGAAGGCC | GCTCCCGCAGCCGCCGCCGCC | AAGGGCAAGG |
| *FBgn0039851* | *mey* | up | up | + | -80 | 3.40E-09 | AAGAGAGTTA | GTCGCCGCTGCCTCTGCCGGC | GTCGCTTGGG |
| *FBgn0035976* | *PGRP-LC* | down | down | - | -1328 | 1.09E-07 | CATACGTGTT | GGCCCAGTCCCAGCCGCGGCC | AACGTAGATG |
| *FBgn0038966* | *pinta* | up | up | - | -839 | 2.38E-10 | AACTGTTGCT | GCTGCTGCTGCTGCTGCTGCA | GATATTGCTG |
| *FBgn0020907* | *Scp2* | up | dnv | - | -158 | 5.02E-07 | CGTCTCGGTC | GACTCTGCCGACGCCTCTGCA | CTCGTCATGG |
| *FBgn0083027* | *snoRNA:Psi18S-531* | down | dnv | + | -1627 | 1.73E-11 | GAAGAAGGCC | GCTCCCGCAGCCGCCGCCGCC | AAGGGCAAGG |
| *FBgn0265052* | *St3* | up | up | - | -860 | 9.05E-08 | TCGAAGGAGA | GGTGTCGCTGCATCTGCTGCA | TCTGCTCCTG |
| *FBgn0034451* | *TBCB* | up | up | - | -1977 | 2.84E-07 | TCAAACAAAA | GCTGCTGGTCCTCCAGCTCCA | CAGCATATTG |
| *FBgn0004841* | *TkR86C* | up | up | - | -9 | 6.13E-09 | TGTCCTTAGC | GCTGCAAGTGCCGCCGCCGAC | TGAATGCTTT |
| *FBgn0052364* | *tut* | down | dnv | - | -622 | 2.67E-09 | TCAAGTGCCT | GCTGCCGCTGCTGCTGCTTCA | TTTTCCTAAT |
| *FBgn0039434* | *TwdlM* | up | dnv | - | 313 | 3.69E-10 | CTCCGATGCT | GCCGCCTCCGATGCTGCCGCC | TCCGATGCTG |
| *FBgn0033911* | *VGAT* | down | up | + | 399 | 1.35E-08 | ACGCCTTCTT | GACACCACCGCAGCCGCCGCC | ATCAGGCCAA |
